# Size reductions and genomic changes associated with harvesting within two generations in wild walleye populations

**DOI:** 10.1101/787374

**Authors:** Ella Bowles, Kia Marin, Stephanie Mogensen, Pamela Macleod, Dylan J. Fraser

## Abstract

The extent and rate of harvest-induced genetic changes in natural populations may impact population productivity, recovery and persistence. While there is substantial evidence for phenotypic changes in harvested fishes, knowledge of genetic change in the wild remains limited, as phenotypic and genetic data are seldom considered in tandem, and the number of generations needed for genetic changes to occur is not well understood. We quantified changes in size-at-age, sex-specific changes in body size, and genomic metrics in three harvested walleye (*Sander vitreus*) populations and a fourth reference population with low harvest levels over a 15-year period in Mistassini Lake, Quebec. We also collected Traditional Ecological Knowledge (TEK) surrounding concerns about these populations over time. Using ∼9000 SNPs, genomic metrics included changes in population structure, neutral genomic diversity, effective population size and signatures of selection. TEK revealed concerns about overall reductions in body size and number of fish caught. Smaller body size, smaller size-at-age, changing population structure (population differentiation within one river and homogenization between two others), and signatures of selection between historical and contemporary samples reflected coupled phenotypic and genomic change in the three harvested populations in both sexes, while no change occurred in the reference population. Sex-specific analyses revealed differences in both body size and genomic metrics but were inconclusive about whether one sex was disproportionately affected. Our results support that harvest-induced genetic changes can arise within 1-2.5 generations in long-lived wild fishes, demonstrating the need to investigate concerns about harvest-induced evolution quickly once they have been raised.

## Introduction

Harvesting of wild populations can affect growth, body-size, maturation and population productivity (Heino *et al*. 2013; Heino & Godø 2002; Hutchings 2005); but it can also alter genetic population structuring, reduce genetic diversity (primarily through reducing population size) and select for different genotypes that underlie phenotypic traits (the latter commonly referred to as fisheries-induced evolution, FIE) (Allendorf *et al*. 2008; Hutchings & Fraser 2008). Because harvest-induced genetic changes can affect population productivity, recovery and persistence, assessing how quickly, to what extent and under what circumstances such changes arise has become an emerging component of contemporary fisheries management (Heino *et al*. 2015; Jorgensen *et al*. 2007; Law & Grey 1989; Rowell 1993).

Many studies have shown rapid phenotypic change towards smaller body size and size-at-age in harvested fish populations, though whether such changes are plastic responses (Law 2007), genetic changes or both is a source of ongoing debate (Heino *et al*. 2015; Jorgensen *et al*. 2007; Sharpe & Hendry 2009). Much of the empirical evidence that fishing causes rapid, genetically-based phenotypic change comes from lab-based studies (e.g. within three generations (van Wijk *et al*. 2013), or two to five generations (Therkildsen *et al*. 2019; Uusi-Heikkilä *et al*. 2017)). However, lab environments can introduce unintended selection pressures (possibly body condition, growth, maturation, Uusi-Heikkilä *et al*. 2017) and may not adequately depict the actual extent or rate of harvest-induced change that wild fishes experience (Fraser *et al*. 2018). Results from the few studies that have integrated phenotypic and genetic evidence in the wild suggest that harvest-induced genetic change may occur within as little as one generation (Chebib *et al*. 2016), to four to eight (Allen *et al*. 2017), or longer (Hutchinson *et al*. 2003; Therkildsen *et al*. 2013), though these studies were based on relatively limited genetic data and/or did not consider sex. Indeed, how genetic change from fishing may differentially affect males and females is understudied in fishes, despite that in many species the sexes exhibit divergent, genetically-based life histories (Fraser *et al*. 2018), and that harvest may affect the sexes differently (Hixon *et al*. 2013; Hutchings & Rowe 2008; Philipp *et al*. 2015). Overall, there remains much to learn in nature about how fishing may drive genetic changes in the life history (e.g. body size, size-at-age, by sex), and genetic characteristics (e.g. population structure, genetic diversity and composition) of wild populations.

While Western Scientific Methods (WSM) are most often used to inform fisheries management, inclusion of Traditional Ecological Knowledge (TEK) has become an integral complement to scientific knowledge for wildlife management and community-based conservation (Berkes *et al*. 2000; Fraser *et al*. 2006; Polfus *et al*. 2016; Polfus *et al*. 2014). TEK is defined as the “cumulative body of knowledge, practice and belief, evolving by adaptive processes and handed down through generations by cultural transmission, about the relationship of living beings (including humans) with one another and with their environment” (Berkes *et al*. 2000). Importantly, TEK provides extensive location-specific knowledge, can detect changes in wildlife more quickly than WSM (Huntington 2011), and often provides increased knowledge of environmental linkages (Chapman 2007; Drew 2005).

Walleye (*Sander vitreus*) are important for commercial, sport and Indigenous subsistence fisheries across North America (Bozek *et al*. 2011; Hansen *et al*. 2015; Scott & Crossman 1979). Mistassini Lake in northern Quebec, Canada, is the province’s largest natural lake (161 km long, 2 335 km^2^, 183 m maximum depth), is in Grand Council of the Crees land, *Eeyou Istchee*, and is considered to be largely pristine (minimal mining, forestry, development; no known invasive species) (Fraser *et al*. 2006; Marin *et al*. 2017). The motivation behind this study was observations by Cree elders and fishers of reduced body size and catch rates in walleye populations in three of Mistassini Lake’s southern tributaries that are close to the community, and a desire by the community to determine if management actions were needed. We also studied a fourth river at the northeastern tip of the lake, where the population was perceived to be largely unaffected by fishing until very recently (∼2015, TEK, see methods). Subsistence harvest takes place on the rivers during spawning in the spring, and walleye from different rivers comprise a mixed-population fishery in the lake during the summer, both recreationally and for subsistence (Table S1). However, recreational non-Cree fishers are only permitted to fish below the 51^st^ parallel when they are without a Cree guide (Figure 1), fishing by Cree appears to be mostly in the south (Table 2), and the genetically-distinct populations that contribute most to the mixed summer fishery are those from the rivers of concern (Dupont *et al*. 2007). Documented catch by non-Cree fishers without a guide has not increased between 1997 and 2015 (Table S1), but we do not have data on direct or latent mortalities due to local fishing derbies. In addition, the human population and the number of households in Mistissini almost doubled between 1997 and 2016 (Table S1). Cumulatively, this information indicates an increase in fishing pressure in the southern rivers.

**Table 1:** Details of sample sizes for walleye caught in each tributary of Mistassini Lake for each sex and year for each analysis, as well as whether samples were caught from a boat or on shore.

| River | Year | Boat or shore fishing | Body size (2002/03 samples grouped)^ | Size-at-age | Genomic after removing aberrant individuals (only 2003 and 2015 samples used)* |
| --- | --- | --- | --- | --- | --- |
| **Chalifour** | 2002 | ‡primarily boat |  | 21(M), 11(F) |  |
|  | 2003 | ‡primarily boat | 164(M), 14(F) |  | 22(M), 8(F) |
|  | 2015 | primarily boat | 118(M), 14(F), 44(U) | 18(M), 3(F) | 39(M), 9(F), 1(U) |
|  | 2016 | primarily boat | 132(M), 29(F), 2(U) |  |  |
|  | 2017 | primarily boat | 96(M), 12(F), 9(U) | 10(M), 4(F) |  |
| **Perch** | 2002 | not available |  | 17(M), 37(F) |  |
|  | 2003 | not available | 113(M), 43(F) |  | 24(M), 24(F) |
|  | 2015 | primarily boat | 34(M), 13(F), 13(U) | 3(M), 3(F) | 31(M), 11(F) |
|  | 2016 | primarily shore | 78(M), 26(F), 9(U) |  |  |
|  | 2017 | primarily shore | 12(M) |  |  |
| **Icon** | 2002 | not available |  | 27(M), 34(F) |  |
|  | 2003 | not available | 77(M), 43(F) |  | 23(M), 24(F) |
|  | 2015 | primarily shore | 106(M), 8(F) | 13(M), 3(F) | 37(M), 7(F) |
|  | 2016 | primarily shore | 156(M), 13(F), 1(U) |  |  |
|  | 2017 | primarily shore | 38(M), 1(F), 1(U) |  |  |
| **Takwa** | 2002 | ‡primarily boat |  | 28(M), 15(F) |  |
|  | 2003 | ‡primarily boat | 64(M), 81(F), 26(U) |  | 17(M), 16(F), 2(U) |
|  | 2015 | boat | 116(M), 15(F), 19(U) | 10(M), 11(F) | 20(M), 19(F) |
|  | 2016 | boat |  |  |  |
|  | 2017 | boat | 51(M), 22(F), 76(U) |  |  |

‡Takwa and Chalifour spawning grounds are only accessible by boat; thus, while we do not have record of exactly how fishing was conducted, boat can be inferred
^Individuals with unknown sex were not used in body size models, and nor were categories with any fewer than 8 observations
\*We extracted DNA from 371 walleye (historical n = 173 from 2003, contemporary n = 198 from 2015), and that Icon and Perch were analyzed as a single population

**Table 2:** TEK for 17 fishers with >25 years of walleye fishing experience on Mistassini Lake. No. is the number of respondents for that answer, and Freq is the frequency of respondents using the number of respondents for the observation as the denominator. Frequency is not given for the observation for which respondents could respond for multiple trends.

| Observation | Trend | Timeline of observation | No. | Freq |
| --- | --- | --- | --- | --- |
| Area fished | Region close to community | Current time | 16 | 0.94 |
|  | Far northeast, close to northern reference river | Current time | 9 | 0.53 |
|  | Other areas of the lake | Current time | ≤7 | 0.41 |
| Size of fish | Smaller | 5 - 25 yrs | 12 | 0.71 |
|  | Smaller in the south | 15 years | 3 | 0.18 |
|  | No change in average, but Takwa River fish are smaller | 5yrs | 1 | 0.06 |
|  | Different sizes in different seasons | within years | 1 | 0.06 |
|  | Not sure |  | 1 | 0.06 |
| Abundance of walleye | Decreasing | 5 - 25yrs | 13 | 0.76 |
|  | Fewer in the south but not in the north |  | 1 | 0.06 |
|  | No change | n/a | 1 | 0.06 |
|  | Does not know | n/a | 2 | 0.12 |
| Other concerns about health of walleye | Fishing during spawning or taking too many during spawning |  | 6 |  |
|  | Use of snares to fish in the spring |  | 3 |  |
|  | Night fishing, taking too many |  | 2 |  |
|  | Overfishing |  | 5 |  |
|  | Damage to fish after being handled |  | 1 |  |
|  | Fishing derby's causing many dead fish |  | 2 |  |
|  | Pike are after walleye eggs |  | 2 |  |
|  | Pollution in lake, dirty water |  | 2 |  |
|  | No other concerns, or they will be fine |  | 8 |  |

**Figure 1.**
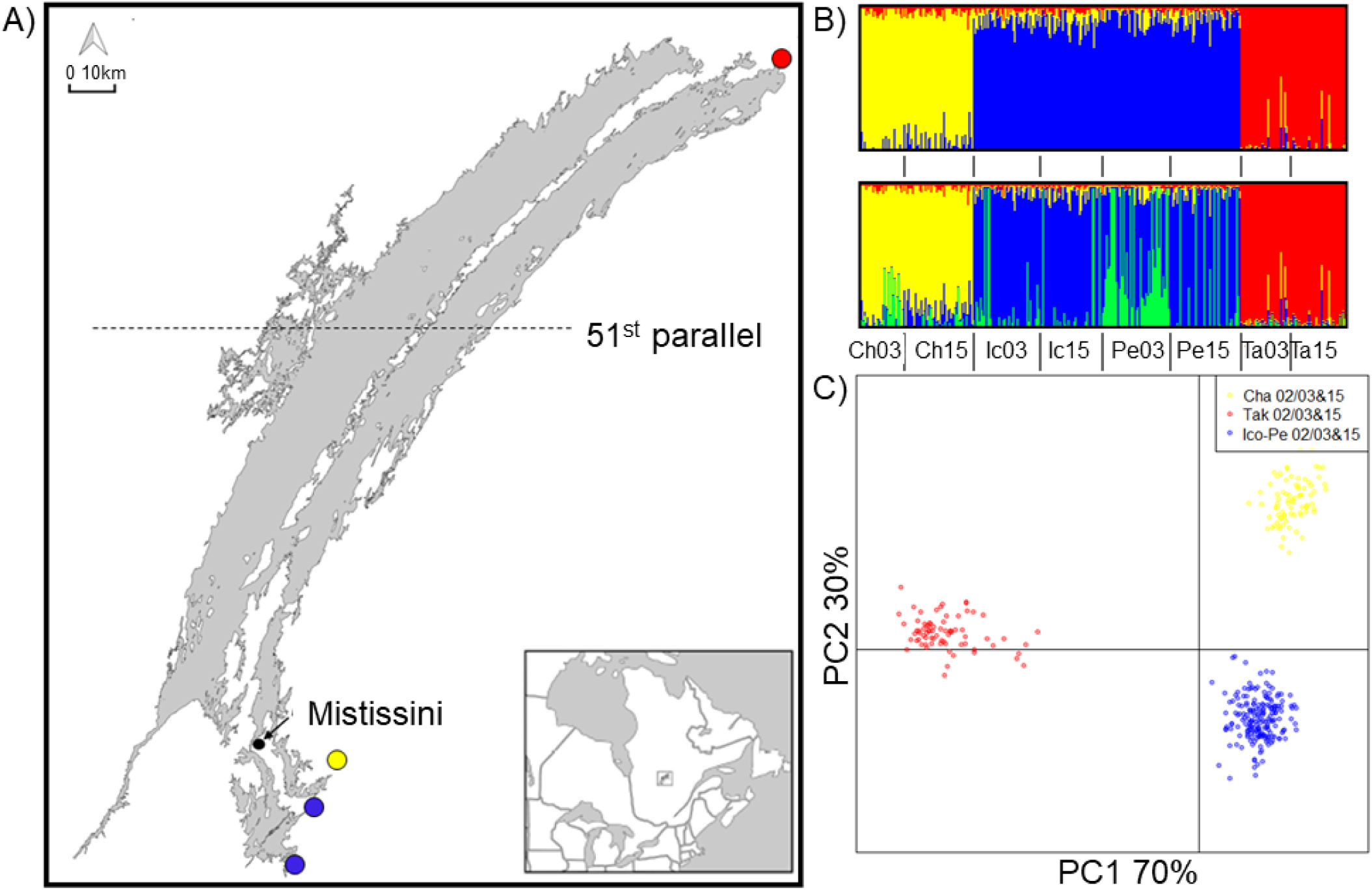
A) Map of sampling sites: red is Takwa River, blue is Icon and Perch Rivers, green is an historical genotype in Perch River, yellow is Chalifour River. B) ADMIXTURE results showing, top to bottom, K = 3 and 4). C) DAPC analysis showing k = 3 (i.e., historical and contemporary sampling years are not separated statistically by DAPC).

Using tissue samples and body size measurements collected in 2002/03 (Dupont *et al*. 2007) (“historical”) and between 2015-2017 (“contemporary”), we tested the general hypothesis that harvesting over a period of 1-2.5 generations (based on ages of spawners in the southern rivers of Mistassini Lake (supplementary data, Dupont *et al*. 2007)) was sufficient to cause coupled phenotypic and genetic changes in wild walleye populations. Specifically, we predicted that, in association with recent, increased fishing effort in Mistassini Lake, the following should be evident within the southern, harvested rivers but not in the northern river with limited harvesting, when comparing contemporary versus historic samples. 1) Reduced body size (total length and mass). 2) Reduced size-at-age. 3) Changes to population structure such as collapsing/homogenization of between-river population structure. 4) Reductions in genetic diversity and effective population size. 5) Signatures of selection, with putatively selected loci related to growth, body size and/or maturation. 6) Greater reductions in body size, size-at-age and stronger signatures of selection in females than in males, as a sexually dimorphic species with larger females than males. As one of the relatively few studies incorporating genomic and phenotypic data in wild populations to date, and the first to show rapid genetic change in a long-lived species, this study could be used to inform population genomics parameters and monitoring practices for the sustainable harvest and management of other similar long-lived species.

## Materials and Methods

### Fishing pressure, Traditional Ecological Knowledge

Currently, there is no mechanism in place for indigenous fisheries to report the number of fish caught in Mistassini Lake. Thus, to establish trends in fishing pressure, fish abundance and body size, we conducted semi-directed interviews as in Fraser et al. (2006) during February and July of 2018 with 17 elders and fisherman (30 – 79 years of age, with 13 respondents > 40 years) (Table 2). Answers were not used for questions where respondents explicitly stated a lack of knowledge as per Gagnon and Berteaux (2009), and the frequency of respondents for a given answer has been provided, using the total number of respondents for that question as the denominator. In addition, we obtained census numbers for all people in the community close to the lake and the number of fish caught by non-Cree fishers for a subset of years (Table S1). Ethics certificate 30008247.

Rivers included in the study were Chalifour, Icon and Perch in the south and Takwa in the north. Communicated by two TEK respondents and incidentally by several Cree fishers and community members in 2017 and 2018, Takwa was perceived to be relatively unaffected by fishing until ∼2015 when larger boat motors made access easier.

### Fish sampling

Fish were sampled during spawning (after ice-off: mid May in the south and early June in the north) at spawning rivers in 2002 and 2003 by Dupont et al. (2007) (historical), and in 2015, 2016 and 2017 (contemporary) by us (see Table 1 for sample sizes). Sampling was collaborative with subsistence fishers for 2015 – 2017. Walleye were captured via angling using the same lures and a combination of boats and shore fishing, from the same locations within rivers, for both historical and contemporary sampling (Table 1). Catch-per-unit-effort was not available for historic samples or collaborative sampling and is therefore not included here for contemporary sampling. After capture, fish were immediately placed in freshwater baths with aerators. From each walleye, we collected total and fork length (TL ± 1 mm), wet mass (± 50 g), sex (M, F, U (unknown, either spawned out or premature)) and a tissue sample for genetics; otoliths were collected from a random subsample. Live walleye were returned to the water near the location of capture. Opercular bones were collected for aging for historic samples (Table 1); 2015 and 2017 otoliths were aged at the Wisconsin Cooperative Fishery Unit, US Geological Service, University of Wisconsin, Stevens Point, USA.

### Body size at spawning and size-at-age

We modeled both body size (total length and mass) and size-at-age in this study because we had a far greater sample size for body size estimates than aged samples, and mass estimates had not been correlated to historic aged samples. Thus, evaluating body size allowed us to investigate changes in length and mass on a per-river, per-sex and per-year basis.

#### Body size

We used multiple regressions and ANOVA in R (R Core Team 2017) to test our prediction that body size of breeding adults had been reduced in southern rivers between 2002/03 and each 2015, 2016 and 2017. Year was set as a factor. Error was normally distributed for total length (TL), and mass was log transformed to improve fit of the error term. Our full model for each TL and mass (*Y*_*i*_) included the following.

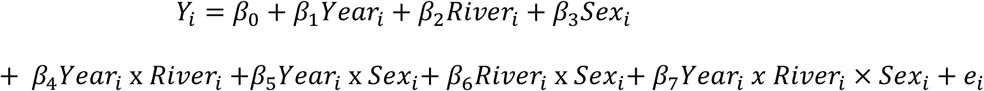

To determine the best model, we used backward step-wise model selection and AIC (Akaike 1974). Significance was detected at an alpha of 0.05 and all multi-comparison *P*-values were adjusted using the false discovery rate (FDR) method for 64 planned contrasts (Tables S3 and S5) (Benjamini and Hochberg 1995).

There were insufficient samples collected across locations in 2002 and 2003 to use these years independently for body size analysis. Since no population genetic structure existed between 2002 and 2003 within rivers (Dupont *et al*. 2007), they were combined for all analyses, and denoted as 2002/03. In addition, in 2017 we were unable to collect any female walleye from Perch River, nor a sufficient number of female walleye in Icon River to be able to use them in length/mass models (see Table 1 for sample numbers).

#### Size-at-age

To test our prediction of reduction in size-at-age in the southern populations relative to the northern one through time, we used a Bayesian hierarchical regression model. We used Bayesian as opposed to frequentist modelling to account for possible bias due to sampling gear, as well as small and variable sample sizes for aging structures across rivers. In addition, only total length was modeled because mass data were not included in the historical dataset containing age information. We assumed walleye total length (TL) was normally distributed such that:

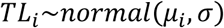

with shape parameters *μ*_*i*_ and *σ* representing the mean and standard deviation for walleye total length, respectively. Mean total length for the *i*th walleye was then modelled using linear regression:

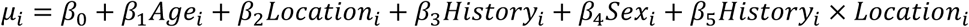

We used vague normal priors for all *β* coefficients and modelled hyperpriors for age and sex by river. Location (southern rivers vs northern river), and history (contemporary vs historic samples) were coded as categorical variables.

The Bayesian model was run using JAGS version 4.3.0 (Plummer 2003) in R, using *rjags* and *run*.*jags* (Denwood 2016). We described the posterior distribution for the model using four MCMC chains. Starting parameter values for each chain were jittered. Each chain took 20,000 samples of the posterior, thinned at a rate of 50. The adaption period was 1000 iterations, and a burn-in rate of 50% was used, for a total chain length of 2,050,00. We evaluated MCMC chain convergence by visual inspection of trace-plots to assess mixing. Additionally, we ensured that each parameter had effective samples sizes >1,000 and that they passed the Gelman-Rubin diagnostic test with potential scale reduction factors (PSRF) <1.1 suggesting convergence on a common posterior mode (Gelman *et al*. 2013).

### Sequencing

DNA was extracted using a modified Qiagen blood and tissue kit protocol (Qiagen Inc., Valencia, CA) (see Table 1 for sample sizes) and was sequenced using individual-based genotyping-by-sequencing (GBS). Libraries for Ion Proton GBS were prepared using the procedure described by Masher et al. (2013) at IBIS, Université Laval, Québec, Canada, with modifications described in Abed et al. (2018). Libraries were prepared for sequencing using an Ion CHEF, Hi-Q reagents and P1 V3 chips (ThermoFisher), and the sequencing was performed for 300 flows. Enzymes used to cleave the DNA were rare cutter *pst1* and frequent cutter *msp1*.

Single-nucleotide polymorphisms (SNPs) were determined from raw sequence reads using the *stacks* pipeline v1.45 (Catchen *et al*. 2013), and *de novo* sequence alignment, on the supercomputer Guillimin from McGill University, managed by Calcul Québec and Compute Canada. Pre-processing of fastq files was completed using fastQC (https://www.bioinformatics.babraham.ac.uk/projects/fastqc/) to assess read-quality before and after using cutadapt (Martin 2011) to trim any remaining adapters and remove sequences <50bp. Our stacks parameter optimization method was similar to Mastretta-Yanes et al. (2015), but we did not estimate error rate because we did not have enough positive controls to do so. Final stacks parameters included default settings with the following custom options: within process_radtags, 80bp trim-length; ustacks, SNP model, alpha = 0.1 for SNP calls, -m = 7; cstacks, -n = 3; rxstacks, log-likelihood cut-off = −30 for SNP calls; *populations*, log-likelihood cut-off of −30 for SNP calls, choose single SNP, maf = 0.01, -r = 0.8. We ran *populations* twice. First, we used the parameters listed here, and generated a blacklist of loci consisting of loci with F_IS_ < −0.3. We then re-ran *populations* with the same parameters listed here using this blacklist, with each – p=6/8, 5/7 and 4/6. No negative controls produced stacks, and all positive controls assigned to the correct populations using Discriminant Analysis of Principal Components (DAPC) analysis in Adegenet (Jombart 2008; Jombart *et al*. 2010). After quality trimming and filtering, an average of 8457 high-quality SNPs were used to estimate population structure, genetic diversity and effective population size (Ne).

### Population structure

To test our prediction that harvesting in the southern populations would change genetic population structure, potentially homogenizing structure between southern rivers over time, we assessed population structure using DAPC, ADMIXTURE (Alexander *et al*. 2009), and genetic distance (F_ST_) (using GenoDive and 999 permutations, Meirmans & Van Tienderen 2004; Weir & Cockerham 1984). The optimal number of principal components (PCs) to retain for DAPC was determined using the xval procedure, using n/3 (recommended by the manual) as the maximum number of PCs allowable, and 500 replicates. The population grouping that best fit the data was assessed using Bayesian Information Criterion (BIC) for DAPC, while for ADMXTURE analysis, we used cross validation (CV) and 500 bootstrap replications. Both analyses were completed at least four times using different bootstrap values and numbers of replicates to ensure results were stable. Based on DAPC, it was clear that 28 individuals, primarily sampled in Icon and Chalifour Rivers, were from different unsampled genetic source populations. We removed these individuals, re-ran the populations module of stacks, and conducted all subsequent analyses using this reduced dataset.

### Removing loci potentially under selection

Global outlier loci (loci putatively under selection) were detected using PCAdapt (Luu *et al*. 2017), using the scree plot method to determine the best number of PCs (K) to retain (Jackson 1993), and Mahalanobis distance with alpha = 0.1 to determine outliers. PCAdapt does not use pre-defined population structure, but instead ascertains structure based on PCA. The program detects outliers based on how they relate to the structure of populations on the PCA (i.e., the distance between a point and a distribution). After removing the outlier loci from the dataset, the effect of linkage disequilibrium (LD) on population structure was assessed by finding markers that were in LD (r^2^ = 0.7) using plink v1.9 (Chang *et al*. 2015). We removed these markers (n = 507) and re-analyzed population structure with DAPC. Since linked loci had no effect on structure, they were retained for all subsequent analyses. Genetic diversity, F_ST_ and Ne analyses were completed both including and excluding global outlier loci – while there was little difference between results including or excluding outlier loci for genetic diversity and Ne, the magnitude of F_ST_ was greater with outlier loci included, and the conclusion changed in one case for F_ST_. Thus, we have included only results for neutral loci here for these two metrics.

### Genetic diversity and Ne

To test the prediction that genetic diversity and Ne were reduced over time in southern populations, genetic diversity estimates were obtained using the *populations* model of *stacks*, and per-generation Ne (5-7 year generation time) was estimated using the linkage disequilibrium method in NeEstimator v2.01 (Do *et al*. 2014). Ne estimates and confidence intervals were corrected for linkage by correcting for chromosome number according to Waples et al. (2016).

### Signatures of selection

To test the prediction that signatures of selection would be most evident between timepoints within southern and not northern river(s), and that putatively selected loci would be associated with relevant biological processes, analyses to determine outlier loci were conducted with PCAdapt using the method described above. Analyses were conducted both for sexes combined and separately: (i) for all populations combined, (ii) for southern rivers only, and (iii) within each population. For all analyses, except when all populations and years were included, the *populations* module of *stacks* was re-run including only the populations and/or sexes that were being contrasted, specifying that loci had to be present in both populations (see Table S7 for sample sizes and numbers of loci in each analysis). For sex-based analysis of the full dataset including all rivers and years, loci were required to be present in 6 of 8 populations.

To determine possible functions for outlier loci, for all within-river historic-contemporary contrasts for which there were outlier loci, FASTA files were blasted, mapped and annotated using blast2go (Götz *et al*. 2008). Default parameters were used with the following custom choices: proprietary cloudblast, fast-blast, UniProtKB/Swiss-Prot (Swissprot_v5) database, blast e-value 1.0E-5, 10 blast hits, filtered GO by taxonomy taking only matches to animals (Metazoa).

## Results

### Fishing pressure, Traditional Ecological Knowledge

Of 17 elders and fishers, most reported reductions in the size and number of walleye caught in the lake (15 and 14 respondents respectively) within the last 5-20 years (Table 2). Eleven respondents expressed concerns directly about overfishing, fishing during spawning or taking of too many fish during spawning. There was no consistent change in the number of fish caught by non-Cree fishers between 1997 and 2015 (virtually the same in 1997, 2011 and 2015, but 54% higher in 2003) (Table S1), but the community of Mistissini (3724 people in 2016) grew by ∼29% between 1997 and 2011, and the population and number of households in the community increased by ∼50% between 1997 and 2016 (Table S1). In addition, while there is currently no data on the number of fishers in the community nor the proportion of the population that fishes, the majority of fishers (16/17) fished in the southern area of concern in the lake, more than double than in all other areas of the lake except Takwa River (where 9/17 fishers fished) (Table 2).

### Body size at spawning and size-at-age

#### Body size

Our prediction that body size would be reduced in southern populations within a 1-2.5-generation period was supported. The main effects, year, river and sex all had a significant effect on total length and mass (Table 3), and the best-fit models for each TL and mass each contained a three-way interaction between year, sex and river. AIC was >10 better for the full model for TL and >6 better for mass. Both regression models were significant (TL, R^2^ adj = 0.418, F_27, 1486_ = 41.24, *p* < 0.001; mass R^2^ adj = 0.3907, F_27, 1481_ = 36.81). Mean TL and mass decreased significantly for both sexes between 2002/03 and each of 2015, 2016 and 2017 in the southern rivers (TL 7 – 21% and mass 22-47% reductions), except for Perch River males in two contrasts and Chalifour females in two contrasts. While there was no significant change between 2002/03 and each of 2016 and 2017 years for females in Chalifour, the trend in decline remained clear (Figure 2); lack of significance may relate to low female sample size in this river (Table 1). Indeed, the trend for Chalifour females was particularly evident when contrasted to Takwa (the reference northern river), where mean sizes of fish were consistent across all sampling years. Finally, a sex-bias for more males than females being captured at spawning sites was consistent for all sampling years and for all rivers, including Takwa (Figure S1).

**Table 3:**
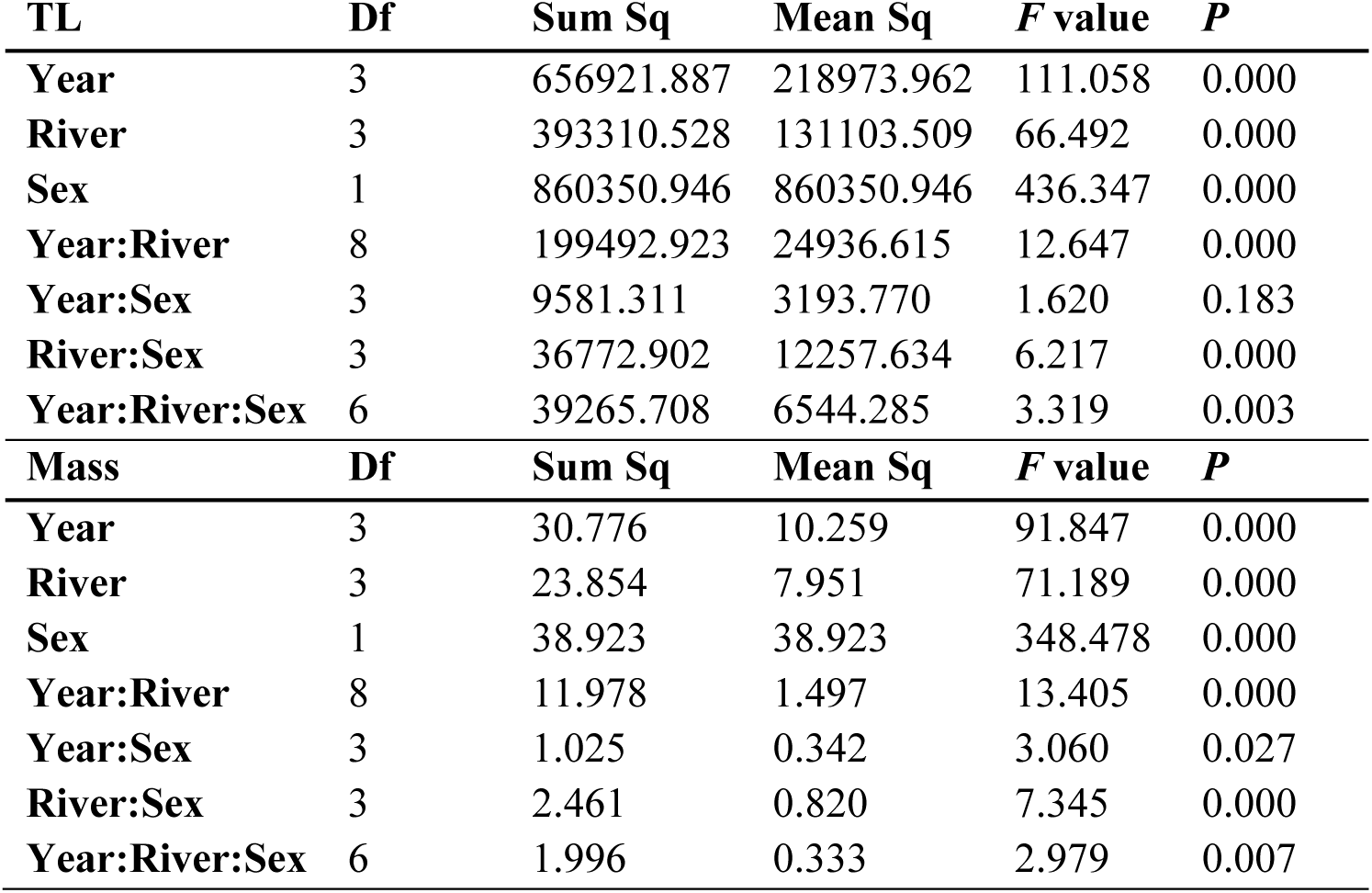
Analysis of variance table for best fit (full) model for each walleye total length (TL) and mass, with response log(mass). Years are each 2002/03, 2015, 2016 and 2017. Rivers are Chalifour, Icon, Perch and Takwa.

**Figure 2:**
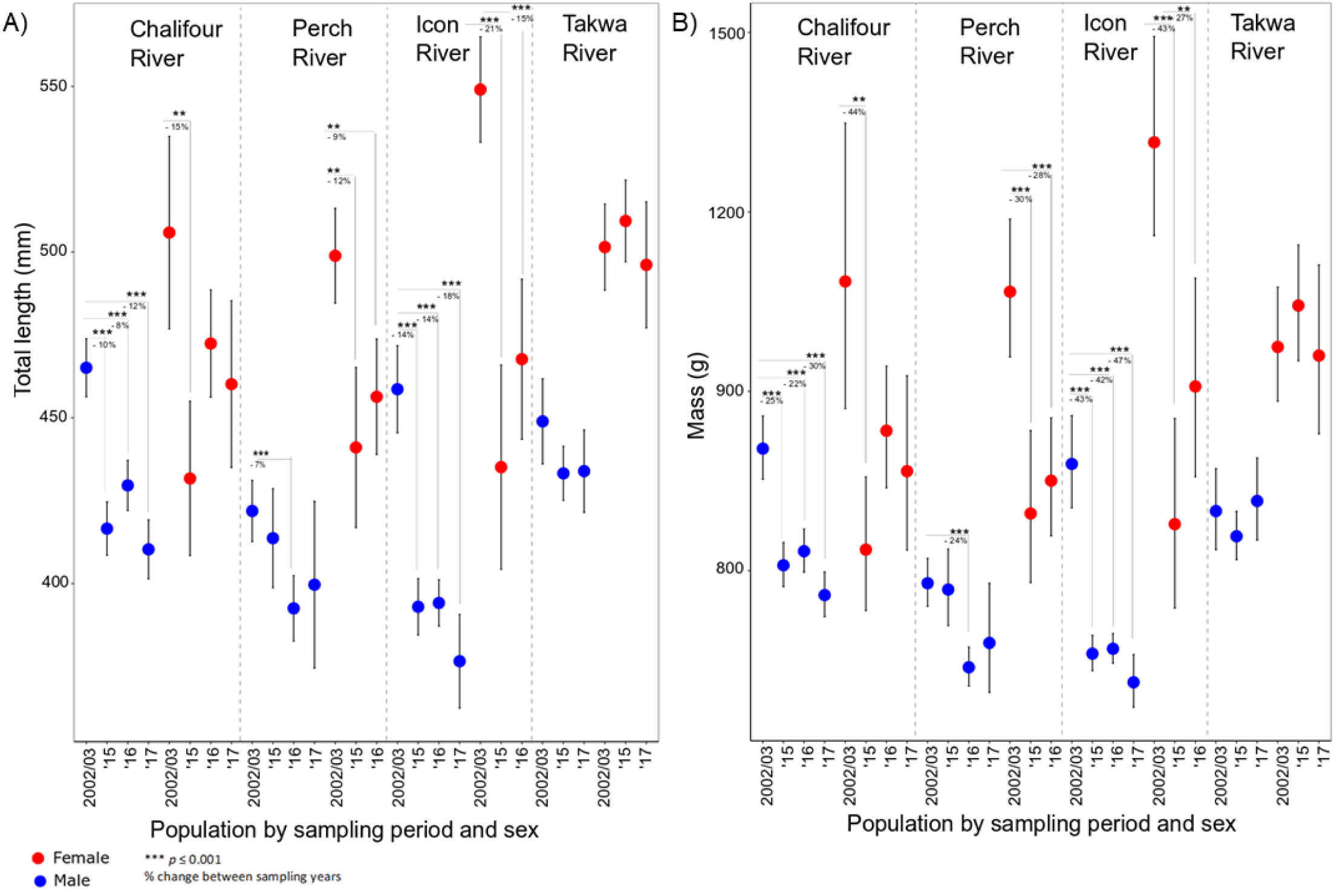
Least squares means (± 95% CI) of A) total length and B) mass for male and female walleye between 2002/03, 2015, 2016 and 2017 in the four rivers surveyed. There was also a significant change between 2016 and 2017 (*p* = 0.0092 for TL and p = 0.003 for mass) for male fish in Chalifour, but this was not shown for clarity.

#### Size-at-age

Our prediction that fish in the southern rivers would be smaller for their age through time was supported, although the reduction in size was small (See Table 4 for posterior means and 95% credible intervals). Overall, fish were larger in the southern than the northern river(s), over all rivers fish were 29.4 mm larger contemporaneously than they were historically, and males were 48.2 mm smaller than females on average. Lastly, and the term that tested our hypothesis, fish in the southern rivers were 13.7 mm smaller relative to fish in the north in contemporary relative to historical samples.

**Table 4:** Posterior means (PM) and 95% credible intervals (CI) for walleye size-at-age model for total length (in mm). Age is the effect of each year on length, location is south versus north and history is contemporary versus historical.

| Parameter | Parameter | PM (mm) | 95% CI | Percent of the posterior mass below 0 |
| --- | --- | --- | --- | --- |
| $\beta_0$ | Intercept | 463.8 | 418.7-506.4 | 0% |
| $\beta_1$ | Age | 10.8 | 8.6-12.9 | 0% |
| $\beta_2$ | Location | 27.8 | -9.7-80.4 | 6% |
| $\beta_3$ | History | -29.4 | -50.0- -9.3 | 100% |
| $\beta_4$ | Sex | -48.2 | -65.6- -32.6 | 100% |
| $\beta_5$ | History x Location | -13.7 | -37.1-9.6 | 87% |

All parameter estimates passed convergence checks. Each parameter had effective samples sizes >1,000 and passed the Gelman-Rubin diagnostic test with potential scale reduction factors (PSRF) <1.1, suggesting convergence on a common posterior mode (Gelman et al. 2013).

### Population structure

Our prediction that population structure would change over time, possibly including homogenization of structure between southern rivers, was supported for Icon and Perch Rivers in two of the three analyses. Specifically, ADMIXTURE and F_ST_ analyses supported this prediction, while DAPC did not. Using both DAPC and ADMIXTURE 3 populations best described the data (k = 3053.162 and CV = 0.421 respectively) (Figure 1), but the difference between CV 2-4 was small for ADMIXTURE (CV of 2 = 0.423 and CV of 4 = 0.428). In the 3-population scenario, Icon and Perch grouped as a metapopulation and Chalifour and Takwa Rivers were independent, with 2002/03 and 2015 samples grouping together for each river. In a 4-population scenario, many Perch 2003 individuals showed a substantial fraction of loci that were different from the Icon-Perch group. Genetic differentiation by F_ST_ mirrored what was evident in the K = 4 scenario. F_ST_ showed weak differentiation between Perch and Icon in 2002/03 (Table 5), and then merged as a single metapopulation in 2015. At a within-population level, Chalifour River was differentiated between timepoints, Icon did not diverge between timepoints, Perch diverged marginally between timepoints, and Takwa did not diverge between timepoints. Given that k = 3 was identified as the best structure overall, subsequent genetic diversity and Ne analyses were conducted using a meta-population structure for Icon-Perch, with Chalifour and Takwa identified separately, but years were defined as separate populations for temporal analysis.

**Table 5.**
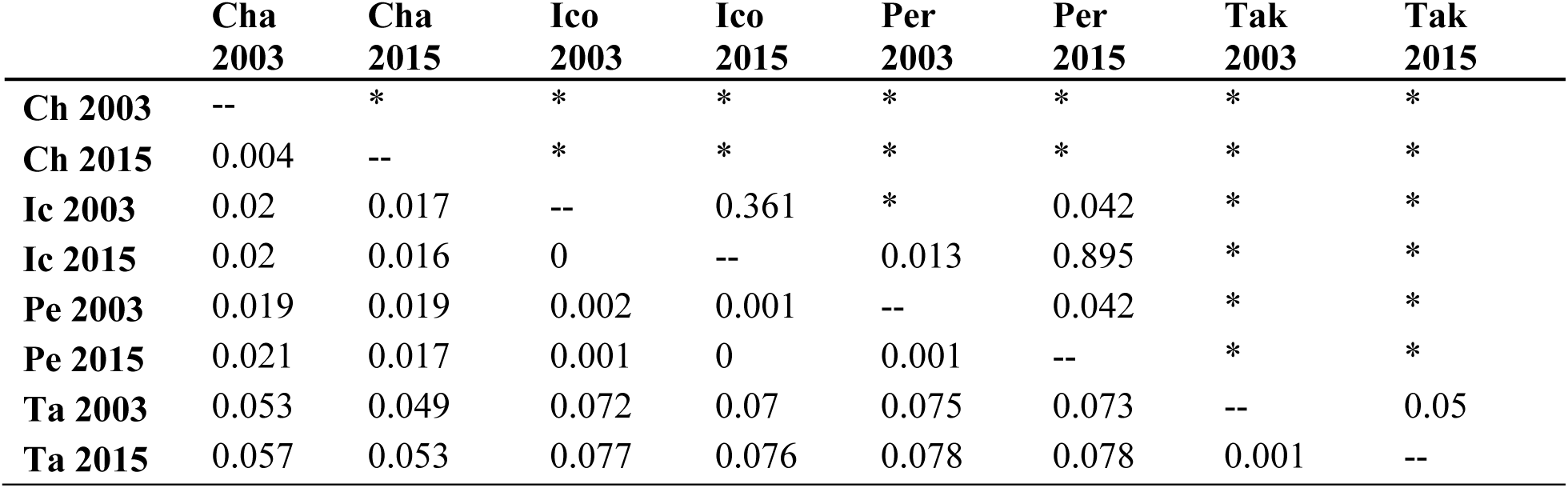
F_ST_ differentiation for walleye within and between rivers for each year sampled. F_ST_ estimates are below the diagonal, and *p*-values are above the diagonal, with a “*” indicating *p* ≤ 0.001.

|  | **Cha**<br>**2003** | **Cha**<br>**2015** | **Ico**<br>**2003** | **Ico**<br>**2015** | **Per**<br>**2003** | **Per**<br>**2015** | **Tak**<br>**2003** | **Tak**<br>**2015** |
| --- | --- | --- | --- | --- | --- | --- | --- | --- |
| **Ch 2003** | -- | * | * | * | * | * | * | * |
| **Ch 2015** | 0.004 | -- | * | * | * | * | * | * |
| **Ic 2003** | 0.02 | 0.017 | -- | 0.361 | * | 0.042 | * | * |
| **Ic 2015** | 0.02 | 0.016 | 0 | -- | 0.013 | 0.895 | * | * |
| **Pe 2003** | 0.019 | 0.019 | 0.002 | 0.001 | -- | 0.042 | * | * |
| **Pe 2015** | 0.021 | 0.017 | 0.001 | 0 | 0.001 | -- | * | * |
| **Ta 2003** | 0.053 | 0.049 | 0.072 | 0.07 | 0.075 | 0.073 | -- | 0.05 |
| **Ta 2015** | 0.057 | 0.053 | 0.077 | 0.076 | 0.078 | 0.078 | 0.001 | -- |

### Genetic diversity and Ne

Our prediction that genetic diversity and Ne would be reduced between historical and contemporary timepoints was not supported. Genetic diversity fell within a tight range for all populations over all years, ranging from 0.21 to 0.23, with the lowest and highest values being in the southern populations (Figure 3a). Confidence Intervals (CIs) overlapped between timepoints for H_E_ in Chalifour and Icon-Perch; there was a 4.9% loss in H_E_ in Takwa, though the reduced H_E_ still fell within the range of southern populations. Point estimates of Ne ranged from 1741 to 3146 individuals across all populations, with the lowest and highest values being in the northern population. Ne CIs also overlapped between timepoints in Chalifour and Icon-Perch Rivers, and the data suggested a doubling in Ne over time in Takwa River (Figure 3b). There were likely insufficient samples to accurately detect a difference between thousands of individuals (Nunziata & Weisrock 2018; Waples & Do 2010) however these results clearly show that all populations remained large (i.e., Ne in the thousands).

**Figure 3:**
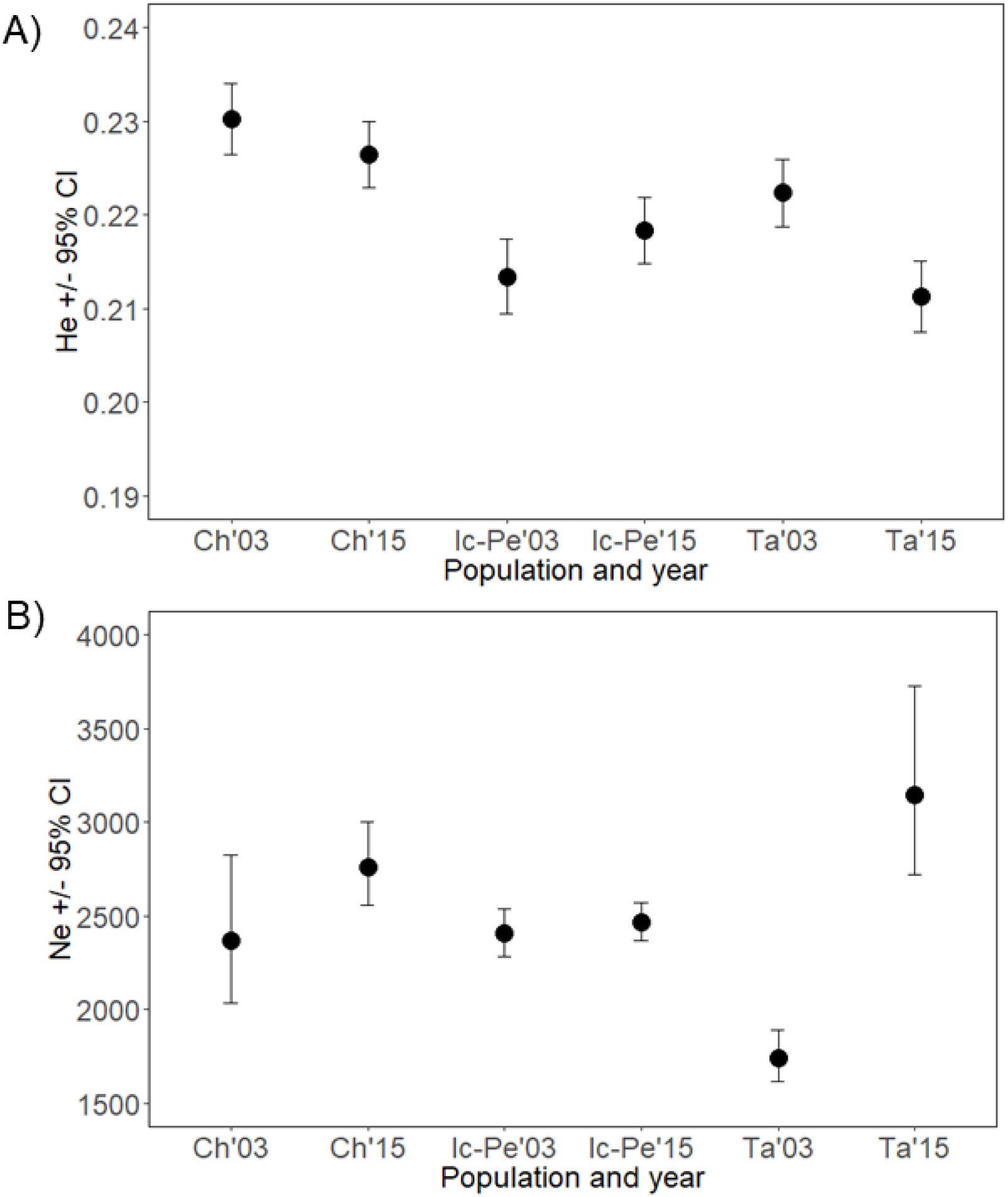
**A**) Expected heterozygosity (H_E_) ± 95% CI and B) effective population size (N_E_) ± 95% CI for walleye from each Chalifour (Ch), Icon-Perch (Ic-Pe) and Takwa (Ta) Rivers in each 2003 and 2015.

### Signatures of selection

Our prediction that signatures of selection would be present between historic and contemporary timepoints within southern rivers but not the northern river, with putatively selected loci related to growth, body size and/or maturation, was supported. Eleven to 263 loci were outliers (0.17-2.83%) (Figures 4a - b). Outliers were found in the global PCA that included all rivers and both timepoints, for each F/M together and F & M individually, and Perch 2003 clustered as a separate population. But removing these outliers did not change the population structure in most cases. On the contrary, when the southern rivers were analyzed as a unit (i.e., the south historic vs contemporary) and on their own (i.e., each Chalifour and Icon-Perch historic vs contemporary years), for F/M combined and for each F & M separately, population structure existed between the two timepoints and removing outlier loci usually collapsed the population structure. Further, there was no population structure between timepoints in Takwa (and thus no outlier loci) (Figure 4c, Table S7 and figures S2-16). In sum, while there were outliers separating north from south, outlier SNPs disproportionately contributed to the PCs between years in the southern subset. In addition, the greatest proportion of outlier SNPs found in each historic/contemporary PCA in the south maintained population structure between years (Figure 4c). Lastly, parallel outliers existed between the southern rivers (Figure 4d); of note, more outliers were in common between Icon-Perch F and Chalifour M (18 outliers) than between Icon-Perch F and Chalifour F (0) or Chalifour M and F (3).

**Figure 4.**
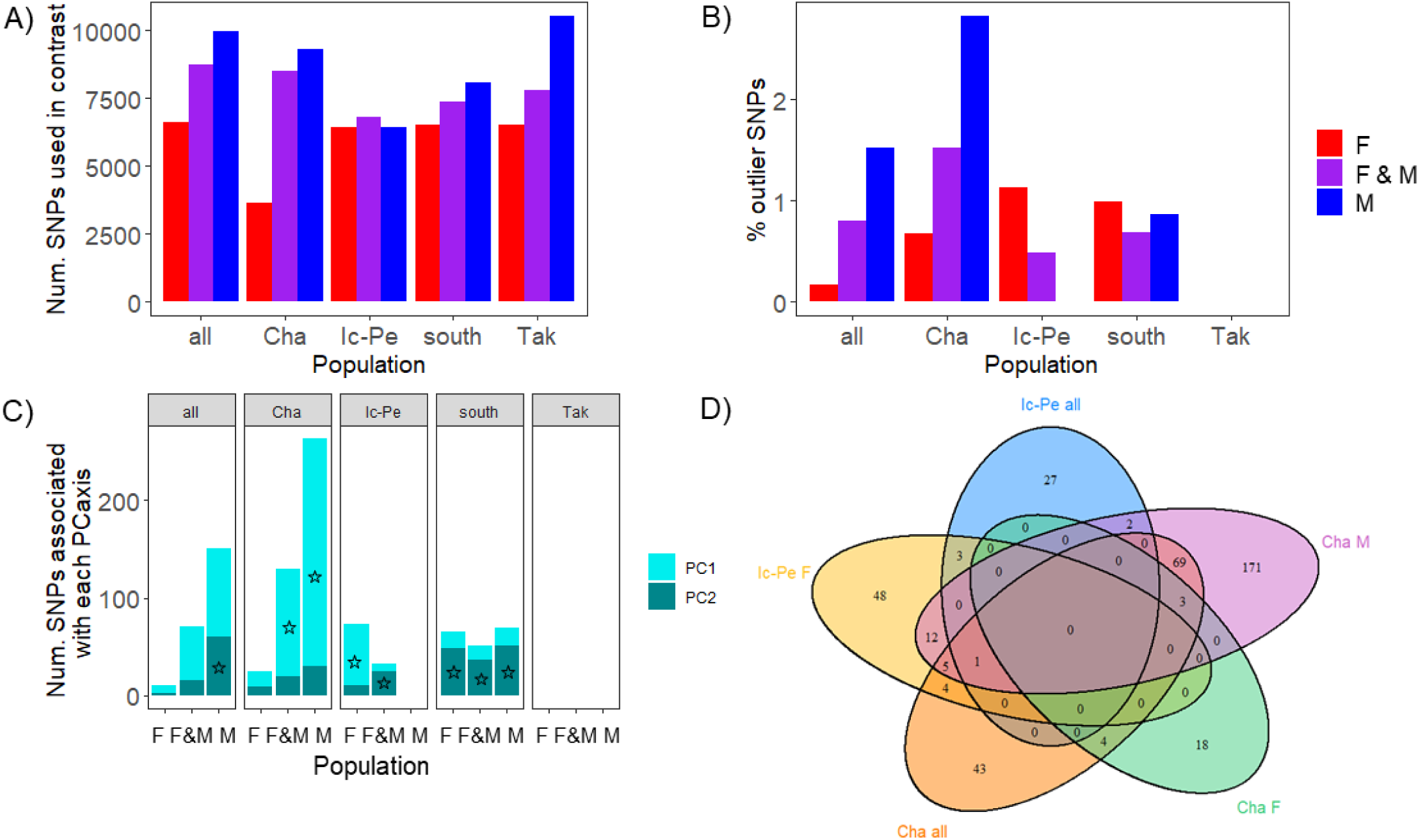
A) Total number of SNPs used in each between-year contrast (i.e., the number of SNPs used was unique for each sex and population). B) Percentage of SNPs that were outliers in each pairwise contrast. C) Number of outlier SNPs associated with each PCaxis. The star (☆) denotes which PC axis separated years. Where there is no star on a bar population structure between years was not maintained by outlier loci. See Table S7 and Figures S2 – S16 for further explanations and detailed descriptions of how outliers on each PC axis maintained the observed population structure. Where no bars are shown (i.e., for Takwa and Icon-Perch m in % outlier SNPs and SNPs associated with each PCaxis) there was no population structure and thus no outlier loci between years. D) Outlier loci overlap between the southern populations.

Sex-specific analyses provided greater resolution than with sexes combined (Figures 4c and d). For example, outlier SNPs maintained population structure between timepoints in more of the southern rivers in sex-specific rather than combined analyses (Table S7, Figures S2-7). In addition, there was no population structure in Icon-Perch males, but there was in females (Figures 4c and S12-13). Lastly, the number of outliers in common between rivers differed between the sexes (Figure 4d), though this may be due in part to the number of individuals sequenced.

Blasts were conducted for southern (F/M combined and separately), Chalifour (F/M combined and separately), and Icon-Perch F (no outliers were found for Icon-Perch M or Takwa) Rivers. Southern F/M combined and M had no blast hits. Otherwise, between 2 and 6 alleles were annotated for each blast. Because annotations were completed against all mapped Metazoa, at level 2 go-annotation, functional annotations included many different biological processes, molecular functions and cellular components. Three relevant processes indicated were growth, metabolism and developmental process (Table S8).

## Discussion

We detected rapid genetic changes associated with harvesting wild populations of walleye within 1-2.5 generations, a shorter timescale than previously observed in other fisheries. Concurrent with reductions in body size within a 15-year period (2002/03 – 2017) (Figure 2), we detected a reduction in size-at-age (Table 4), genomic change evidenced by changing genomic population structure (Figure 1, Table 5) and putative signatures of selection within rivers (Figure 4), both with sexes combined and separately. These changes were present in the southern rivers most-impacted by increased fishing pressure by Cree and non-Cree fishers alike (Tables 2 and S1), and not in the northern river where there were fewer boats and fishers. Importantly, not only is fishing pressure greatest in the south, but southern fish from the affected spawning runs remain close to those spawning runs in the summer mixed population fishery (Dupont *et al*. 2007). A genetic drift hypothesis could posit that the observed genomic changes are stochastic. However, all of the observed populations had large Ne, making it unlikely that drift caused the phenotypic or genetic changes observed. The difference between neutral and adaptive genomic results, however, illustrates the capacity for genetically-large populations in nature to rapidly respond to harvest-induced selective pressures.

Reductions in body size (or trends indicating such reductions) were consistent between 2002/03 and each of 2015 – 2017 within all southern rivers, except for Perch River male body size (Figure 2); moreover, size-at-age was reduced in the south over time (Table 4). In fact, size reductions in all southern rivers were likely underestimates of the true change. Namely, 2016 monitoring was largely collaborative; approximately 48% of all sampled walleye (216 of 446) were harvested and donated by fishers. Donated 2016 walleye were 639 g (stderr ± 21 mm) and 424 mm (stderr ± 4 mm) on average compared to 603 g (stderr ± 20 g) and 410 mm (stderr ± 4 mm) from our caught and released 2016 walleye (note that sampling was not collaborative in this way in 2002/03). Lastly, given that Perch River males were smaller than females, it is less likely that they would be subject to size-selective harvesting. In sum, these results support the idea that fishers often target larger fish, and this type of size-selective harvesting has been documented to lead to the evolution of smaller body size (Heino *et al*. 2015; Hutchings 2005; Swain *et al*. 2007).

Genomic change occurred between timepoints in the southern Rivers but not in the northern river. Population structure was homogenized over time between Icon and Perch Rivers. Signatures of selection were evident within southern Rivers: rivers were genetically differentiated between years (Table 5), outlier loci maintained that structure (Figure 4, Table S7), and exploratory analysis revealed relevant biological functions associated with a small number of those outlier loci (Table S8). In addition, although the extent of parallelism in events of natural selection is variable (Oke *et al*. 2017), parallel outlier loci were detected between timepoints in the southern rivers (Figure 4c). However, genomic change was clearly nascent. Neutral genetic diversity did not change between timepoints. Differences between the preferred population structures in ADMIXTURE were small (Figure 1). F_ST_ within rivers between years and between Icon and Perch 2015 were small (Table 5), and scree plots for outlier locus detection showed weak structure in two cases (Figures S9 and S12). Nonetheless, results were generally consistent when sexes were analyzed together and separately; although congruent with reductions in body size between 2002/03 and 2015 in Perch River, genetic structure was present between timepoints in Icon-Perch females but not males (Figure 4).

Alternative explanations for the genomic change evident in Icon-Perch include sampling bias, spatial movement, or increased gene flow. If sampling bias was present, Perch 2003 individuals could have been from a different population, but 2002/03 were genetically indistinct within rivers (Dupont *et al*. 2007), and Perch 2003 grouped with 2015 samples by DAPC. Alternatively, Perch 2003 individuals that were different historically could have moved to a different spawning location in later years (Bigrigg 2008), though Mistassini Lake populations generally have strong spawning site fidelity (Dupont *et al*. 2007). Another possibility is that individuals from Icon River could be using Perch River to spawn much more now than historically, either replacing genotypes that have been fished out, or increasing gene flow substantially (Allendorf *et al*. 2008). Although the observed neutral and putatively selective genomic change in Icon-Perch is rapid, it is not without precedent (3 generations or less, Chebib *et al*. 2016; van Wijk *et al*. 2013), and even though these are genetically large populations (Figure 3a), rapid adaptation is possible via soft sweeps (Hermisson 2005; Messer & Petrov 2013). In sum, nascent genomic change occurred within a 12-year period (genomic samples were 2003 and 2015) within the southern most-harvested rivers, which represents 1-2.5 generations maximum.

Our data are consistent with rapid genetic changes to population structure and genetically-based phenotypes due to harvest, but alternative explanations must be explored. The observed reduction in size-at-age in the south was small relative to the overall change in body size between historical and contemporary samples, and may be due to a difference in aging structures used (historical using opercula, contemporary using otoliths) (Faust & Scholten 2017). However, ages calculated using opercula and otoliths have been highly correlated in walleye (Geisler 2012), and opercula have been validated to the age of 16 in walleye (94% of aged fish in Mistassini were < 16) (Faust & Scholten 2017).

Another alternative could be that the large body size changes are entirely plastic due to changes in the environment or a habitat shift unrelated to fishing. However, climate change is expected to warm the Mistassini region; as a cold oligotrophic lake, Mistassini is not ideal habitat for walleye, which prefer mesotrophic lakes (Kitchell *et al*. 1977; Niemuth *et al*. 1972). Climate warming is expected to increase the growing season length for walleye, and thus in the absence of fishing an increase in body size is a more likely response with climate warming than a decrease. Regarding plasticity, growing degree day (GDD) was shown to account for 96% of the variation in length of immature walleye over 416 populations in Ontario and Quebec, though variation in growth associated with food availability was also evident (Neuheimer & Taggart 2007; Venturelli *et al*. 2010). Thus, although it is unlikely that all observed changes are due to selection, smaller body size at spawning and smaller size-at-age could indicate that fish are selectively growing slower (Enberg *et al*. 2012).

Under variable recruitment (Bozek *et al*. 2011; Hansen *et al*. 1998), fish captured in 2002 and 2003 could represent distinct, strong year classes, biasing estimates of mean size and contributing to temporal genetic differences. However, there was no genetic differentiation within rivers between 2002 and 2003 (Dupont *et al*. 2007), Takwa River walleye had consistent body size in all years sampled, and we found a significant reduction in size-at-age in the southern rivers.

We estimated the selective pressure required to generate the observed changes in body size within 1-2.5 generations to assess whether they were biologically plausible using the breeders equation (*R = h*^*2*^*S*, where *R =* response to selection, *h*^*2*^ *=* heritability, *S* = selection differential). Using averages for 11-year old walleye in the south for each 2002 (515 mm) and 2015 (424 mm) (a 13-year interval), *R* = −7 mm per year. Given realistic *h*^*2*^ estimates (0.3) (Law 2000; Nussle *et al*. 2009), if observed changes were entirely due to selection, *S* would need to be −23 mm per year. Our Bayesian model held all variables constant when assessing the size change associated with harvest, and in so doing found that the size change attributed to selection was smaller than what is shown here. Thus, it is very likely that some of the observed body size change was plastic and/or due to stochasticity in addition to selection pressure.

### Conclusions and management implications

We have presented coupled phenotypic and genetic evidence consistent with fisheries-induced genetic changes within 1-2.5 generations in wild walleye populations; with rare comprehensive evidence, this study sets a precedent for the timeframe needed for investigating concerns regarding harvest-induced evolution in fisheries. Furthermore, sex-specific dynamics for both body size and genomics herein highlight the importance of collecting sex-specific data.

Our study illustrates the power of integrating life history and genomics methods for conservation in order to understand the factors affecting population change (Bernatchez *et al*. 2017), of integrating these with TEK, and of iterative population monitoring practices (Flanagan *et al*. 2018); i.e., this study would not have been possible without the historic data. Considerations for Cree management could include that observed phenotypic and genetic changes may cause reduced productivity (Allendorf *et al*. 2008; Hutchings 2005), and that genomic change is clearly nascent here. Depending on the severity of harvest (which is not precisely known in this case) and life-history (Audzijonyte & Kuparinen 2016), fisheries-induced changes may be reversed in 9 generations (Conover *et al*. 2009) or less (Feiner *et al*. 2015) if fishing is halted.

## Data accessibility

Supporting data are available on Dryad: to be completed after the manuscript has been accepted for publication.

## Acknowledgements

We thank many people who contributed to this work. PP Dupont and L Bernatchez generously shared their historical tissue samples and length weight data. JP Coon-Come, D Schecapio, E Coon-Come and D Petawabano were excellent Cree guides. M Rabbitskin, our interpreter, and 17 anonymous Cree fishers and elders shared their knowledge of the species and the lake. R Arax Koumrouyan helped conduct the blast2go analysis. M Yates, E Lawrence, J-M Matte, A Prevost, N Hill, A Harbicht, B Brookes, P Peres-Neto, A Cantin, K Wilson, B Allen, W Larson, Z Feiner and D Isermann contributed critical feedback and discussion throughout the project and/or on drafts of the article. M Dunn facilitated engagement with Niskamoon Corporation. This project was funded by fisheries monitoring grants from Niskamoon Corporation to EB/DJF/PM, and by a Mitacs Elevate Postdoctoral Fellowship to EB.

