## Supplementary tables and figures for "Size reductions and genomic changes associated with harvesting within two generations in wild walleye populations"

### Supporting Tables

### Supporting Figures

Figure S1 Sex-bias at sampling sites, where blue = males, red = females and grey = unknowns in 2002-03, 2015, 2016 and 2017. Also shown are the total captured walleye in each river (n). 8
Figure S2 All populations, all years, m and f combined, k=2. Top: scree panels for selecting best k. Bottom left, with outliers left in. Bottom right, excluding outliers. Result: PC1 separates the north from the south, and also Icon03/15Perch15 a bit, and the second PC separates Chalifour from the other pops. Shown in the SNP list that associates SNPs with PCs below, the vast majority of outliers are associated with PC1 (15 to PC2). However, removing outliers does not change the structure in a meaningful way. 12
Figure S3 All populations and years, f, k=2. Top panels are scree plots for determining the best k. Bottom panels are: (left) including outliers, and (right) excluding outliers. Result: PC1 separates north and south, and maybe Chali03/15 and Pe03 a little bit from the Ic03,15,Pe15 group. PC2 separates Ch03 from Ch15, and also both Chalifour years from the other southern pops. Result looks similar to when M and F are combined, except that PC2 also separates Ch03 and 15. Removing outliers has no meaningful affect on the population structure. number of pops that I required loci to be present in for the full dataset is 6/8. 15
Figure S4 All populations and years, m, k=2. Top panels are scree plots used to determine best k. Bottom panels are: (left) including outliers, and (right) excluding outliers. Result: PC1 separates north from south, and also Pe03 and the Chalifours. PC2 separates Cha03 and 15 from the other pops, Pe03 from the other pops, and Ch03 from 15. Removing outliers removes the structure between Ch03 and 15. 17
Figure S5 Southern rivers, m & f combined, k=2. Top right panel is with outliers included and bottom right panel is outliers excluded. Conclusions: structure is sufficient to detect outliers. PCI separates Ch from other pops, PC2 separates Ch 03 from 15 and Pe03. Removing outliers removes the structure. 22
Figure S6 southern rivers, f, k=2. Top panels are the scree plots used to find the best k. Bottom panels are: (left) including outliers and (right) excluding outliers. Result: PC1 separates Chalifour from the other

**sites and PC2 separates Pe03 and Cha03 from the other sites. Removing outliers collapses PC2 structure, collapsing Pe03 into the other southern rivers, and Ch03 and 15. This result is the inverse of the result when both sexes are analyzed together.** 24
**Figure S7 south, m, k = 2. Top panels are scree plots used to select the best k. Bottom panels are: (left) including outliers, and (right) excluding outliers. Result: PC1 separates Chalifours from the other sites and Pe03 from Ic. PC2 seaprates Cha03 from 15. Removing outliers collapses Cha 03 and 15, and also collapses Pe03 completely into the other sites.** 27
**Figure S8 Chalifour, m and f combined. Upper left is scree plot showing that k=2 is best. Top right is including outliers, bottom right is excluding outliers. Outliers play a role in generating population structure within this river between years, shown by changes in population structure when outliers are excluded.** 30
**Figure S9 Chalifour f, k = 2. Top, with outliers, bottom, without outliers. There is sufficient structure to determine outliers. Based on the number of individuals it isn't totally clear that the outliers are maintaining the structure present. Populations might overlap more when outliers are removed.** 34
**Figure S10 Chalifour m, k = 2. Top left panel is the scree plot showing best k. Top right is including outliers and lower right is excluding outliers. Result: PC1 separates the timepoints.** 36
**Figure S11 Ic-Pe, m and f combined. Top left is the scree plot showing that k = 5 looks best. However, contribution of PC3 to population structure was tested, and it does not change the conclusion. Thus, k=2 was chosen. Top right is with outlier loci included, and bottom right is with outlier loci excluded. Conclusion: although structure is relatively weak within Ic-Pe between years, there is structure, and outlier loci contribute to this structure. Pe03 separates out on PC2. In SNP list associated with PCs below, you can see that 8 SNPs are associated with this PC.** 43
**Figure S12 Ic-Pe f, k=2. Top panels are scree plots used to determine best k. Bottom plots are (right) with outlier loci included, and (left) excluding outlier loci. Result: PC1 separates Pe03 from the other populations.** 45
**Figure S13 Ic-Pe m, k = 4. Top panels are scree plots showing that there is insufficient structure to find a best k. Bottom panel is including outliers. Both k = 4 and k = 6 were tested with the same result.** 48
**Figure S14 Takwa, m and f combined k=2, The top panel is scree plots showing the best k. Bottom panel is with outliers included, and shows that there is no structure between years. The structure seen in the scree plot is likely due to those outlier individuals.** 49
**Figure S15 Takwa River, f, k=2. Top panels are scree plots used to find the best k. Bottom panel is including outliers. Result: There is no structure between years within Takwa River. Outliers cannot be assessed.** 50
**Figure S16 Takwa m, k=3. Top left panel showing scree plot used to find best k. Top right panel is including outliers. Bottom right panel is excluding outliers. Result: there is no population structure between years, and thus no outliers can be found.** 51

**Supplementary Tables**

*Demography*

*Table S1 Demographic trends for the population in Mistissini and documented number of fish harvested by non-Cree fishers. The fish caught by non-Cree fishers were all in the southern half of lake below the 51st parallel (num fish non-Cree). Aboriginal affairs branch, Ministry of health and social services, Government of Quebec (GC). Ministry of Forests, Wildlife and Parks, Government of Quebec (MFFP). Cree Nation of Mistissini (CNM), Statistics Canada (GC).*

| | 1997 | Source | 2003 | Source | 2011 | Source | 2015 | Source | 2016 | Source |
|---|---|---|---|---|---|---|---|---|---|---|
| **Population** | 2479 | GQ | 2863 | GQ | 3469 | GQ | 3654 | GQ | 3724 | GQ |
| **Num. of households** | 424 | CNM | 556 | CNM | - | - | - | - | 800 | GC |
| **Num. fish non-Cree** | 5800 | MFFP | 8916 | MFFP | 5572 | MFFP | 5725 | MFFP | | |

*Life History*

<u>Body size</u>

*Table S2 Total length (TL) lsmeans with the lower (LCI) and upper (UCI) 95% confidence intervals.*

| River | Year | Sex | lsmean | SE | df | LCI | UCI |
|---|---|---|---|---|---|---|---|
| Chalifour | Y2002/03 | F | 505.889 | 14.801 | 1486 | 476.855 | 534.923 |
| Chalifour | Y2015 | F | 431.714 | 11.867 | 1486 | 408.436 | 454.993 |
| Chalifour | Y2016 | F | 472.414 | 8.246 | 1486 | 456.240 | 488.588 |
| Chalifour | Y2017 | F | 460.167 | 12.818 | 1486 | 435.023 | 485.311 |
| Chalifour | Y2002/03 | M | 465.111 | 4.463 | 1486 | 456.357 | 473.865 |
| Chalifour | Y2015 | M | 416.581 | 4.105 | 1486 | 408.529 | 424.634 |
| Chalifour | Y2016 | M | 429.606 | 3.865 | 1486 | 422.025 | 437.187 |
| Chalifour | Y2017 | M | 410.333 | 4.532 | 1486 | 401.444 | 419.223 |
| Icon | Y2002/03 | F | 549.033 | 8.107 | 1486 | 533.131 | 564.936 |
| Icon | Y2015 | F | 435.125 | 15.699 | 1486 | 404.330 | 465.920 |
| Icon | Y2016 | F | 467.692 | 12.315 | 1486 | 443.535 | 491.850 |
| Icon | Y2002/03 | M | 458.636 | 6.694 | 1486 | 445.505 | 471.767 |
| Icon | Y2015 | M | 393.000 | 4.333 | 1486 | 384.500 | 401.500 |
| Icon | Y2016 | M | 394.161 | 3.567 | 1486 | 387.165 | 401.157 |
| Icon | Y2017 | M | 376.579 | 7.203 | 1486 | 362.449 | 390.709 |
| Perch | Y2002/03 | F | 498.892 | 7.300 | 1486 | 484.573 | 513.211 |
| Perch | Y2015 | F | 441.077 | 12.315 | 1486 | 416.919 | 465.234 |
| Perch | Y2016 | F | 456.360 | 8.881 | 1486 | 438.940 | 473.780 |
| Perch | Y2002/03 | M | 421.899 | 4.707 | 1486 | 412.666 | 431.132 |
| Perch | Y2015 | M | 413.706 | 7.615 | 1486 | 398.768 | 428.644 |

| | | | | | | | |
|---|---|---|---|---|---|---|---|
| Perch | Y2016 | M | 392.538 | 5.028 | 1486 | 382.676 | 402.401 |
| Perch | Y2017 | M | 399.667 | 12.818 | 1486 | 374.523 | 424.811 |
| Takwa | Y2002/03 | F | 501.511 | 6.619 | 1486 | 488.527 | 514.495 |
| Takwa | Y2015 | F | 509.400 | 6.280 | 1486 | 497.082 | 521.718 |
| Takwa | Y2017 | F | 496.143 | 9.690 | 1486 | 477.136 | 515.150 |
| Takwa | Y2002/03 | M | 448.935 | 6.547 | 1486 | 436.092 | 461.777 |
| Takwa | Y2015 | M | 433.254 | 4.159 | 1486 | 425.097 | 441.412 |
| Takwa | Y2017 | M | 433.918 | 6.343 | 1486 | 421.475 | 446.361 |

*Table S3 Total length contrasts. Where insufficient data existed to make a contrast, NA is shown.*

| **River** | **Contrast** | **Sex** | **estimate** | **SE** | **df** | ***t.ratio*** | ***P*** |
|---|---|---|---|---|---|---|---|
| Chalifour | Y2002/03 - Y2015 | F | 74.175 | 18.971 | 1486 | 3.910 | 0.003 |
| Chalifour | Y2002/03 - Y2016 | F | 33.475 | 16.943 | 1486 | 1.976 | 0.232 |
| Chalifour | Y2002/03 - Y2017 | F | 45.722 | 19.580 | 1486 | 2.335 | 0.157 |
| Chalifour | Y2015 - Y2016 | F | -40.700 | 14.451 | 1486 | -2.816 | 0.067 |
| Chalifour | Y2015 - Y2017 | F | -28.452 | 17.468 | 1486 | -1.629 | 0.379 |
| Chalifour | Y2016 - Y2017 | F | 12.247 | 15.241 | 1486 | 0.804 | 0.954 |
| Chalifour | Y2002/03 - Y2015 | M | 48.530 | 6.064 | 1486 | 8.003 | 0.000 |
| Chalifour | Y2002/03 - Y2016 | M | 35.505 | 5.904 | 1486 | 6.014 | 0.000 |
| Chalifour | Y2002/03 - Y2017 | M | 54.778 | 6.360 | 1486 | 8.612 | 0.000 |
| Chalifour | Y2015 - Y2016 | M | -13.025 | 5.638 | 1486 | -2.310 | 0.108 |
| Chalifour | Y2015 - Y2017 | M | 6.248 | 6.115 | 1486 | 1.022 | 0.756 |
| Chalifour | Y2016 - Y2017 | M | 19.273 | 5.956 | 1486 | 3.236 | 0.009 |
| Icon | Y2002/03 - Y2015 | F | 113.908 | 17.669 | 1486 | 6.447 | 0.000 |
| Icon | Y2002/03 - Y2016 | F | 81.341 | 14.744 | 1486 | 5.517 | 0.000 |
| Icon | Y2002/03 - Y2017 | F | NA | NA | NA | NA | NA |
| Icon | Y2015 - Y2016 | F | -32.567 | 19.953 | 1486 | -1.632 | 0.577 |
| Icon | Y2015 - Y2017 | F | NA | NA | NA | NA | NA |
| Icon | Y2016 - Y2017 | F | NA | NA | NA | NA | NA |
| Icon | Y2002/03 - Y2015 | M | 65.636 | 7.974 | 1486 | 8.231 | 0.000 |
| Icon | Y2002/03 - Y2016 | M | 64.475 | 7.585 | 1486 | 8.500 | 0.000 |
| Icon | Y2002/03 - Y2017 | M | 82.057 | 9.834 | 1486 | 8.345 | 0.000 |
| Icon | Y2015 - Y2016 | M | -1.161 | 5.612 | 1486 | -0.207 | 1.000 |
| Icon | Y2015 - Y2017 | M | 16.421 | 8.406 | 1486 | 1.953 | 0.207 |
| Icon | Y2016 - Y2017 | M | 17.582 | 8.038 | 1486 | 2.187 | 0.163 |
| Perch | Y2002/03 - Y2015 | F | 57.815 | 14.316 | 1486 | 4.038 | 0.002 |
| Perch | Y2002/03 - Y2016 | F | 42.532 | 11.496 | 1486 | 3.700 | 0.004 |
| Perch | Y2002/03 - Y2017 | F | NA | NA | NA | NA | NA |
| Perch | Y2015 - Y2016 | F | -15.283 | 15.184 | 1486 | -1.007 | 1.000 |
| Perch | Y2015 - Y2017 | F | NA | NA | NA | NA | NA |
| Perch | Y2016 - Y2017 | F | NA | NA | NA | NA | NA |
| Perch | Y2002/03 - Y2015 | M | 8.193 | 8.952 | 1486 | 0.915 | 0.954 |
| Perch | Y2002/03 - Y2016 | M | 29.360 | 6.887 | 1486 | 4.263 | 0.001 |

| | | | | | | | |
|---|---|---|---|---|---|---|---|
| Perch | Y2002/03 - Y2017 | M | 22.232 | 13.655 | 1486 | 1.628 | 0.577 |
| Perch | Y2015 - Y2016 | M | 21.167 | 9.125 | 1486 | 2.320 | 0.207 |
| Perch | Y2015 - Y2017 | M | 14.039 | 14.910 | 1486 | 0.942 | 0.954 |
| Perch | Y2016 - Y2017 | M | -7.128 | 13.769 | 1486 | -0.518 | 1.000 |
| Takwa | Y2002/03 - Y2015 | F | -7.889 | 9.124 | 1486 | -0.865 | 1.000 |
| Takwa | Y2002/03 - Y2016 | F | NA | NA | NA | NA | NA |
| Takwa | Y2002/03 - Y2017 | F | 5.368 | 11.735 | 1486 | 0.457 | 1.000 |
| Takwa | Y2015 - Y2016 | F | NA | NA | NA | NA | NA |
| Takwa | Y2015 - Y2017 | F | 13.257 | 11.547 | 1486 | 1.148 | 1.000 |
| Takwa | Y2016 - Y2017 | F | NA | NA | NA | NA | NA |
| Takwa | Y2002/03 - Y2015 | M | 15.680 | 7.756 | 1486 | 2.022 | 0.694 |
| Takwa | Y2002/03 - Y2016 | M | NA | NA | NA | NA | NA |
| Takwa | Y2002/03 - Y2017 | M | 15.016 | 9.116 | 1486 | 1.647 | 0.756 |
| Takwa | Y2015 - Y2016 | M | NA | NA | NA | NA | NA |
| Takwa | Y2015 - Y2017 | M | -0.664 | 7.585 | 1486 | -0.088 | 1.000 |
| Takwa | Y2016 - Y2017 | M | NA | NA | NA | NA | NA |

| **contrast** | **River** | **estimate** | **SE** | **df** | **p.val.adjusted** |
|---|---|---|---|---|---|
| Y2002/03,F - Y2002/03,M | Chalifour | 40.778 | 15.45949 | 1486 | 0.095 |
| Y2015,F - Y2015,M | Chalifour | 15.133 | 12.55743 | 1486 | 1.000 |
| Y2016,F - Y2016,M | Chalifour | 42.808 | 9.106448 | 1486 | 0.000 |
| Y2017,F - Y2017,M | Chalifour | 49.833 | 13.59589 | 1486 | 0.005 |
| Y2002/03 F - Y2002/03 M | Icon | 90.397 | 10.51359 | 1486 | 0.000 |
| Y2015 F - Y2015 M | Icon | 42.125 | 16.28627 | 1486 | 0.114 |
| Y2016 F - Y2016 M | Icon | 73.531 | 12.82151 | 1486 | 0.000 |
| Y2017 F - Y2017 M | Icon | NA | NA | NA | NA |
| Y2002/03 F - Y2002/03 M | Perch | 76.993 | 8.685832 | 1486 | 0.000 |
| Y2015 F - Y2015 M | Perch | 27.371 | 14.47971 | 1486 | 0.587 |
| Y2016 F - Y2016 M | Perch | 63.822 | 10.20524 | 1486 | 0.000 |
| Y2017 F - Y2017 M | Perch | NA | NA | NA | NA |
| Y2002/03 F - Y2002/03 M | Takwa | 52.576 | 9.310169 | 1486 | 0.000 |
| Y2015 F - Y2015 M | Takwa | 76.146 | 7.531934 | 1486 | 0.000 |
| Y2016 F - Y2016 M | Takwa | NA | NA | NA | NA |
| Y2017 F - Y2017 M | Takwa | 62.224 | 11.58146 | 1486 | 0.000 |

*Table S4 Mass lsmean with lower (LCI) and upper (UCI) 95% confidence interval*

| **River** | **Year** | **Sex** | **lsmean** | **SE** | **df** | **LCI** | **UCI** |
|---|---|---|---|---|---|---|---|
| Chalifour | Y2002/03 | F | 1083.781 | 1.118 | 1481 | 871.040 | 1348.482 |
| Chalifour | Y2015 | F | 635.239 | 1.093 | 1481 | 533.144 | 756.884 |
| Chalifour | Y2016 | F | 834.081 | 1.064 | 1481 | 738.480 | 942.058 |
| Chalifour | Y2017 | F | 766.353 | 1.101 | 1481 | 634.220 | 926.015 |

| | | | | | | | |
|---|---|---|---|---|---|---|---|
| Chalifour | Y2002/03 | M | 804.122 | 1.034 | 1481 | 753.097 | 858.604 |
| Chalifour | Y2015 | M | 609.089 | 1.031 | 1481 | 573.270 | 647.146 |
| Chalifour | Y2016 | M | 632.621 | 1.030 | 1481 | 597.534 | 669.768 |
| Chalifour | Y2017 | M | 559.218 | 1.035 | 1481 | 523.025 | 597.914 |
| Icon | Y2002/03 | F | 1316.329 | 1.066 | 1481 | 1160.305 | 1493.335 |
| Icon | Y2015 | F | 677.842 | 1.125 | 1481 | 537.611 | 854.651 |
| Icon | Y2016 | F | 908.121 | 1.097 | 1481 | 757.145 | 1089.202 |
| Icon | Y2017 | F | NA | NA | 1481 | NA | NA |
| Icon | Y2002/03 | M | 778.540 | 1.052 | 1481 | 705.276 | 859.414 |
| Icon | Y2015 | M | 461.328 | 1.033 | 1481 | 432.737 | 491.807 |
| Icon | Y2016 | M | 469.495 | 1.027 | 1481 | 445.413 | 494.879 |
| Icon | Y2017 | M | 413.395 | 1.056 | 1481 | 371.688 | 459.781 |
| Perch | Y2002/03 | F | 1066.553 | 1.056 | 1481 | 957.583 | 1187.924 |
| Perch | Y2015 | F | 695.730 | 1.097 | 1481 | 580.064 | 834.460 |
| Perch | Y2016 | F | 750.420 | 1.069 | 1481 | 658.207 | 855.552 |
| Perch | Y2017 | F | NA | NA | 1481 | NA | NA |
| Perch | Y2002/03 | M | 579.106 | 1.036 | 1481 | 540.230 | 620.779 |
| Perch | Y2015 | M | 568.516 | 1.059 | 1481 | 508.060 | 636.166 |
| Perch | Y2016 | M | 438.469 | 1.039 | 1481 | 407.101 | 472.254 |
| Perch | Y2017 | M | 479.248 | 1.101 | 1481 | 396.617 | 579.095 |
| Takwa | Y2002/03 | F | 974.184 | 1.051 | 1481 | 883.485 | 1074.196 |
| Takwa | Y2015 | F | 1043.415 | 1.048 | 1481 | 951.027 | 1144.778 |
| Takwa | Y2016 | F | NA | NA | 1481 | NA | NA |
| Takwa | Y2017 | F | 959.763 | 1.078 | 1481 | 828.898 | 1111.290 |
| Takwa | Y2002/03 | M | 699.694 | 1.051 | 1481 | 635.229 | 770.702 |
| Takwa | Y2015 | M | 657.593 | 1.032 | 1481 | 618.431 | 699.234 |
| Takwa | Y2016 | M | NA | NA | 1481 | NA | NA |
| Takwa | Y2017 | M | 716.593 | 1.050 | 1481 | 651.243 | 788.501 |

*Table S5 Mass contrasts. Where insufficient data existed to make a contrast, NA is indicated.*

| River | contrast | Sex | River | estimate | SE | df | t.ratio | p.val.adjusted |
|---|---|---|---|---|---|---|---|---|
| Chalifour | Y2002/03 - Y2015 | F | Chalifour | 0.534 | 0.143 | 1481 | 3.741 | 0.006 |
| Chalifour | Y2002/03 - Y2016 | F | Chalifour | 0.262 | 0.128 | 1481 | 2.054 | 0.214 |
| Chalifour | Y2002/03 - Y2017 | F | Chalifour | 0.347 | 0.147 | 1481 | 2.352 | 0.151 |
| Chalifour | Y2015 - Y2016 | F | Chalifour | -0.272 | 0.109 | 1481 | -2.504 | 0.151 |
| Chalifour | Y2015 - Y2017 | F | Chalifour | -0.188 | 0.131 | 1481 | -1.427 | 0.513 |
| Chalifour | Y2016 - Y2017 | F | Chalifour | 0.085 | 0.115 | 1481 | 0.738 | 1 |
| Chalifour | Y2002/03 - Y2015 | M | Chalifour | 0.278 | 0.046 | 1481 | 6.103 | 0 |
| Chalifour | Y2002/03 - Y2016 | M | Chalifour | 0.24 | 0.044 | 1481 | 5.414 | 0 |
| Chalifour | Y2002/03 - Y2017 | M | Chalifour | 0.363 | 0.048 | 1481 | 7.606 | 0 |
| Chalifour | Y2015 - Y2016 | M | Chalifour | -0.038 | 0.042 | 1481 | -0.893 | 0.952 |
| Chalifour | Y2015 - Y2017 | M | Chalifour | 0.085 | 0.046 | 1481 | 1.856 | 0.257 |
| Chalifour | Y2016 - Y2017 | M | Chalifour | 0.123 | 0.045 | 1481 | 2.751 | 0.041 |
| Icon | Y2002/03 - Y2015 | F | Icon | 0.664 | 0.135 | 1481 | 4.933 | 0 |

| | | | | | | | | |
|---|---|---|---|---|---|---|---|---|
| Icon | Y2002/03 - Y2016 | F | Icon | 0.371 | 0.113 | 1481 | 3.29 | 0.015 |
| Icon | Y2002/03 - Y2017 | F | Icon | NA | NA | NA | NA | NA |
| Icon | Y2015 - Y2016 | F | Icon | -0.292 | 0.15 | 1481 | -1.947 | 0.315 |
| Icon | Y2015 - Y2017 | F | Icon | NA | NA | NA | NA | NA |
| Icon | Y2016 - Y2017 | F | Icon | NA | NA | NA | NA | NA |
| Icon | Y2002/03 - Y2015 | M | Icon | 0.523 | 0.06 | 1481 | 8.719 | 0 |
| Icon | Y2002/03 - Y2016 | M | Icon | 0.506 | 0.057 | 1481 | 8.859 | 0 |
| Icon | Y2002/03 - Y2017 | M | Icon | 0.633 | 0.074 | 1481 | 8.553 | 0 |
| Icon | Y2015 - Y2016 | M | Icon | -0.018 | 0.042 | 1481 | -0.415 | 1 |
| Icon | Y2015 - Y2017 | M | Icon | 0.11 | 0.063 | 1481 | 1.734 | 0.315 |
| Icon | Y2016 - Y2017 | M | Icon | 0.127 | 0.06 | 1481 | 2.103 | 0.201 |
| Perch | Y2002/03 - Y2015 | F | Perch | 0.427 | 0.108 | 1481 | 3.965 | 0.001 |
| Perch | Y2002/03 - Y2016 | F | Perch | 0.352 | 0.087 | 1481 | 4.063 | 0.001 |
| Perch | Y2002/03 - Y2017 | F | Perch | NA | NA | NA | NA | NA |
| Perch | Y2015 - Y2016 | F | Perch | -0.076 | 0.114 | 1481 | -0.662 | 1 |
| Perch | Y2015 - Y2017 | F | Perch | NA | NA | NA | NA | NA |
| Perch | Y2016 - Y2017 | F | Perch | NA | NA | NA | NA | NA |
| Perch | Y2002/03 - Y2015 | M | Perch | 0.018 | 0.067 | 1481 | 0.274 | 1 |
| Perch | Y2002/03 - Y2016 | M | Perch | 0.278 | 0.052 | 1481 | 5.367 | 0 |
| Perch | Y2002/03 - Y2017 | M | Perch | 0.189 | 0.103 | 1481 | 1.842 | 0.382 |
| Perch | Y2015 - Y2016 | M | Perch | 0.26 | 0.069 | 1481 | 3.782 | 0.003 |
| Perch | Y2015 - Y2017 | M | Perch | 0.171 | 0.112 | 1481 | 1.522 | 0.513 |
| Perch | Y2016 - Y2017 | M | Perch | -0.089 | 0.104 | 1481 | -0.858 | 1 |
| Takwa | Y2002/03 - Y2015 | F | Takwa | -0.069 | 0.069 | 1481 | -1 | 1 |
| Takwa | Y2002/03 - Y2016 | F | Takwa | NA | NA | NA | NA | NA |
| Takwa | Y2002/03 - Y2017 | F | Takwa | 0.015 | 0.09 | 1481 | 0.166 | 1 |
| Takwa | Y2015 - Y2016 | F | Takwa | NA | NA | NA | NA | NA |
| Takwa | Y2015 - Y2017 | F | Takwa | 0.084 | 0.088 | 1481 | 0.945 | 1 |
| Takwa | Y2016 - Y2017 | F | Takwa | NA | NA | NA | NA | NA |
| Takwa | Y2002/03 - Y2015 | M | Takwa | 0.062 | 0.058 | 1481 | 1.063 | 1 |
| Takwa | Y2002/03 - Y2016 | M | Takwa | NA | NA | NA | NA | NA |
| Takwa | Y2002/03 - Y2017 | M | Takwa | -0.024 | 0.069 | 1481 | -0.344 | 1 |
| Takwa | Y2015 - Y2016 | M | Takwa | NA | NA | NA | NA | NA |
| Takwa | Y2015 - Y2017 | M | Takwa | -0.086 | 0.058 | 1481 | -1.483 | 1 |
| Takwa | Y2016 - Y2017 | M | Takwa | NA | NA | NA | NA | NA |

| **contrast** | **River** | **estimate** | **SE** | **df** | **p.val.adjusted** |
|---|---|---|---|---|---|
| Y2002/03,F - Y2002/03,M | Chalifour | 0.298 | 0.116 | 1481 | 0.130 |
| Y2015,F - Y2015,M | Chalifour | 0.042 | 0.095 | 1481 | 1.000 |
| Y2016,F - Y2016,M | Chalifour | 0.276 | 0.069 | 1481 | 0.002 |
| Y2017,F - Y2017,M | Chalifour | 0.315 | 0.102 | 1481 | 0.034 |
| Y2002/03 F - Y2002/03 M | Icon | 0.525 | 0.082 | 1481 | 0.000 |
| Y2015 F - Y2015 M | Icon | 0.385 | 0.123 | 1481 | 0.025 |
| Y2016 F - Y2016 M | Icon | 0.660 | 0.097 | 1481 | 0.000 |
| Y2017 F - Y2017 M | Icon | NA | NA | NA | NA |
| Y2002/03 F - Y2002/03 M | Perch | 0.611 | 0.065 | 1481 | 0.000 |
| Y2015 F - Y2015 M | Perch | 0.202 | 0.109 | 1481 | 0.577 |

| | | | | | |
|---|---|---|---|---|---|
| Y2016 F - Y2016 M | Perch | 0.537 | 0.077 | 1481 | 0.000 |
| Y2017 F - Y2017 M | Perch | NA | NA | NA | NA |
| Y2002/03 F - Y2002/03 M | Takwa | 0.331 | 0.070 | 1481 | 0.000 |
| Y2015 F - Y2015 M | Takwa | 0.462 | 0.057 | 1481 | 0.000 |
| Y2016 F - Y2016 M | Takwa | NA | NA | NA | NA |
| Y2017 F - Y2017 M | Takwa | 0.292 | 0.089 | 1481 | 0.025 |

<u>Sex Bias</u>

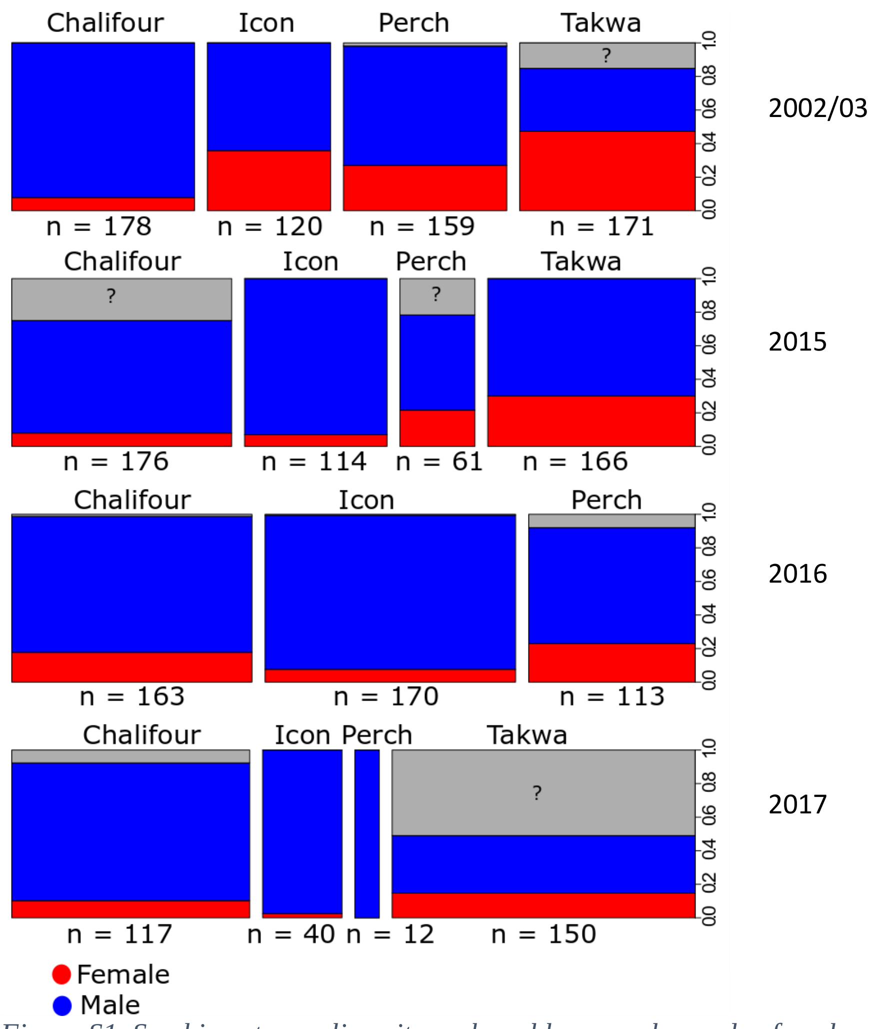

*Figure S1  Sex-bias at sampling sites, where blue = males, red = females and grey = unknowns in 2002-03, 2015, 2016 and 2017. Also shown are the total captured walleye in each river (n).*

*Genomics*

*Table S6 Number of individuals successfully sequenced and retained after all filtering (n), number of private alleles (private), number of SNPs (SNPs), percent polymorphic loci (% poly), expected heterozygosity (HE), observed heterozygosity (HO), pi (π).*

| Population & year | n | Private | SNPs | % poly | $H_E$ | StdErr | $H_O$ | StdErr | π | StdErr |
|---|---|---|---|---|---|---|---|---|---|---|
| **Chalifour 2003** | 27.5725 | 12 | 7995 | 0.1792 | 0.2302 | 0.0019 | 0.2256 | 0.002 | 0.2345 | 0.0019 |
| **Chalifour 2015** | 46.8723 | 14 | 9236 | 0.1961 | 0.2264 | 0.0018 | 0.2251 | 0.0019 | 0.2289 | 0.0018 |
| **Icon-Perch 2003** | 83.9165 | 53 | 7028 | 0.1838 | 0.2134 | 0.002 | 0.2014 | 0.0019 | 0.2146 | 0.002 |
| **Icon-Perch 2015** | 85.3662 | 44 | 9196 | 0.2017 | 0.2183 | 0.0018 | 0.2099 | 0.0018 | 0.2196 | 0.0018 |
| **Takwa 2003** | 33.3336 | 51 | 9253 | 0.1868 | 0.2223 | 0.0018 | 0.2141 | 0.0018 | 0.2257 | 0.0018 |
| **Takwa 2015** | 35.9956 | 7 | 8031 | 0.1754 | 0.2113 | 0.0019 | 0.2011 | 0.0019 | 0.2143 | 0.002 |

Choice of Outlier analysis programs

For historical-contemporary comparisons, we did not use OutFlank because only two "populations" were sampled, and the authors do not recommend it's use in these situations (Whitlock & Lotterhos 2015)

*Table S7 Outlier loci, PC axis on which they are relevant, and explanation of the population structure explained by each PC axis for historic-contemporary contrasts for each population separately and all populations combined. Figures S2 to S16 illustrate the results described in this table.*

| Population | Num. of individuals in each analysis: all (unknowns plus m & f) and for each sex separately | Num. of SNPs used in contrast | Num. of outliers | % of outliers (NS for no population structure) | Population structure explained by each PC when outliers are included in the data | Num. of outlier loci associated with relevant PC/total no. of outliers |
|---|---|---|---|---|---|---|
| **All populations and years, global** | all, 342 | 8728 | 70 | 0.80 | PC1 separates north from the south, and also Pe03 and Ch03/15 from Ic03/15/Pe15 marginally; PC2 separates Ch03/15 from the other populations, and also Pe03 from the other individuals. Outliers do not maintain the population structure. | PC1, 55/70; PC2, 15/70 |
| | f, 118 | 6613 | 11 | 0.17 | PC1 separates north from the south, and also Pe03 and Ch03/15 from Ic03/15/Pe15 marginally; PC2 separates Ch03 from Ch15, and both Chalifour years from the other southern populations. Outliers do not maintain the population structure. | PC1, 9/11; PC2, 2/11 |

| | | | | | | |
|---|---|---|---|---|---|---|
| | m, 213 | 9958 | 151 | 1.52 | PC1 separates north from the south, and also Pe03 and Ch03/15 from Ic03/15/Pe15 marginally; PC2 separates Ch03/15 from the other populations, Ch03 from Ch15, and also Pe03 from the other individuals. Outliers maintain population structure in Ch. | PC1, 91/151; PC2, 60/151 |
| **south** | all, 268 | 7358 | 51 | 0.69 | PCI separates Ch from other pops, PC2 separates Ch 03 from 15 and Pe03. Outliers maintain population structure in Ch. | PC1, 15/51; PC2, 36/51 |
| | f, 83 | 6547 | 65 | 0.99 | PC1 separates Ch03/15 from the other sites. PC2 separates Pe03 and Cha03 from the other sites. Outliers maintain population structure. | PC1, 17/65; PC2, 48/65 |
| | m, 176 | 8068 | 69 | 0.86 | PC1 separates Ch03/15 from other sites. PC2 separates Chali03 from 2015 and Pe03 a bit from rest of that group. Outliers maintain population structure | PC1, 18/69; PC2, 51/69 |
| **Chalifour** | all, 79 | 8513 | 129 | 1.52 | PC1, separates years. Outliers maintain population structure. | PC1, 109/129; PC2, 20/129 |
| | f, 17 | 3665 | 25 | 0.68 | PC1, separates years. Outliers do not maintain population structure | PC1, 16/25; PC2, 9/25 |
| | m, 61 | 9294 | 263 | 2.83 | PC1, separates years. Outliers maintain population structure. | PC1, 233/263; PC2, 30/263 |
| **Icon-Perch metapopulation** | all, 189 | 6802 | 33 | 0.49 | PC2 separates Pe03 from the other populations. Outliers maintain population structure | PC2, 8/33 |
| | f, 66 | 6442 | 73 | 1.13 | PC1 separates Pe03 from the other populations. Outliers maintain population structure. | PC1, 63/73 outliers |
| | m, 115 | 6452 | NS | NS | NS | NS |
| **Takwa** | all, 74 | 7820 | NS | NS | NS | NS |
| | f, 35 | 6543 | NS | NS | NS | NS |
| | m, 37 | 10540 | NS | NS | NS | NS |

**^ loci were not required to be present in all populations to be used (filter requirements were the same as for the main dataset**

*Table S8 Outlier loci blasted, mapped and annotated using blast2go. Annotation for gene ontology level 2. Biological process (BP), Molecular function (MF) and Cellular component (CC).*

| **Popul-ation** | **Sex** | **Number of sequences blasted** | **Number of alleles mapped** | **Number of alleles annotated** | **Annotation (Gene Ontology = 2)** | | |
|---|---|---|---|---|---|---|---|
| | | | | | **BP** | **MF** | **CC** |
| Cha | all | 253 | 3 | 2 | Localization | binding | cell part, organelle part, organelle, cell, membrane, protein containing complex |

| | | | | | | | |
|---|---|---|---|---|---|---|---|
| Cha | f | 46 | 0 | 4 | biological regulation, signaling, response to stimulus, regulation of biological process, cellular process, metabolic process, localization, negative regulation of biological process, positive regulation of biological process, cellular component organisation or biogenesis, multicellular organismal process | binding, transporter activity, catalytic activity | organelle, cell, cell part, synapse, cell junction, protein-containing complex, synapse part, membrane-enclosed lumen, organelle part, membrane part, membrane |
| Cha | m | 513 | 7 | 6 | biological regulation, signaling, localization, response to stimulus, regulation of biological process, negative regulation of biological process, cellular component organisation of biogenesis, developmental process, multicellular organismal process, cellular process, cell aggregation, biological adhesion, behavior, metabolic process, positive regulation of biological process, growth, multi-organismal process | binding, molecular transducer activity, structural molecule activity, transporter activity | cell, protein-containing complex, cell part, organelle, organelle part, membrane, synapse, supermolecular complex, cell junction, synapse part, extracellular region, extracellular region part, membrane part |
| IcPe | all | 64 | 0 | 2 | negative regulation of biological process, biological regulation, metabolic process, signaling, response to stimulus, cellular process, regulation of biological process | catalytic activity, binding | cell, cell part |
| IcPe | f | 144 | 0 | 2 | signaling, response to stimulus, regulation of biological process, growth, locomotion, developmental process, multicellular organismal process, cellular process, biological regulation, metabolic process, localization, reproductive process, negative regulation of biological process, positive regulation of biological process, cell population proliferation, multi-organism process, cellular component organization or biogenesis, reproduction | molecular transducer activity, transcription regulator activity, binding | organelle, cell, membrane-enclosed lumen, cell part, organelle part, protein-containing complex |
| Sou | all | 128 | 0 | 2 | signaling, response to stimulus, regulation of biological process, growth, locomotion, developmental process, multicellular organismal process, cellular process, biological regulation, metabolic process, localization, reproductive process, negative regulation of biological process, positive regulation of biological process, cell population proliferation, multi-organism process, cellular component organization of biogenesis, reproduction | molecular transducer activity, transcription regulator activity, binding | organelle, cell, membrane-enclosed lumen, cell part, organelle part, protein-containing complex |

Outlier analysis results presentation

For each configuration of populations assessed, results for m and f combined are followed by results for each sex separately. Meaningful population structure that allowed for outlier detection was present in most cases, sometimes based on geography and sometimes temporal between timepoints.

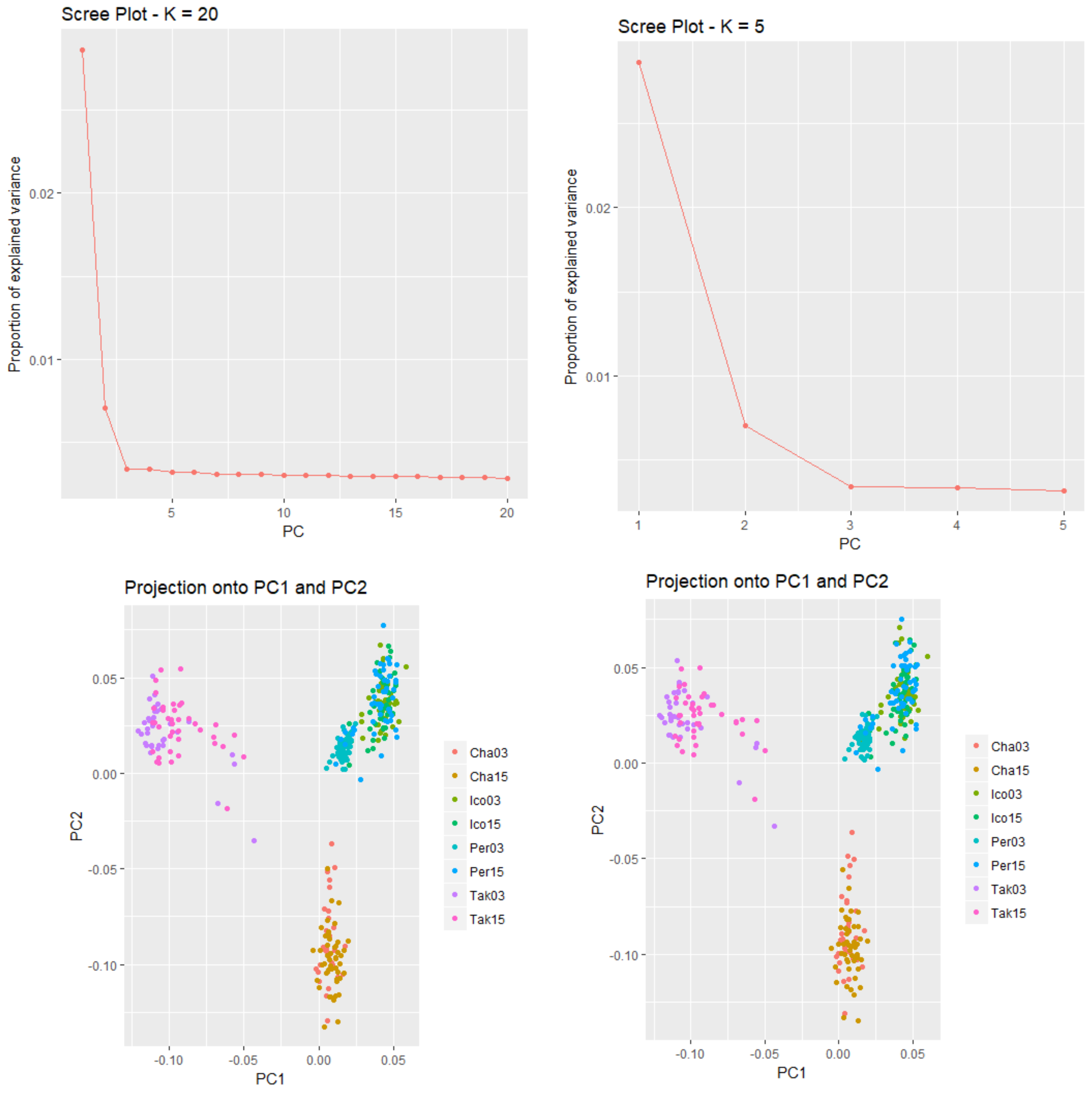

*Figure S2 All populations, all years, m and f combined, k=2. Top: scree panels for selecting best k. Bottom left, with outliers left in. Bottom right, excluding outliers. Result: PC1 separates the north from the south, and also Icon03/15Perch15 a bit, and the second PC separates Chalifour from the other pops. Shown in the SNP list that associates SNPs with PCs below, the vast majority of outliers are associated with PC1 (15 to PC2). However, removing outliers does not change the structure in a meaningful way.*

##overall outliers

```
> print(ID[outliers])
[1] "2764_47"   "5777_26"   "7221_52"   "7315_29"   "9664_69"   "10060_79"  "10881_53"
 [8] "11507_79"  "12647_55"  "14845_11"  "15362_57"  "15426_49"  "16571_28"  "23892_66"
[15] "25728_47"  "26281_46"  "27021_71"  "28192_35"  "29788_76"  "31425_7"   "31886_71"
[22] "34289_47"  "35235_60"  "36864_48"  "37605_43"  "38039_35"  "39685_62"  "41527_40"
[29] "41635_58"  "45518_67"  "45711_42"  "46331_65"  "48658_18"  "49504_49"  "50030_36"
[36] "50404_26"  "51264_20"  "53498_56"  "62465_26"  "65438_60"  "65770_56"  "69567_54"
[43] "69971_62"  "71554_26"  "72916_70"  "74122_76"  "74452_59"  "75343_34"  "76517_75"
[50] "81466_14"  "83593_61"  "84174_76"  "85630_70"  "88020_72"  "88250_71"  "89842_9"
[57] "91022_14"  "91315_70"  "91507_23"  "93560_78"  "93677_42"  "97401_34"  "97840_17"
[64] "99863_28"  "99888_60"  "101083_6"  "102928_19" "103420_43" "104231_8"  "104441_34"
```

=70 outlier loci --- note that did not require all loci to be present in all pops here, so can't really just divide by total number of SNPs to get the percent outliers. Average number of SNPs per pop was 7817, but ranged between 3766 and 8694.

```
> snp_pc <- get.pc(x, outliers)
> snp_pc
    SNP PC
1    99  1
2   341  1
3   456  1
4   468  2
5   671  2
6   708  1
7   779  1
8   832  1
9   944  1
10 1151  1
11 1190  1
12 1199  1
13 1292  2
14 1938  1
15 2122  1
16 2172  2
17 2254  2
18 2347  1
19 2487  1
20 2639  1
21 2684  1
22 2914  1
23 3010  1
24 3154  1
25 3227  1
26 3269  1
27 3408  1
28 3556  1
29 3562  1
30 3882  2
31 3900  1
```

| | | |
|---|---|---|
| 32 | 3956 | 1 |
| 33 | 4139 | 1 |
| 34 | 4204 | 1 |
| 35 | 4241 | 2 |
| 36 | 4265 | 1 |
| 37 | 4315 | 2 |
| 38 | 4444 | 2 |
| 39 | 4871 | 1 |
| 40 | 5035 | 1 |
| 41 | 5063 | 1 |
| 42 | 5301 | 1 |
| 43 | 5320 | 1 |
| 44 | 5464 | 1 |
| 45 | 5583 | 2 |
| 46 | 5699 | 1 |
| 47 | 5733 | 1 |
| 48 | 5813 | 2 |
| 49 | 5907 | 1 |
| 50 | 6398 | 2 |
| 51 | 6602 | 1 |
| 52 | 6662 | 1 |
| 53 | 6781 | 2 |
| 54 | 6974 | 1 |
| 55 | 7000 | 1 |
| 56 | 7139 | 1 |
| 57 | 7244 | 1 |
| 58 | 7269 | 1 |
| 59 | 7287 | 1 |
| 60 | 7498 | 1 |
| 61 | 7515 | 1 |
| 62 | 7867 | 1 |
| 63 | 7896 | 1 |
| 64 | 8078 | 2 |
| 65 | 8081 | 1 |
| 66 | 8190 | 1 |
| 67 | 8356 | 1 |
| 68 | 8405 | 2 |
| 69 | 8489 | 1 |
| 70 | 8511 | 1 |

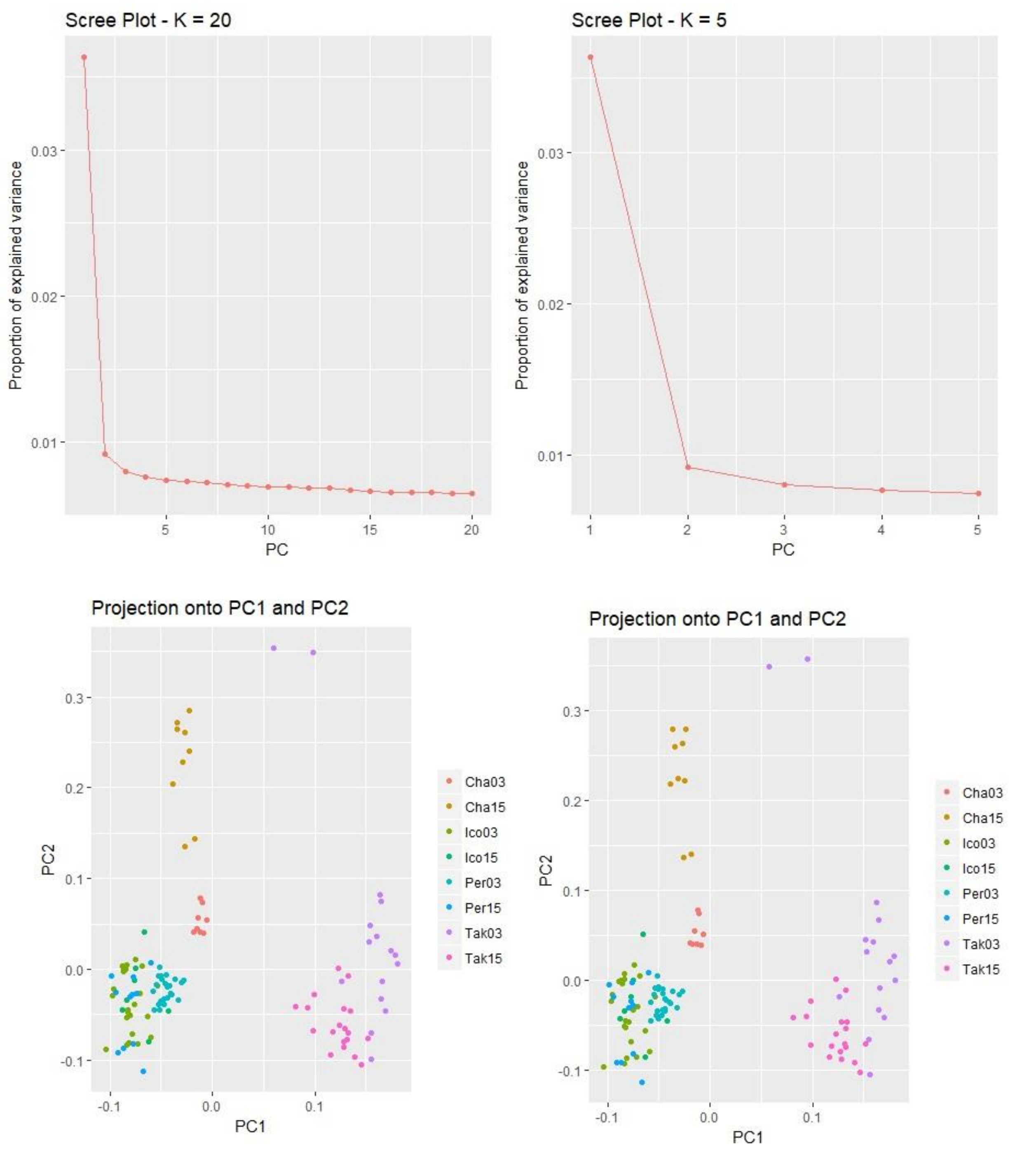

*Figure S3 All populations and years, f, k=2. Top panels are scree plots for determining the best k. Bottom panels are: (left) including outliers, and (right) excluding outliers. Result: PC1 separates north and south, and maybe Chali03/15 and Pe03 a little bit from the Ic03,15,Pe15 group. PC2 separates Ch03 from Ch15, and also both Chalifour years from the other southern pops. Result looks similar to when M and F are combined, except that PC2 also separates Ch03 and 15. Removing outliers has no meaningful affect on the population structure. number of pops that I required loci to be present in for the full dataset is 6/8.*

```
> print(ID[outliers])
[1] "2764_47"   "12315_60"  "13844_69"  "16571_28"  "27138_61"  "34289_47"  "36864_48"
 [8] "45711_42"  "89842_9"   "93560_78"  "103420_43"
```

=11/6613 SNPs, but not all SNPs would have been present in all pops

```
> snp_pc
   SNP PC
1    70  1
2   669  2
3   774  2
4   950  1
5  1696  1
6  2224  1
7  2410  1
8  2970  1
9  5472  1
10 5739  1
11 6434  1
```

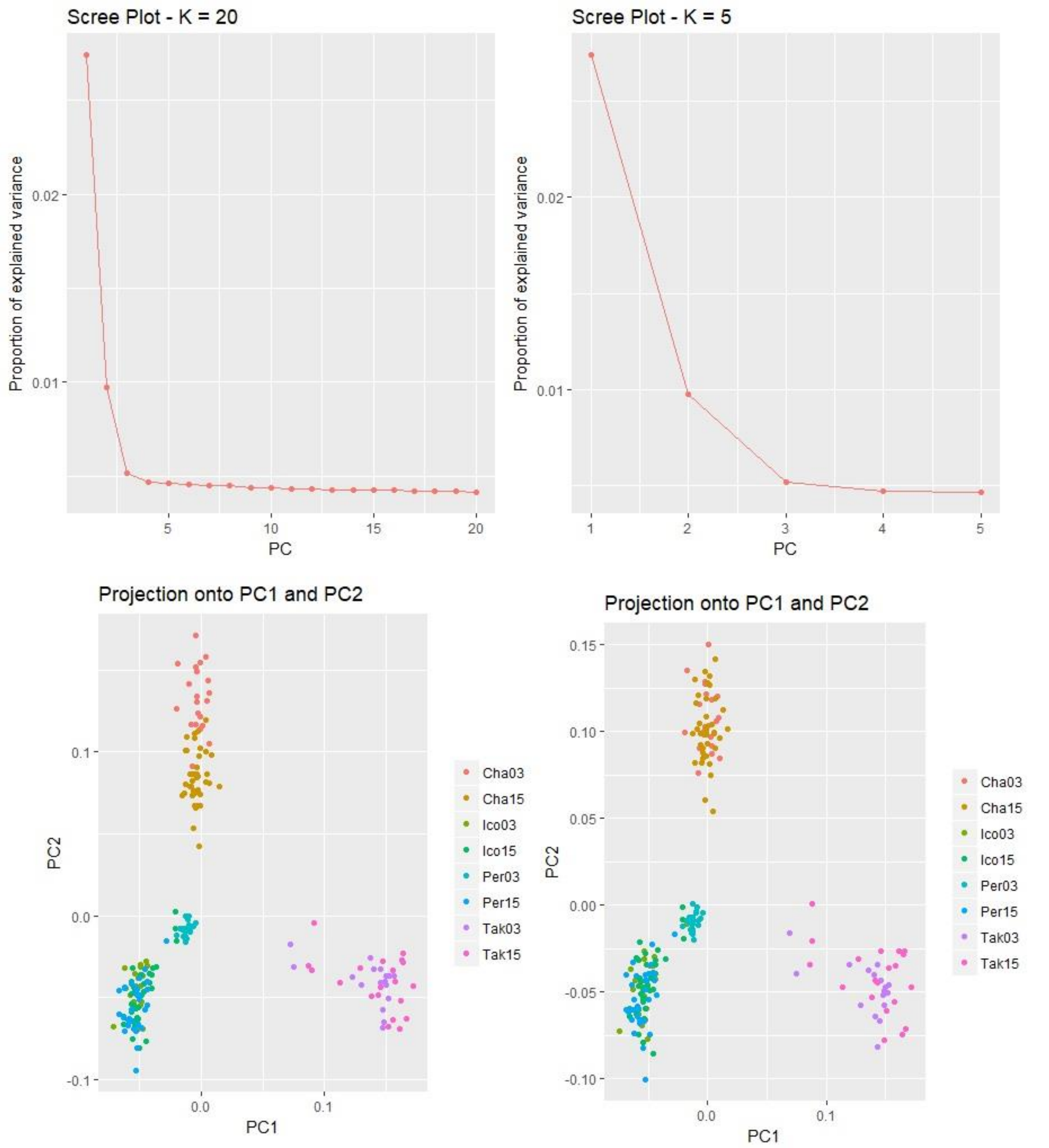

*Figure S4 All populations and years, m, k=2. Top panels are scree plots used to determine best k. Bottom panels are: (left) including outliers, and (right) excluding outliers. Result: PC1 separates north from south, and also Pe03 and the Chalifours. PC2 separates Cha03 and 15 from the other pops, Pe03 from the other pops, and Ch03 from 15. Removing outliers removes the structure between Ch03 and 15.*

```
> print(ID[outliers])
[1] "1721_78"   "2468_66"   "2764_47"   "2916_78"   "5486_58"   "5777_26"   "7221_52"
 [8] "7232_14"   "7393_79"   "8976_18"   "9415_78"   "9629_79"   "9664_69"   "9942_79"
```

```
 [15] "10060_79"  "10157_79"  "10696_78"  "10881_53"  "11507_79"  "12647_55"  "12994_48"
 [22] "14431_76"  "15362_57"  "15426_49"  "16571_28"  "18456_6"   "19262_77"  "19850_46"
 [29] "21844_79"  "22374_17"  "23892_66"  "23960_64"  "24841_61"  "25728_47"  "26281_46"
 [36] "26667_79"  "27021_71"  "28192_35"  "29164_79"  "29788_76"  "31135_79"  "31385_26"
 [43] "31886_71"  "32228_78"  "33113_74"  "33150_19"  "33608_78"  "34382_79"  "34803_77"
 [50] "34842_79"  "35235_60"  "35988_62"  "36864_48"  "37605_43"  "38039_35"  "38403_14"
 [57] "38443_39"  "39301_13"  "39685_62"  "41527_40"  "41635_58"  "41986_78"  "42743_77"
 [64] "43077_77"  "43351_78"  "44213_21"  "44786_79"  "45557_77"  "45606_14"  "45711_42"
 [71] "46331_65"  "47350_50"  "48658_18"  "49504_49"  "50030_36"  "50404_26"  "51264_20"
 [78] "53093_25"  "53498_56"  "54744_32"  "58417_15"  "59518_78"  "61672_14"  "62465_26"
 [85] "62534_37"  "62632_56"  "63988_78"  "65438_60"  "65770_56"  "66402_17"  "67781_28"
 [92] "69537_35"  "69567_54"  "71145_48"  "71554_26"  "71655_77"  "72447_18"  "72944_78"
 [99] "76517_75"  "77651_61"  "77767_20"  "78676_53"  "79047_76"  "79305_23"  "79365_74"
[106] "80664_78"  "81466_14"  "82281_25"  "82349_79"  "82679_76"  "83593_61"  "83684_34"
[113] "84174_76"  "84455_39"  "85630_70"  "86067_79"  "87113_79"  "87783_79"  "87972_79"
[120] "88020_72"  "88250_71"  "88935_12"  "89842_9"   "90459_77"  "91022_14"  "91315_70"
[127] "91507_23"  "91852_78"  "93560_78"  "93677_42"  "94056_12"  "94159_79"  "94696_71"
[134] "95326_42"  "96032_32"  "96940_76"  "97401_34"  "97840_17"  "99888_60"  "101049_10"
[141] "101083_6"  "101770_56" "102928_19" "103420_43" "103612_10" "104231_8"  "104441_34"
[148] "113216_68" "114365_30" "117698_63" "123135_44"
```

=151 outliers
#interesting that there are so many more outliers for males
#variable number of SNPs per population

```
> snp_pc
    SNP PC
1    31  2
2    95  1
3   118  1
4   135  2
5   366  1
6   401  1
7   537  1
8   539  1
9   560  2
10  695  2
11  743  2
12  765  2
13  769  2
14  803  2
15  813  1
16  824  2
17  878  2
18  896  1
19  958  1
20 1089  1
21 1123  1
22 1283  2
23 1381  1
24 1391  2
```

| | | |
|---|---|---|
| 25 | 1498 | 2 |
| 26 | 1679 | 1 |
| 27 | 1759 | 2 |
| 28 | 1831 | 1 |
| 29 | 2030 | 2 |
| 30 | 2090 | 2 |
| 31 | 2241 | 1 |
| 32 | 2251 | 2 |
| 33 | 2343 | 1 |
| 34 | 2434 | 1 |
| 35 | 2493 | 2 |
| 36 | 2548 | 2 |
| 37 | 2582 | 2 |
| 38 | 2685 | 1 |
| 39 | 2782 | 2 |
| 40 | 2842 | 1 |
| 41 | 2980 | 2 |
| 42 | 3005 | 1 |
| 43 | 3061 | 1 |
| 44 | 3096 | 2 |
| 45 | 3187 | 1 |
| 46 | 3192 | 1 |
| 47 | 3245 | 2 |
| 48 | 3325 | 2 |
| 49 | 3373 | 2 |
| 50 | 3377 | 2 |
| 51 | 3424 | 1 |
| 52 | 3507 | 1 |
| 53 | 3580 | 1 |
| 54 | 3661 | 1 |
| 55 | 3708 | 1 |
| 56 | 3736 | 1 |
| 57 | 3739 | 1 |
| 58 | 3824 | 1 |
| 59 | 3862 | 1 |
| 60 | 4028 | 1 |
| 61 | 4035 | 1 |
| 62 | 4067 | 2 |
| 63 | 4140 | 1 |
| 64 | 4171 | 2 |
| 65 | 4201 | 2 |
| 66 | 4279 | 1 |
| 67 | 4328 | 2 |
| 68 | 4410 | 2 |
| 69 | 4413 | 1 |
| 70 | 4423 | 1 |
| 71 | 4486 | 1 |
| 72 | 4580 | 1 |
| 73 | 4686 | 2 |
| 74 | 4754 | 1 |
| 75 | 4800 | 2 |

| | | |
|---|---|---|
| 76 | 4829 | 1 |
| 77 | 4893 | 2 |
| 78 | 5027 | 1 |
| 79 | 5054 | 2 |
| 80 | 5147 | 2 |
| 81 | 5357 | 1 |
| 82 | 5400 | 2 |
| 83 | 5511 | 1 |
| 84 | 5557 | 1 |
| 85 | 5563 | 1 |
| 86 | 5569 | 1 |
| 87 | 5656 | 2 |
| 88 | 5750 | 1 |
| 89 | 5782 | 1 |
| 90 | 5836 | 2 |
| 91 | 5926 | 1 |
| 92 | 6059 | 1 |
| 93 | 6061 | 1 |
| 94 | 6198 | 1 |
| 95 | 6245 | 1 |
| 96 | 6252 | 2 |
| 97 | 6332 | 2 |
| 98 | 6384 | 2 |
| 99 | 6747 | 1 |
| 100 | 6876 | 1 |
| 101 | 6885 | 1 |
| 102 | 6968 | 1 |
| 103 | 7009 | 2 |
| 104 | 7037 | 1 |
| 105 | 7047 | 1 |
| 106 | 7189 | 2 |
| 107 | 7286 | 2 |
| 108 | 7374 | 1 |
| 109 | 7387 | 2 |
| 110 | 7427 | 2 |
| 111 | 7510 | 1 |
| 112 | 7524 | 1 |
| 113 | 7574 | 1 |
| 114 | 7598 | 1 |
| 115 | 7705 | 2 |
| 116 | 7748 | 2 |
| 117 | 7841 | 2 |
| 118 | 7907 | 2 |
| 119 | 7927 | 2 |
| 120 | 7931 | 1 |
| 121 | 7954 | 1 |
| 122 | 8030 | 1 |
| 123 | 8114 | 1 |
| 124 | 8179 | 2 |
| 125 | 8228 | 1 |
| 126 | 8256 | 1 |

127 8276  1
128 8315  2
129 8512  1
130 8528  1
131 8576  1
132 8588  2
133 8647  1
134 8712  1
135 8782  1
136 8871  2
137 8924  1
138 8960  1
139 9159  1
140 9275  1
141 9281  1
142 9340  1
143 9460  2
144 9513  2
145 9543  1
146 9612  1
147 9636  2
148 9784  1
149 9804  1
150 9866  1
151 9914  1

#60 outliers are associated with PC2

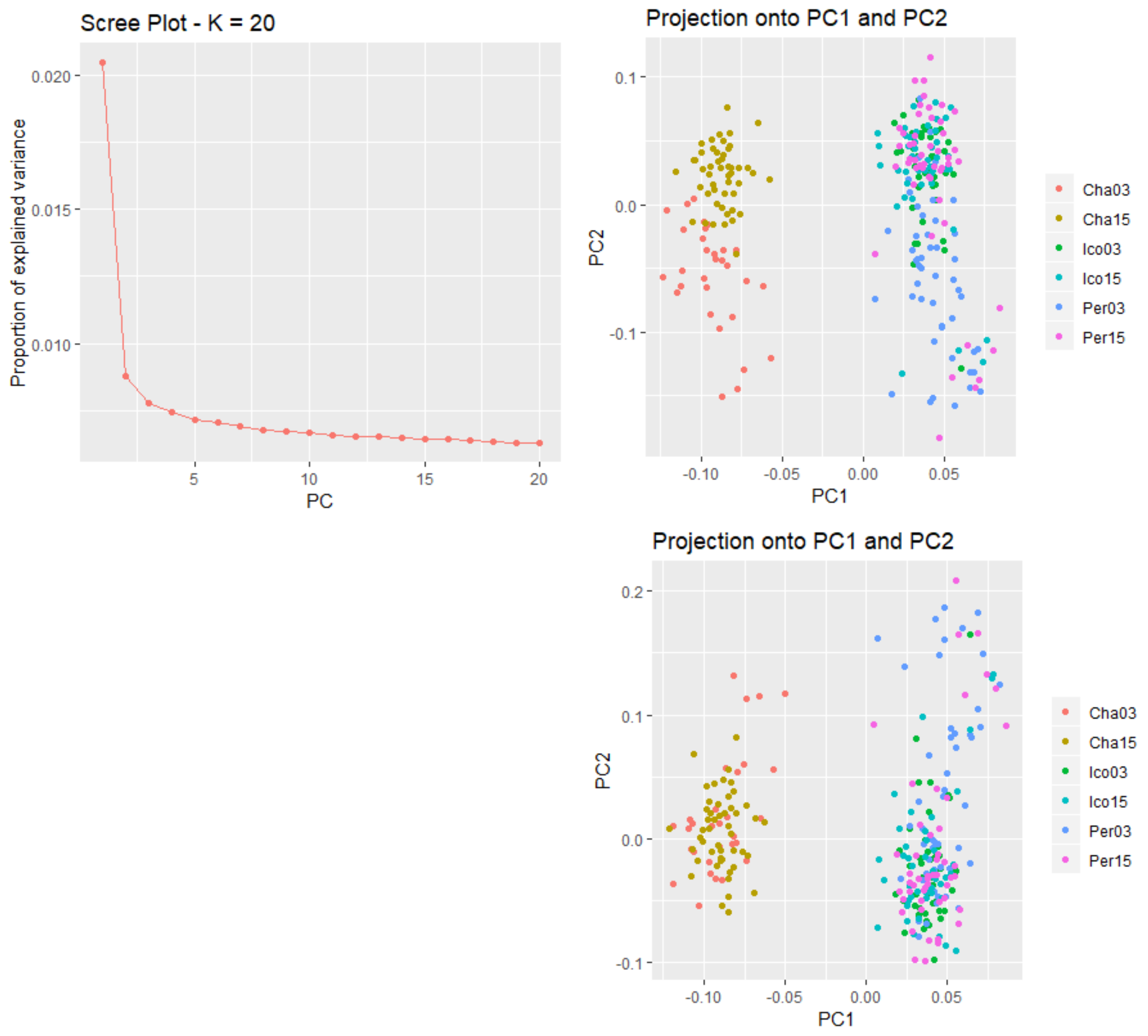

*Figure S5 Southern rivers, m & f combined, k=2. Top right panel is with outliers included and bottom right panel is outliers excluded. Conclusions: structure is sufficient to detect outliers. PCI separates Ch from other pops, PC2 separates Ch 03 from 15 and Pe03. Removing outliers removes the structure.*

[1] "866_78"  "3193_79"  "4975_78"  "5302_79"  "7583_78"  "8206_78"  "9415_78"  "9942_79"
[9] "10957_77" "11235_78" "16571_28" "23360_77" "24957_79" "27021_71" "28075_78" "29208_61"
[17] "29693_78" "31135_79" "34289_47" "34803_77" "34842_79" "35262_76" "36253_79" "37605_43"
[25] "42691_45" "47695_78" "48658_18" "49486_6"  "53498_56" "56508_40" "62534_37" "70726_79"
[33] "72944_78" "74122_76" "74436_48" "74568_79" "75276_78" "78450_6"  "83973_79" "84111_22"
[41] "87113_79" "88895_78" "89820_78" "92110_78" "96076_79" "99202_76" "101300_12" "102745_64"
[49] "103420_43" "104310_77" "104441_34"

```
> snp_pc
    SNP PC
1    10  2
2   127  2
3   231  2
4   250  2
5   413  2
6   457  2
7   542  2
8   585  2
9   665  2
10  684  2
11 1110  1
12 1617  2
13 1747  2
14 1928  1
15 1997  2
16 2076  2
17 2118  2
18 2242  2
19 2503  1
20 2552  1
21 2556  2
22 2596  2
23 2682  2
24 2782  1
25 3137  2
26 3486  2
27 3550  1
28 3612  2
29 3815  1
30 3948  1
31 4172  1
32 4584  2
33 4774  2
34 4865  1
35 4894  1
36 4902  2
37 4957  2
38 5210  2
39 5687  2
40 5694  2
41 5902  2
42 6051  2
43 6119  2
44 6305  1
45 6664  2
46 6896  2
47 7053  1
48 7159  2
49 7217  1
```

50 7296  2
51 7308  1

Pc1, 15/51

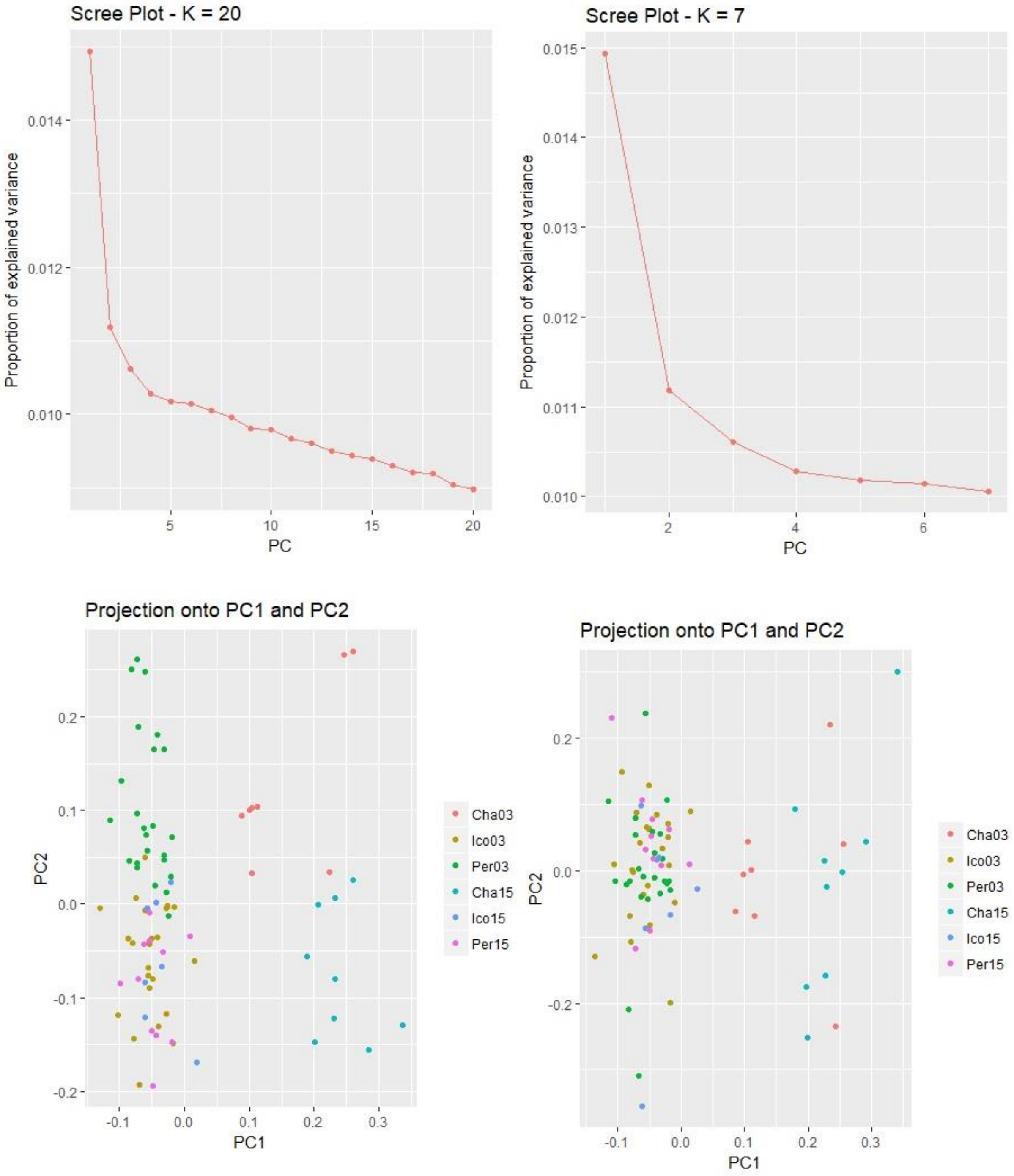

*Figure S6 southern rivers, f, k=2. Top panels are the scree plots used to find the best k. Bottom panels are: (left) including outliers and (right) excluding outliers. Result: PC1 separates Chalifour from the other sites and PC2 separates Pe03 and Cha03 from the other sites. Removing outliers collapses PC2*

*structure, collapsing Pe03 into the other southern rivers, and Ch03 and 15. This result is the inverse of the result when both sexes are analyzed together.*

```
> print(ID[outliers])
 [1] "2003_6"    "2179_39"   "3193_79"   "4975_78"   "6298_8"    "7025_29"   "7583_78"
 [8] "8206_79"   "9533_59"   "10957_77"  "11815_79"  "16571_28"  "17580_8"   "18297_78"
[15] "20589_78"  "22042_79"  "23360_77"  "23519_75"  "24957_79"  "25319_5"   "27138_61"
[22] "28075_78"  "28650_42"  "29693_78"  "31509_74"  "32700_77"  "32789_78"  "33821_6"
[29] "34289_47"  "34803_77"  "34842_79"  "35262_76"  "35866_16"  "41258_78"  "46439_76"
[36] "46573_78"  "46592_79"  "47695_78"  "48338_67"  "48757_78"  "50171_79"  "53917_60"
[43] "55665_78"  "57534_66"  "62150_77"  "70726_79"  "71278_78"  "74216_22"  "74568_79"
[50] "75276_78"  "77273_78"  "79332_79"  "80680_79"  "83973_79"  "88895_78"  "91894_55"
[57] "92110_78"  "94277_78"  "95663_78"  "96076_79"  "96628_71"  "99202_76"  "99476_17"
[64] "102675_77" "104310_77"
```

=65/6547 SNPs = 0.99% of loci are outliers

```
> snp_pc
    SNP PC
1    37  1
2    47  1
3   110  2
4   200  2
5   271  1
6   308  1
7   346  2
8   385  2
9   464  1
10  555  2
11  611  2
12  938  1
13  995  1
14 1047  2
15 1206  2
16 1310  2
17 1398  2
18 1411  2
19 1524  2
20 1547  2
21 1694  1
22 1749  2
23 1787  1
24 1856  2
25 2002  2
26 2098  2
27 2106  1
28 2190  1
29 2217  1
30 2263  1
31 2266  2
32 2298  2
33 2337  2
```

| | | |
|---|---|---|
| 34 | 2679 | 2 |
| 35 | 2987 | 2 |
| 36 | 2989 | 2 |
| 37 | 2990 | 2 |
| 38 | 3070 | 2 |
| 39 | 3107 | 1 |
| 40 | 3133 | 2 |
| 41 | 3223 | 2 |
| 42 | 3383 | 1 |
| 43 | 3449 | 2 |
| 44 | 3515 | 1 |
| 45 | 3663 | 2 |
| 46 | 4036 | 2 |
| 47 | 4086 | 2 |
| 48 | 4305 | 1 |
| 49 | 4329 | 2 |
| 50 | 4369 | 2 |
| 51 | 4511 | 2 |
| 52 | 4662 | 2 |
| 53 | 4765 | 2 |
| 54 | 5024 | 2 |
| 55 | 5361 | 2 |
| 56 | 5573 | 2 |
| 57 | 5595 | 2 |
| 58 | 5782 | 2 |
| 59 | 5892 | 2 |
| 60 | 5923 | 2 |
| 61 | 5959 | 2 |
| 62 | 6137 | 2 |
| 63 | 6154 | 2 |
| 64 | 6367 | 2 |
| 65 | 6495 | 2 |

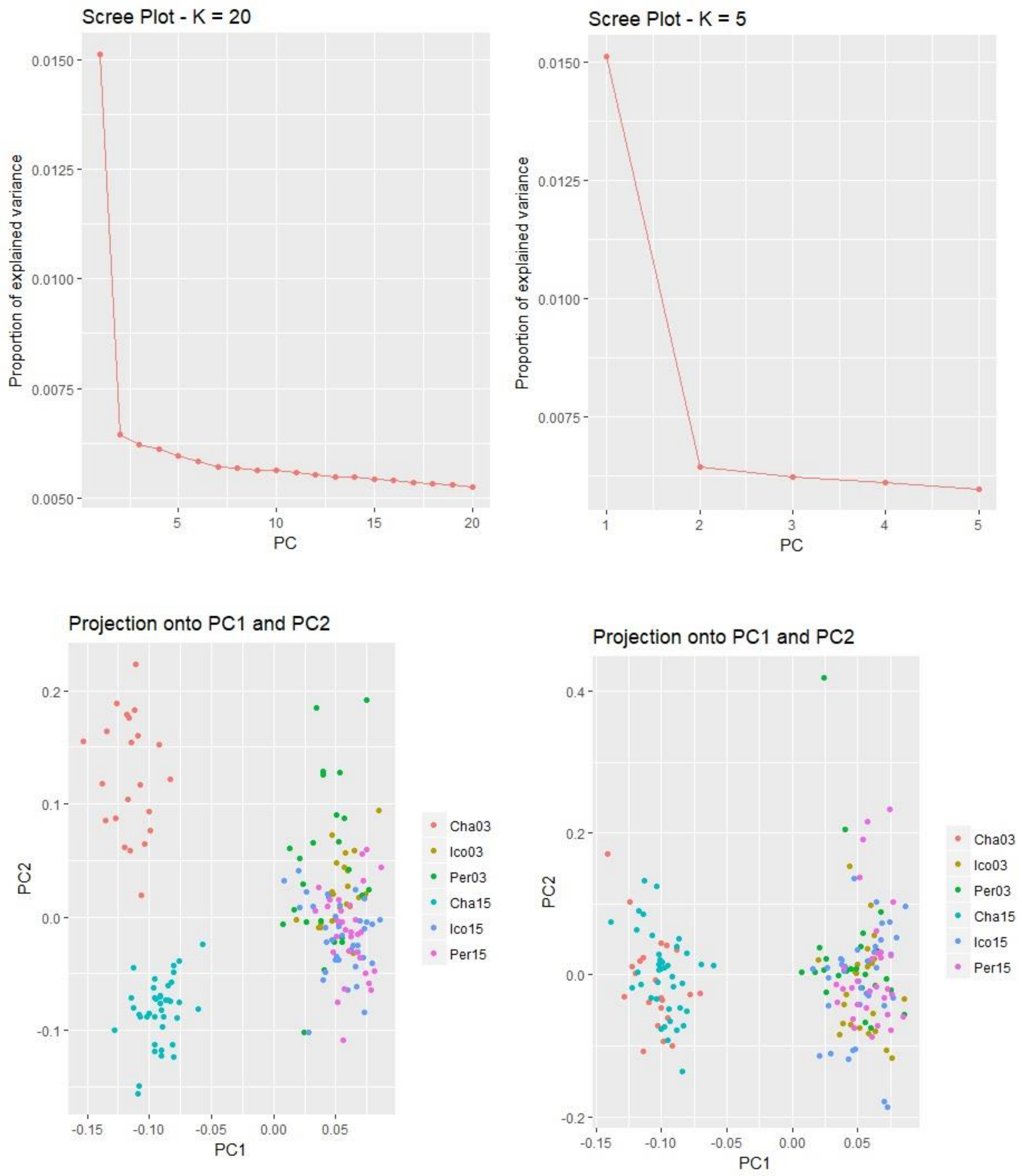

*Figure S7 south, m, k = 2. Top panels are scree plots used to select the best k. Bottom panels are: (left) including outliers, and (right) excluding outliers. Result: PC1 separates Chalifours from the other sites and Pe03 from Ic. PC2 seaprates Cha03 from 15. Removing outliers collapses Cha 03 and 15, and also collapses Pe03 completely into the other sites.*

```
> print(ID[outliers])
 [1] "866_78"   "1137_78"  "1745_76"  "1910_79"  "2364_79"  "3183_76"  "4431_19"
 [8] "4885_78"  "5302_79"  "7221_52"  "7583_78"  "9415_78"  "9664_69"  "9942_79"
[15] "10957_77" "11235_78" "13218_79" "13833_79" "14431_79" "16571_28" "18719_79"
```

```
[22] "20292_25"  "20378_78"  "21844_79"  "23629_79"  "23745_79"  "24957_79"  "25144_79"
[29] "26667_79"  "27021_71"  "31135_79"  "32440_79"  "34289_47"  "35231_79"  "37447_48"
[36] "37605_43"  "38039_35"  "39974_49"  "42257_74"  "46581_77"  "47695_78"  "48658_18"
[43] "50030_36"  "52351_78"  "53498_56"  "56508_40"  "62534_37"  "63490_78"  "70726_79"
[50] "71059_77"  "72944_78"  "73353_79"  "74122_76"  "74788_54"  "75276_78"  "75947_43"
[57] "80019_79"  "85905_79"  "87113_79"  "87783_79"  "89820_78"  "92110_79"  "95210_79"
[64] "95709_79"  "101352_79" "102737_78" "103420_43" "104310_77" "104441_34"
```
=69/8068 SNPs = 0.9% SNPs

PC2, 51 loci

```
> snp_pc
    SNP PC
1    10  2
2    12  2
3    24  2
4    35  2
5    62  2
6   135  2
7   219  2
8   255  2
9   281  2
10  433  1
11  468  2
12  609  1
13  630  1
14  657  2
15  752  2
16  772  2
17  947  2
18 1010  2
19 1062  2
20 1238  1
21 1414  2
22 1552  1
23 1556  2
24 1671  2
25 1823  2
26 1835  2
27 1943  2
28 1959  2
29 2110  2
30 2138  1
31 2467  2
32 2580  2
33 2748  1
34 2845  2
35 3030  2
36 3045  1
37 3086  1
38 3228  2
```

| | | |
|---|---|---|
| 39 | 3398 | 2 |
| 40 | 3737 | 2 |
| 41 | 3821 | 2 |
| 42 | 3888 | 1 |
| 43 | 3987 | 1 |
| 44 | 4125 | 2 |
| 45 | 4189 | 1 |
| 46 | 4344 | 1 |
| 47 | 4590 | 1 |
| 48 | 4650 | 1 |
| 49 | 5058 | 2 |
| 50 | 5092 | 2 |
| 51 | 5265 | 2 |
| 52 | 5301 | 2 |
| 53 | 5370 | 1 |
| 54 | 5422 | 2 |
| 55 | 5468 | 2 |
| 56 | 5526 | 2 |
| 57 | 5886 | 2 |
| 58 | 6406 | 2 |
| 59 | 6489 | 2 |
| 60 | 6548 | 2 |
| 61 | 6722 | 2 |
| 62 | 6916 | 2 |
| 63 | 7217 | 2 |
| 64 | 7270 | 2 |
| 65 | 7719 | 2 |
| 66 | 7831 | 2 |
| 67 | 7893 | 1 |
| 68 | 7979 | 2 |
| 69 | 7992 | 1 |

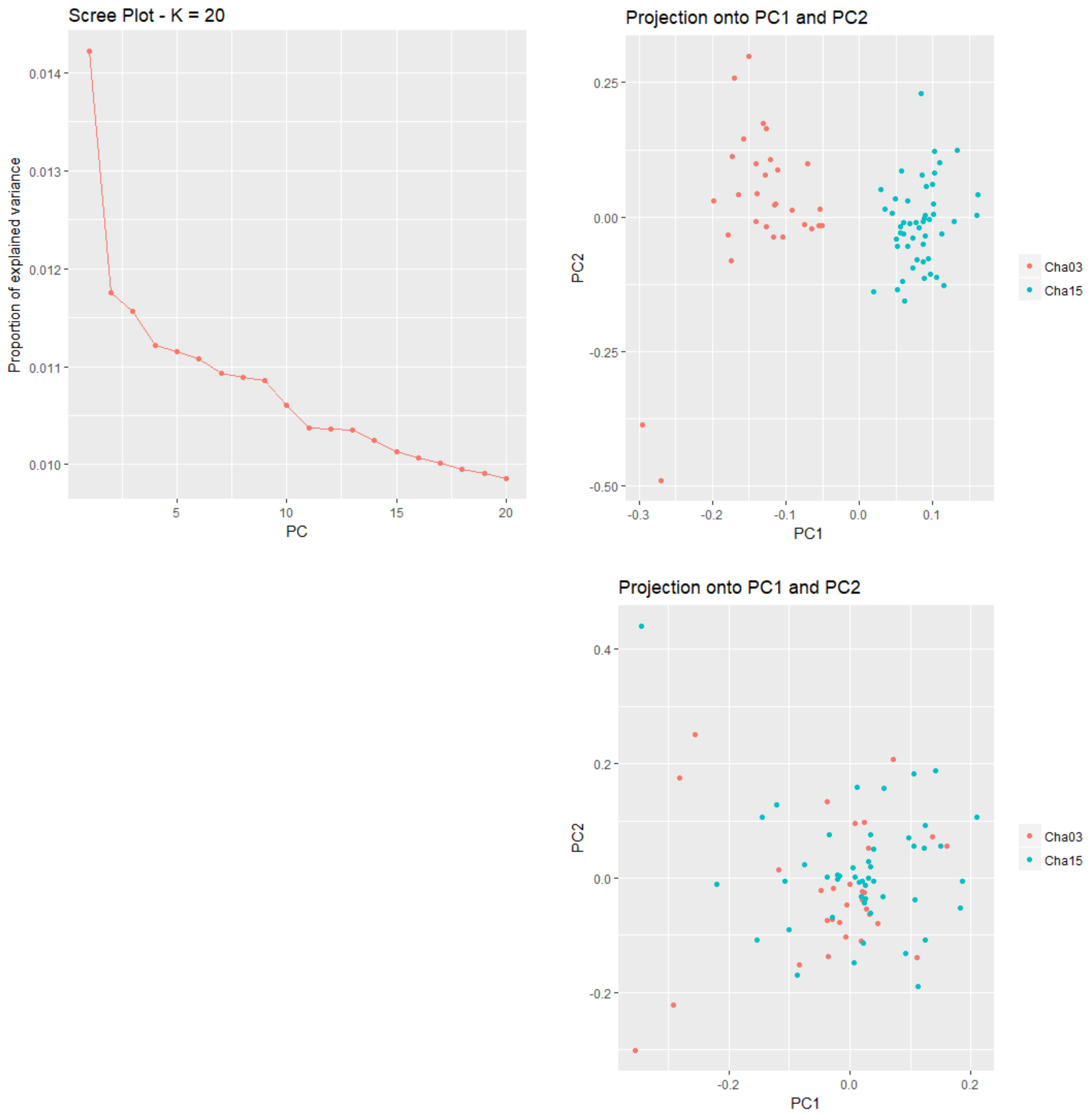

*Figure S8 Chalifour, m and f combined. Upper left is scree plot showing that k=2 is best. Top right is including outliers, bottom right is excluding outliers. Outliers play a role in generating population structure within this river between years, shown by changes in population structure when outliers are excluded.*

##list of outlier loci

[1] "866_78" "1137_78" "1721_78" "1910_79" "1972_79" "3995_79" "4975_78" "5410_79" "6284_10" "7083_79"
[11] "7205_77" "7393_79" "7583_78" "8039_8" "8206_79" "8525_79" "8979_6" "9415_78" "9985_79" "10151_77"
[21] "10154_65" "10696_78" "10903_77" "10957_77" "11235_78" "11521_76" "11606_77" "11815_79" "12116_77" "13160_47"

```
 [31] "13218_79"  "14431_79"  "15115_77"  "18075_79"  "18117_78"  "18719_79"  "19386_76"
"20090_61"  "20218_69"  "20708_79"
 [41] "21651_73"  "23670_78"  "23745_79"  "23907_79"  "28075_79"  "28547_78"  "29693_79"
"31717_34"  "32583_79"  "32700_77"
 [51] "35231_79"  "35262_76"  "36788_78"  "38592_75"  "38867_58"  "39743_77"  "39974_49"
"42735_20"  "43313_77"  "45325_76"
 [61] "45642_79"  "46439_76"  "46825_12"  "47058_78"  "47354_76"  "47695_78"  "48159_76"
"48329_79"  "49162_60"  "49921_75"
 [71] "49954_34"  "50404_26"  "51477_59"  "51976_28"  "53073_79"  "54508_79"  "55665_78"
"57110_77"  "57652_79"  "58632_29"
 [81] "62869_78"  "66231_29"  "66290_62"  "68634_75"  "70511_54"  "70882_78"  "72447_18"
"72620_77"  "72944_78"  "74712_73"
 [91] "75225_79"  "77093_45"  "77273_78"  "78044_40"  "78090_11"  "78754_11"  "80019_79"
"80680_79"  "81351_78"  "81730_77"
[101] "82672_77"  "82817_79"  "83973_79"  "85905_79"  "87141_78"  "87342_79"  "87508_12"
"87783_79"  "88453_78"  "88537_30"
[111] "88666_78"  "88895_78"  "89820_78"  "90763_79"  "91449_77"  "93522_79"  "94770_78"
"96420_52"  "96940_76"  "97531_77"
[121] "98454_43"  "98768_69"  "100129_79" "101049_10" "101352_79" "102737_78" "102740_43"
"108226_76" "115082_57"
```

=129/8513 SNPs = 1.5%

```
##note that changing the number of PCs does change the number of outliers found.
#20 SNPs are associated with PC2
> snp_pc <- get.pc(x, outliers)
> snp_pc
    SNP PC
1    18  1
2    21  1
3    36  1
4    48  1
5    53  1
6   208  1
7   277  1
8   305  1
9   384  2
10  454  1
11  467  1
12  491  1
13  512  1
14  555  1
15  566  1
16  590  1
17  624  2
18  665  1
19  723  1
20  741  1
21  742  1
22  794  1
23  814  1
```

| | | |
|---|---|---|
| 24 | 822 | 1 |
| 25 | 842 | 1 |
| 26 | 865 | 1 |
| 27 | 875 | 1 |
| 28 | 895 | 1 |
| 29 | 915 | 1 |
| 30 | 1022 | 1 |
| 31 | 1027 | 1 |
| 32 | 1149 | 1 |
| 33 | 1216 | 1 |
| 34 | 1457 | 1 |
| 35 | 1463 | 1 |
| 36 | 1522 | 1 |
| 37 | 1579 | 1 |
| 38 | 1649 | 2 |
| 39 | 1659 | 2 |
| 40 | 1693 | 1 |
| 41 | 1780 | 1 |
| 42 | 1966 | 1 |
| 43 | 1973 | 1 |
| 44 | 1987 | 1 |
| 45 | 2371 | 1 |
| 46 | 2405 | 1 |
| 47 | 2503 | 1 |
| 48 | 2686 | 2 |
| 49 | 2769 | 1 |
| 50 | 2783 | 1 |
| 51 | 3027 | 1 |
| 52 | 3031 | 1 |
| 53 | 3159 | 1 |
| 54 | 3322 | 1 |
| 55 | 3346 | 1 |
| 56 | 3416 | 1 |
| 57 | 3433 | 2 |
| 58 | 3654 | 1 |
| 59 | 3707 | 1 |
| 60 | 3860 | 1 |
| 61 | 3882 | 1 |
| 62 | 3958 | 1 |
| 63 | 3980 | 2 |
| 64 | 4005 | 1 |
| 65 | 4029 | 1 |
| 66 | 4053 | 1 |
| 67 | 4088 | 1 |
| 68 | 4101 | 1 |
| 69 | 4164 | 1 |
| 70 | 4222 | 1 |
| 71 | 4225 | 2 |
| 72 | 4256 | 2 |
| 73 | 4328 | 2 |
| 74 | 4360 | 2 |

| | | |
|---|---|---|
| 75 | 4421 | 1 |
| 76 | 4507 | 1 |
| 77 | 4551 | 1 |
| 78 | 4619 | 1 |
| 79 | 4651 | 1 |
| 80 | 4690 | 2 |
| 81 | 4870 | 1 |
| 82 | 5058 | 2 |
| 83 | 5064 | 1 |
| 84 | 5202 | 1 |
| 85 | 5319 | 1 |
| 86 | 5363 | 1 |
| 87 | 5505 | 1 |
| 88 | 5520 | 1 |
| 89 | 5557 | 1 |
| 90 | 5716 | 2 |
| 91 | 5759 | 1 |
| 92 | 5935 | 1 |
| 93 | 5954 | 1 |
| 94 | 6037 | 1 |
| 95 | 6042 | 1 |
| 96 | 6087 | 2 |
| 97 | 6202 | 1 |
| 98 | 6271 | 1 |
| 99 | 6345 | 1 |
| 100 | 6372 | 1 |
| 101 | 6474 | 1 |
| 102 | 6491 | 2 |
| 103 | 6592 | 1 |
| 104 | 6757 | 1 |
| 105 | 6843 | 1 |
| 106 | 6860 | 1 |
| 107 | 6872 | 2 |
| 108 | 6901 | 1 |
| 109 | 6971 | 1 |
| 110 | 6978 | 2 |
| 111 | 6993 | 1 |
| 112 | 7011 | 1 |
| 113 | 7087 | 1 |
| 114 | 7162 | 1 |
| 115 | 7225 | 1 |
| 116 | 7427 | 1 |
| 117 | 7559 | 1 |
| 118 | 7702 | 1 |
| 119 | 7748 | 1 |
| 120 | 7797 | 1 |
| 121 | 7870 | 1 |
| 122 | 7901 | 1 |
| 123 | 8025 | 1 |
| 124 | 8112 | 2 |
| 125 | 8132 | 1 |

126 8253  1
127 8254  1
128 8450  1
129 8486  2

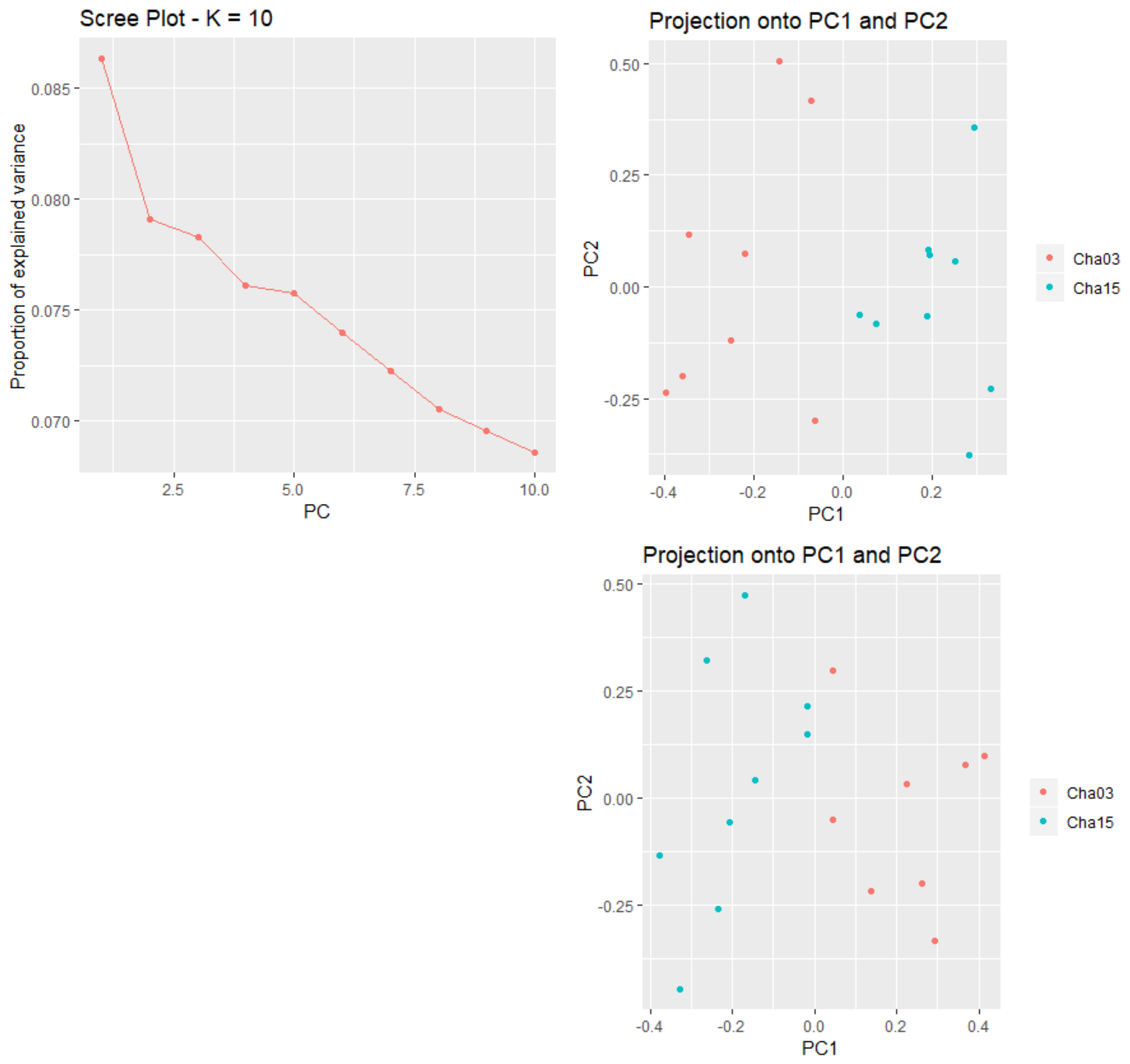

*Figure S9 Chalifour f, k = 2. Top, with outliers, bottom, without outliers. There is sufficient structure to determine outliers. Based on the number of individuals it isn't totally clear that the outliers are maintaining the structure present. Populations might overlap more when outliers are removed.*

> print(ID[outliers])
 [1] "3995_79"   "5684_73"   "7219_48"   "8525_79"   "12497_23"  "15859_28"  "20708_79"  "21808_48"  "23410_5"   "23745_79"
[11] "28075_79"  "31310_73"  "39743_77"  "46439_76"  "46592_79"  "51267_29"  "72339_30"  "76464_55"  "76666_67"  "79490_66"
[21] "81931_71"  "88280_65"  "90944_29"  "96126_78"  "103143_13"

SNP PC

| | | |
|---|---|---|
| 1 | 94 | 1 |
| 2 | 141 | 1 |
| 3 | 201 | 2 |
| 4 | 248 | 1 |
| 5 | 401 | 2 |
| 6 | 525 | 1 |
| 7 | 704 | 1 |
| 8 | 740 | 1 |
| 9 | 801 | 1 |
| 10 | 815 | 1 |
| 11 | 1004 | 1 |
| 12 | 1138 | 2 |
| 13 | 1481 | 1 |
| 14 | 1721 | 1 |
| 15 | 1725 | 1 |
| 16 | 1890 | 1 |
| 17 | 2340 | 2 |
| 18 | 2498 | 2 |
| 19 | 2511 | 2 |
| 20 | 2620 | 2 |
| 21 | 2730 | 2 |
| 22 | 2979 | 2 |
| 23 | 3073 | 1 |
| 24 | 3319 | 1 |
| 25 | 3598 | 1 |

PC1, 16/25

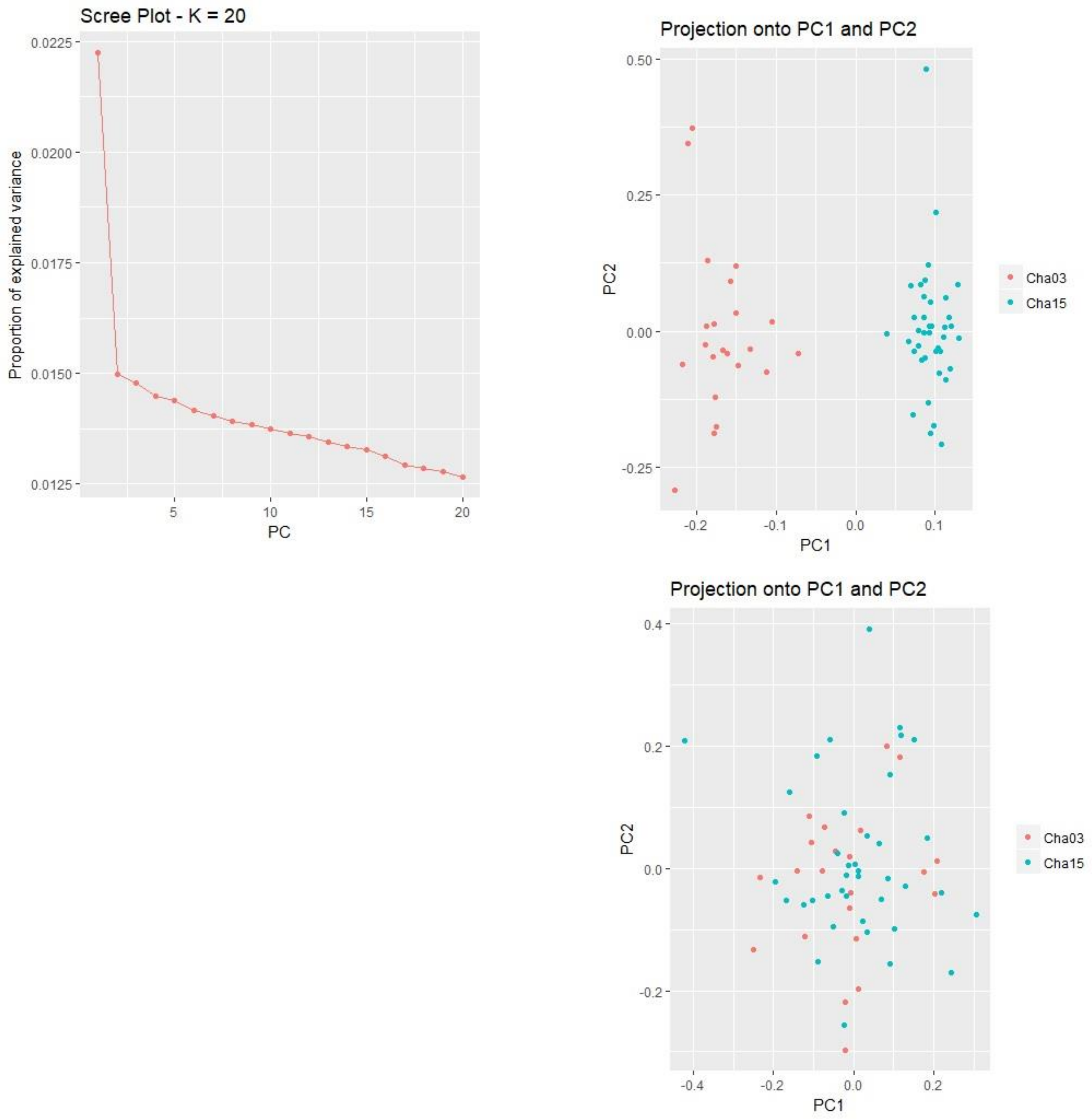

*Figure S10 Chalifour m, k = 2. Top left panel is the scree plot showing best k. Top right is including outliers and lower right is excluding outliers. Result: PC1 separates the timepoints.*

#outliers
[1] "866_78" "1137_78" "1721_78" "1745_79" "1910_79" "1972_76" "2776_77"
[8] "3049_79" "3084_15" "3995_78" "4066_36" "4431_19" "4885_78" "4975_78"
[15] "5155_43" "5302_79" "5410_79" "6056_10" "6527_51" "6583_79" "7083_79"
[22] "7205_77" "7393_79" "7664_78" "7828_36" "8039_8" "8206_78" "8521_79"
[29] "8525_79" "9283_28" "9290_77" "9415_78" "9629_79" "9942_79" "9985_79"
[36] "10154_65" "10157_79" "10696_78" "10903_77" "10916_78" "10957_77" "11235_78"
[43] "11433_78" "11521_76" "11761_37" "11815_79" "12116_77" "12894_79" "13192_59"
[50] "13218_79" "13263_13" "14431_76" "14895_79" "14996_79" "15115_77" "15863_78"
[57] "16462_79" "16510_18" "16851_79" "16860_79" "17897_78" "18075_78" "18117_78"
[64] "19036_78" "19262_77" "19386_76" "20154_78" "20357_79" "20708_79" "21844_79"
[71] "21874_78" "22119_76" "23670_78" "23745_79" "23771_79" "23907_79" "23960_64"

```
 [78] "24304_78"  "24957_79"  "25144_79"  "25199_77"  "25396_43"  "26211_77"  "26578_78"
 [85] "28026_42"  "28547_78"  "29069_79"  "29208_61"  "29284_77"  "29693_79"  "31135_79"
 [92] "31509_74"  "32228_78"  "32421_46"  "32583_79"  "32700_77"  "33919_79"  "34382_79"
 [99] "34542_79"  "34842_79"  "35231_79"  "35267_78"  "35878_58"  "36142_29"  "36714_75"
[106] "36788_78"  "36958_77"  "37237_77"  "37307_55"  "39371_78"  "39866_79"  "39983_22"
[113] "40111_28"  "40299_7"   "41112_79"  "41251_79"  "41774_78"  "41835_79"  "41986_78"
[120] "42673_76"  "42823_69"  "43077_77"  "43313_77"  "43351_78"  "44182_45"  "44786_79"
[127] "45325_76"  "45557_77"  "45594_57"  "45642_79"  "46825_12"  "46963_72"  "47058_78"
[134] "47354_76"  "47468_79"  "47695_78"  "48329_79"  "48513_31"  "48737_77"  "48833_75"
[141] "49162_60"  "49364_39"  "49548_30"  "49921_75"  "50987_25"  "52351_78"  "53073_79"
[148] "53362_77"  "55882_46"  "56093_48"  "57110_77"  "57652_79"  "58649_77"  "58827_77"
[155] "59518_78"  "60277_76"  "60974_79"  "62211_78"  "62222_79"  "62869_78"  "63988_78"
[162] "65789_33"  "66676_29"  "68634_75"  "68770_77"  "70693_79"  "70699_56"  "70882_78"
[169] "71655_77"  "72007_79"  "72079_78"  "72447_18"  "72620_77"  "72944_78"  "74712_73"
[176] "75225_79"  "75276_78"  "75295_75"  "76810_78"  "77032_78"  "77088_77"  "77093_45"
[183] "77332_78"  "77709_78"  "77786_79"  "78044_40"  "78552_47"  "79015_78"  "79047_76"
[190] "79613_79"  "80019_79"  "80664_78"  "80680_79"  "81320_50"  "81351_78"  "81405_79"
[197] "81596_78"  "81671_78"  "81730_77"  "81745_23"  "81922_79"  "82349_79"  "82672_77"
[204] "82679_76"  "82892_78"  "83863_67"  "83973_79"  "84062_75"  "85260_79"  "85504_79"
[211] "85977_63"  "86067_79"  "86409_78"  "86842_79"  "87113_79"  "87141_78"  "87276_75"
[218] "87342_79"  "87476_79"  "87758_78"  "87783_79"  "87972_79"  "88453_78"  "88749_71"
[225] "88815_79"  "88895_78"  "89780_78"  "89820_78"  "90422_77"  "90459_77"  "90763_78"
[232] "91449_77"  "91852_78"  "92230_78"  "92855_35"  "92867_79"  "93140_19"  "93437_61"
[239] "94159_79"  "94442_79"  "95224_79"  "95709_79"  "95720_79"  "95961_15"  "96076_79"
[246] "96533_78"  "96940_76"  "97365_77"  "97657_76"  "98454_43"  "98768_69"  "99817_38"
[253] "100129_79" "101352_79" "101890_19" "102377_79" "102505_79" "102737_78" "102857_77"
[260] "105475_78" "108226_76" "117457_61" "118233_49"
```

#that is a lot of outliers
=263/9294 SNPs = 2.8% of SNPs are outliers.

PC2, 30 loci

```
> snp_pc
   SNP PC
1    17  1
2    20  1
3    36  1
4    37  1
5    50  1
6    55  1
7   119  1
8   154  1
9   159  2
10  228  1
11  237  2
12  270  1
13  305  1
14  310  1
15  324  2
16  332  1
```

| | | |
|---|---|---|
| 17 | 341 | 1 |
| 18 | 410 | 1 |
| 19 | 457 | 1 |
| 20 | 462 | 1 |
| 21 | 502 | 1 |
| 22 | 517 | 1 |
| 23 | 541 | 1 |
| 24 | 571 | 1 |
| 25 | 586 | 2 |
| 26 | 607 | 1 |
| 27 | 618 | 1 |
| 28 | 643 | 1 |
| 29 | 644 | 1 |
| 30 | 705 | 1 |
| 31 | 706 | 1 |
| 32 | 721 | 1 |
| 33 | 742 | 1 |
| 34 | 781 | 1 |
| 35 | 784 | 1 |
| 36 | 803 | 1 |
| 37 | 804 | 1 |
| 38 | 857 | 1 |
| 39 | 877 | 1 |
| 40 | 878 | 1 |
| 41 | 886 | 1 |
| 42 | 909 | 1 |
| 43 | 934 | 1 |
| 44 | 939 | 1 |
| 45 | 962 | 2 |
| 46 | 973 | 1 |
| 47 | 993 | 1 |
| 48 | 1083 | 1 |
| 49 | 1117 | 2 |
| 50 | 1119 | 1 |
| 51 | 1127 | 2 |
| 52 | 1250 | 1 |
| 53 | 1307 | 1 |
| 54 | 1318 | 1 |
| 55 | 1327 | 1 |
| 56 | 1397 | 1 |
| 57 | 1458 | 1 |
| 58 | 1464 | 1 |
| 59 | 1499 | 1 |
| 60 | 1500 | 1 |
| 61 | 1583 | 1 |
| 62 | 1603 | 1 |
| 63 | 1611 | 1 |
| 64 | 1707 | 1 |
| 65 | 1726 | 1 |
| 66 | 1739 | 1 |
| 67 | 1825 | 1 |

| | | |
|---|---|---|
| 68 | 1840 | 1 |
| 69 | 1869 | 1 |
| 70 | 1975 | 1 |
| 71 | 1980 | 1 |
| 72 | 2010 | 1 |
| 73 | 2162 | 1 |
| 74 | 2173 | 1 |
| 75 | 2175 | 1 |
| 76 | 2190 | 1 |
| 77 | 2196 | 1 |
| 78 | 2231 | 1 |
| 79 | 2297 | 1 |
| 80 | 2316 | 1 |
| 81 | 2321 | 1 |
| 82 | 2340 | 1 |
| 83 | 2427 | 1 |
| 84 | 2478 | 1 |
| 85 | 2598 | 1 |
| 86 | 2640 | 1 |
| 87 | 2694 | 1 |
| 88 | 2706 | 2 |
| 89 | 2714 | 1 |
| 90 | 2749 | 1 |
| 91 | 2894 | 1 |
| 92 | 2925 | 1 |
| 93 | 2997 | 1 |
| 94 | 3018 | 2 |
| 95 | 3034 | 1 |
| 96 | 3048 | 1 |
| 97 | 3170 | 1 |
| 98 | 3215 | 1 |
| 99 | 3234 | 1 |
| 100 | 3268 | 1 |
| 101 | 3312 | 1 |
| 102 | 3317 | 1 |
| 103 | 3376 | 1 |
| 104 | 3406 | 2 |
| 105 | 3450 | 1 |
| 106 | 3458 | 1 |
| 107 | 3475 | 1 |
| 108 | 3506 | 1 |
| 109 | 3516 | 2 |
| 110 | 3708 | 1 |
| 111 | 3753 | 1 |
| 112 | 3762 | 1 |
| 113 | 3773 | 2 |
| 114 | 3789 | 1 |
| 115 | 3855 | 1 |
| 116 | 3869 | 1 |
| 117 | 3910 | 1 |
| 118 | 3921 | 1 |

| | | |
|---|---|---|
| 119 | 3930 | 1 |
| 120 | 3995 | 2 |
| 121 | 4006 | 2 |
| 122 | 4030 | 1 |
| 123 | 4055 | 1 |
| 124 | 4061 | 1 |
| 125 | 4129 | 1 |
| 126 | 4179 | 1 |
| 127 | 4227 | 1 |
| 128 | 4247 | 1 |
| 129 | 4249 | 1 |
| 130 | 4252 | 1 |
| 131 | 4353 | 1 |
| 132 | 4370 | 2 |
| 133 | 4379 | 1 |
| 134 | 4405 | 1 |
| 135 | 4413 | 1 |
| 136 | 4428 | 1 |
| 137 | 4477 | 1 |
| 138 | 4493 | 2 |
| 139 | 4511 | 1 |
| 140 | 4521 | 1 |
| 141 | 4544 | 1 |
| 142 | 4554 | 2 |
| 143 | 4571 | 2 |
| 144 | 4602 | 1 |
| 145 | 4679 | 2 |
| 146 | 4779 | 1 |
| 147 | 4826 | 1 |
| 148 | 4842 | 1 |
| 149 | 4989 | 1 |
| 150 | 5000 | 2 |
| 151 | 5051 | 1 |
| 152 | 5083 | 1 |
| 153 | 5129 | 1 |
| 154 | 5139 | 1 |
| 155 | 5156 | 1 |
| 156 | 5202 | 1 |
| 157 | 5234 | 1 |
| 158 | 5292 | 1 |
| 159 | 5294 | 1 |
| 160 | 5329 | 1 |
| 161 | 5394 | 1 |
| 162 | 5511 | 2 |
| 163 | 5571 | 1 |
| 164 | 5696 | 1 |
| 165 | 5704 | 1 |
| 166 | 5841 | 1 |
| 167 | 5842 | 1 |
| 168 | 5865 | 1 |
| 169 | 5940 | 1 |

| | | |
|---|---|---|
| 170 | 5977 | 1 |
| 171 | 5987 | 1 |
| 172 | 6019 | 1 |
| 173 | 6036 | 1 |
| 174 | 6072 | 1 |
| 175 | 6240 | 2 |
| 176 | 6290 | 1 |
| 177 | 6299 | 1 |
| 178 | 6302 | 1 |
| 179 | 6453 | 1 |
| 180 | 6478 | 1 |
| 181 | 6485 | 1 |
| 182 | 6486 | 1 |
| 183 | 6512 | 1 |
| 184 | 6553 | 1 |
| 185 | 6560 | 1 |
| 186 | 6593 | 1 |
| 187 | 6631 | 2 |
| 188 | 6679 | 1 |
| 189 | 6681 | 1 |
| 190 | 6741 | 1 |
| 191 | 6777 | 1 |
| 192 | 6849 | 1 |
| 193 | 6851 | 1 |
| 194 | 6927 | 2 |
| 195 | 6930 | 1 |
| 196 | 6932 | 1 |
| 197 | 6950 | 1 |
| 198 | 6955 | 1 |
| 199 | 6959 | 1 |
| 200 | 6961 | 2 |
| 201 | 6987 | 1 |
| 202 | 7033 | 1 |
| 203 | 7070 | 1 |
| 204 | 7072 | 1 |
| 205 | 7098 | 1 |
| 206 | 7185 | 1 |
| 207 | 7198 | 1 |
| 208 | 7202 | 1 |
| 209 | 7302 | 1 |
| 210 | 7327 | 1 |
| 211 | 7378 | 1 |
| 212 | 7382 | 1 |
| 213 | 7412 | 1 |
| 214 | 7442 | 1 |
| 215 | 7464 | 1 |
| 216 | 7467 | 1 |
| 217 | 7477 | 1 |
| 218 | 7485 | 1 |
| 219 | 7497 | 1 |
| 220 | 7526 | 1 |

| | | |
|---|---|---|
| 221 | 7528 | 1 |
| 222 | 7546 | 1 |
| 223 | 7601 | 1 |
| 224 | 7628 | 2 |
| 225 | 7636 | 1 |
| 226 | 7644 | 1 |
| 227 | 7725 | 1 |
| 228 | 7729 | 1 |
| 229 | 7792 | 1 |
| 230 | 7796 | 1 |
| 231 | 7818 | 1 |
| 232 | 7885 | 1 |
| 233 | 7922 | 1 |
| 234 | 7966 | 1 |
| 235 | 8025 | 1 |
| 236 | 8028 | 1 |
| 237 | 8061 | 2 |
| 238 | 8091 | 1 |
| 239 | 8168 | 1 |
| 240 | 8194 | 1 |
| 241 | 8271 | 1 |
| 242 | 8324 | 1 |
| 243 | 8326 | 1 |
| 244 | 8340 | 2 |
| 245 | 8352 | 2 |
| 246 | 8396 | 1 |
| 247 | 8431 | 1 |
| 248 | 8478 | 1 |
| 249 | 8499 | 1 |
| 250 | 8568 | 1 |
| 251 | 8600 | 1 |
| 252 | 8697 | 1 |
| 253 | 8721 | 1 |
| 254 | 8840 | 1 |
| 255 | 8879 | 2 |
| 256 | 8929 | 1 |
| 257 | 8943 | 1 |
| 258 | 8965 | 1 |
| 259 | 8978 | 1 |
| 260 | 9169 | 1 |
| 261 | 9178 | 1 |
| 262 | 9260 | 1 |
| 263 | 9267 | 1 |

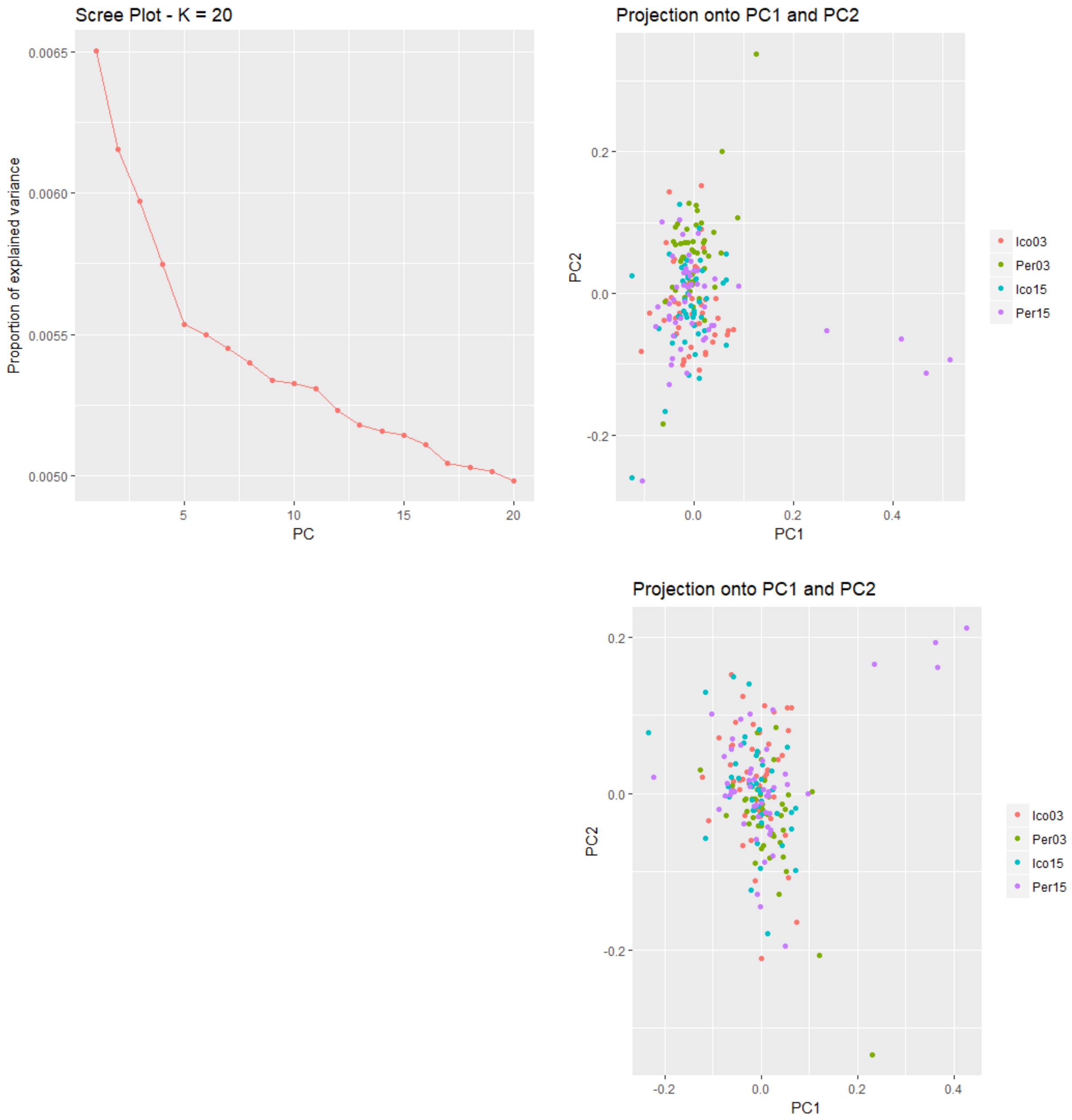

*Figure S11 Ic-Pe, m and f combined. Top left is the scree plot showing that k = 5 looks best. However, contribution of PC3 to population structure was tested, and it does not change the conclusion. Thus, k=2 was chosen. Top right is with outlier loci included, and bottom right is with outlier loci excluded. Conclusion: although structure is relatively weak within Ic-Pe between years, there is structure, and outlier loci contribute to this structure. Pe03 separates out on PC2. In SNP list associated with PCs below, you can see that 8 SNPs are associated with this PC.*

```
> print(ID[outliers])
 [1] "3718_64"  "7174_45"  "10957_77" "16314_49" "18923_13" "19294_69" "20378_78" "23360_77"
 [9] "31135_79" "31873_28" "32359_23" "33513_45" "35165_18" "41903_57" "43027_6"  "44920_15"
[17] "46714_51" "50605_54" "54214_10" "66470_39" "72908_11" "73469_62" "73918_69" "77907_75"
[25] "79366_26" "80339_30" "86537_49" "86863_71" "87113_79" "93777_10" "96469_21" "96847_31"
```

```
[33] "99581_77"
```

=33/6802 = 0.49% of loci are outliers. Note that there are 51 outliers when k=5 is used. When k=5 is used, there are some SNPs that assign to Pcs 3, 4 and 5.

```
    SNP PC
1   141  2
2   352  1
3   613  2
4  1000  1
5  1179  1
6  1206  1
7  1294  2
8  1502  2
9  2089  2
10 2142  1
11 2177  1
12 2277  1
13 2405  1
14 2851  1
15 2929  2
16 3048  1
17 3158  1
18 3403  1
19 3561  1
20 4016  1
21 4400  1
22 4441  1
23 4471  1
24 4771  1
25 4877  1
26 4959  1
27 5409  1
28 5424  1
29 5437  2
30 5970  1
31 6180  1
32 6203  1
33 6399  2
```

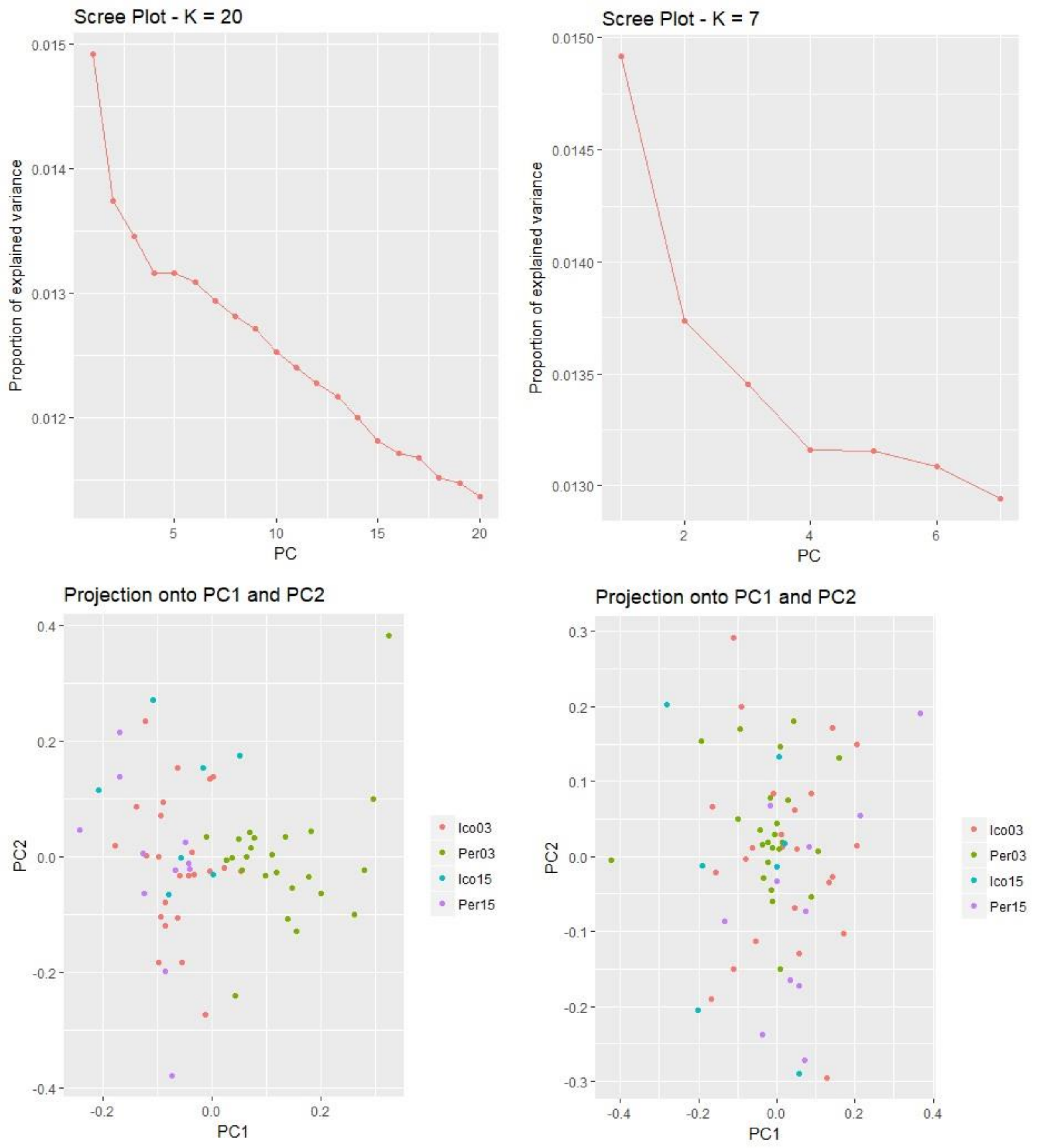

*Figure S12 Ic-Pe f, k=2. Top panels are scree plots used to determine best k. Bottom plots are (right) with outlier loci included, and (left) excluding outlier loci. Result: PC1 separates Pe03 from the other populations.*

```
> print(ID[outliers])
 [1] "639_75"    "3193_79"   "4975_78"   "7583_78"   "8169_79"   "8206_79"   "10957_77"
 [8] "13323_72"  "14431_76"  "16324_79"  "18171_79"  "18297_78"  "19848_54"  "19893_60"
[15] "23360_77"  "23519_75"  "24836_77"  "24957_79"  "25199_77"  "27499_78"  "28075_78"
```

```
[22] "28601_43"  "28887_76"  "29693_78"  "31509_74"  "32239_7"   "32700_77"  "32851_77"
[29] "34803_77"  "34842_79"  "35262_76"  "35267_78"  "41258_78"  "41419_78"  "42767_66"
[36] "45557_77"  "46292_78"  "46573_78"  "47695_78"  "48757_78"  "49415_76"  "50171_79"
[43] "55665_78"  "61999_58"  "62150_77"  "64088_78"  "70726_79"  "71278_78"  "72944_79"
[50] "73353_79"  "73469_62"  "75276_78"  "75331_77"  "76823_79"  "77966_78"  "83973_79"
[57] "84062_75"  "87476_79"  "88895_78"  "89347_55"  "89405_57"  "94671_78"  "94835_32"
[64] "96076_79"  "96289_77"  "97365_77"  "98249_5"   "99081_19"  "99581_77"  "99999_73"
[71] "102675_77" "104310_77" "104326_23"
```

##whoa, that is a lot of outliers. 73/6442 SNPs = 1.1% of SNPs

##note that this is the same result as we saw with both males and females included in the dataset

```
> snp_pc <- get.pc(x, outliers)
> snp_pc
    SNP PC
1    11  1
2   120  1
3   214  1
4   363  1
5   396  1
6   398  1
7   560  1
8   802  1
9   918  1
10 1044  1
11 1396  1
12 1411  1
13 1435  2
14 1510  1
15 1520  1
16 1537  1
17 1660  2
18 1708  1
19 1741  1
20 1775  1
21 1800  1
22 1854  1
23 1997  1
24 2050  2
25 2086  1
26 2104  1
27 2247  1
28 2250  1
29 2282  1
30 2283  1
31 2639  2
32 2660  1
33 2669  1
34 2670  1
35 2914  1
```

| | | |
|---|---|---|
| 36 | 2963 | 1 |
| 37 | 2972 | 1 |
| 38 | 3046 | 1 |
| 39 | 3106 | 1 |
| 40 | 3148 | 1 |
| 41 | 3193 | 1 |
| 42 | 3405 | 1 |
| 43 | 3612 | 1 |
| 44 | 3685 | 1 |
| 45 | 3789 | 2 |
| 46 | 3982 | 1 |
| 47 | 4032 | 1 |
| 48 | 4150 | 1 |
| 49 | 4180 | 1 |
| 50 | 4270 | 1 |
| 51 | 4311 | 1 |
| 52 | 4317 | 1 |
| 53 | 4383 | 2 |
| 54 | 4412 | 1 |
| 55 | 4512 | 1 |
| 56 | 4950 | 1 |
| 57 | 4953 | 1 |
| 58 | 5170 | 1 |
| 59 | 5280 | 1 |
| 60 | 5337 | 1 |
| 61 | 5683 | 1 |
| 62 | 5717 | 1 |
| 63 | 5768 | 2 |
| 64 | 5824 | 1 |
| 65 | 5837 | 1 |
| 66 | 5918 | 1 |
| 67 | 6025 | 2 |
| 68 | 6031 | 1 |
| 69 | 6057 | 1 |
| 70 | 6091 | 2 |
| 71 | 6256 | 1 |
| 72 | 6392 | 1 |
| 73 | 6394 | 2 |

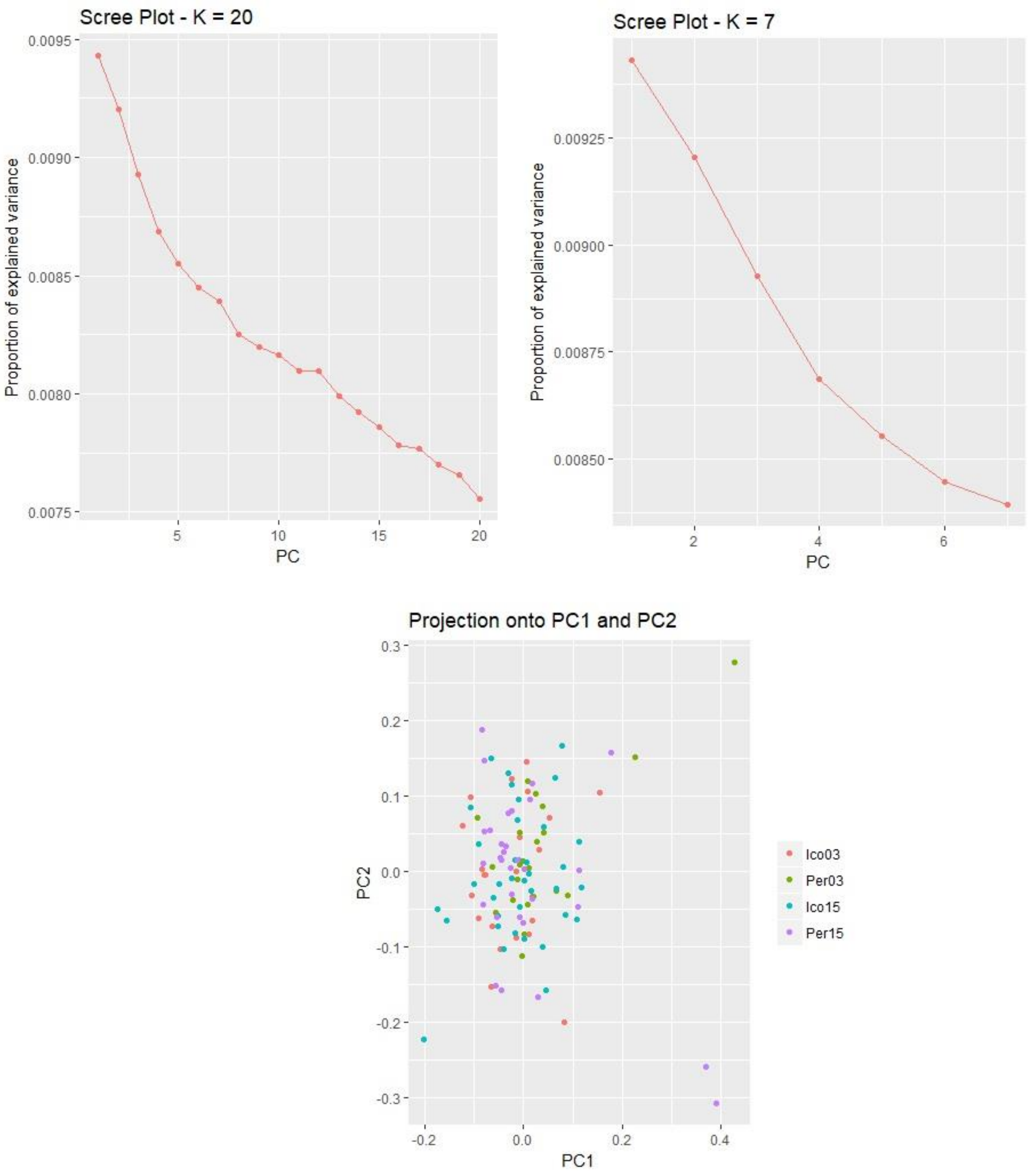

*Figure S13 Ic-Pe m, k = 4. Top panels are scree plots showing that there is insufficient structure to find a best k. Bottom panel is including outliers. Both k = 4 and k = 6 were tested with the same result.*

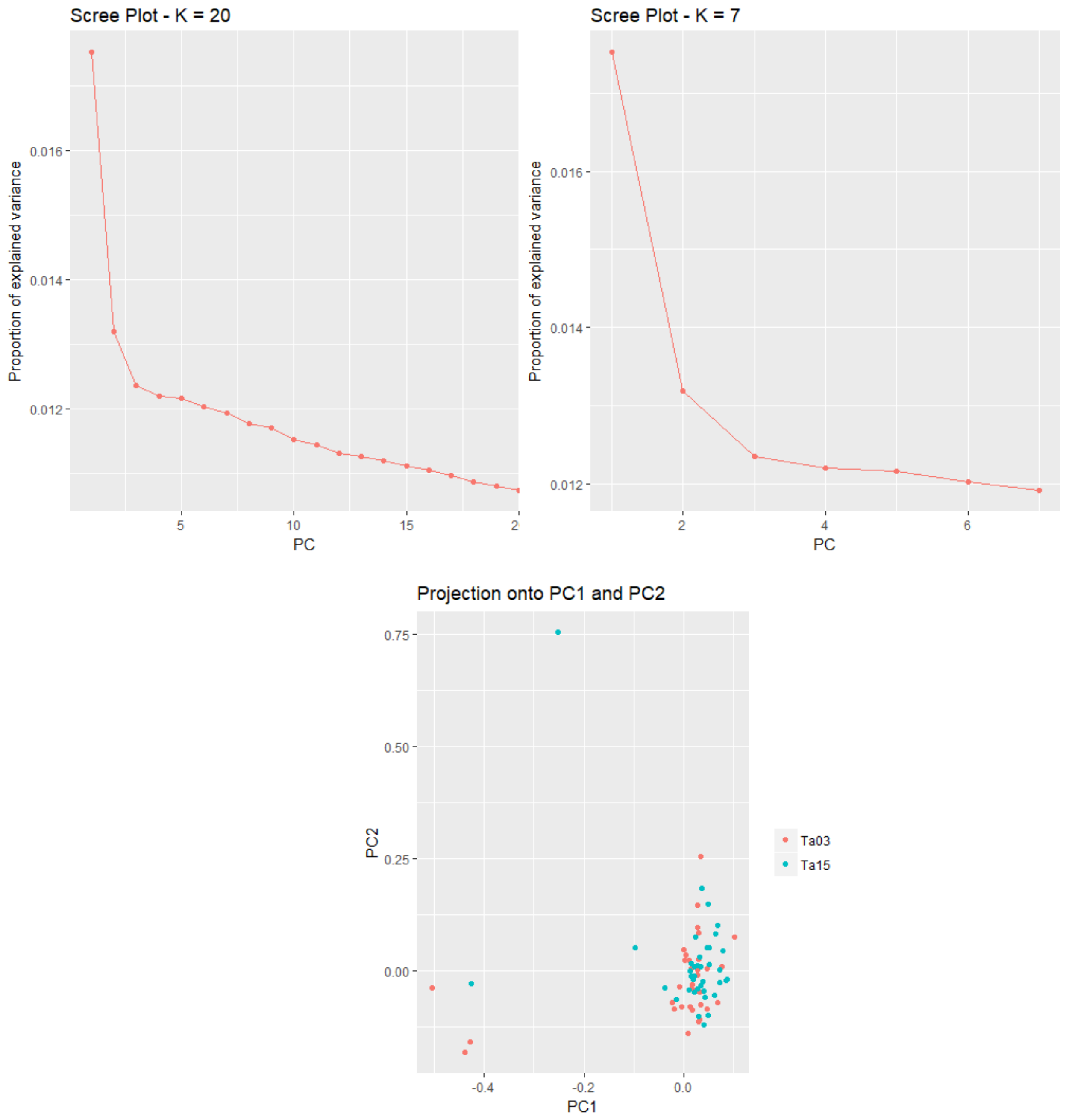

*Figure S14 Takwa, m and f combined k=2, The top panel is scree plots showing the best k. Bottom panel is with outliers included, and shows that there is no structure between years. The structure seen in the scree plot is likely due to those outlier individuals.*

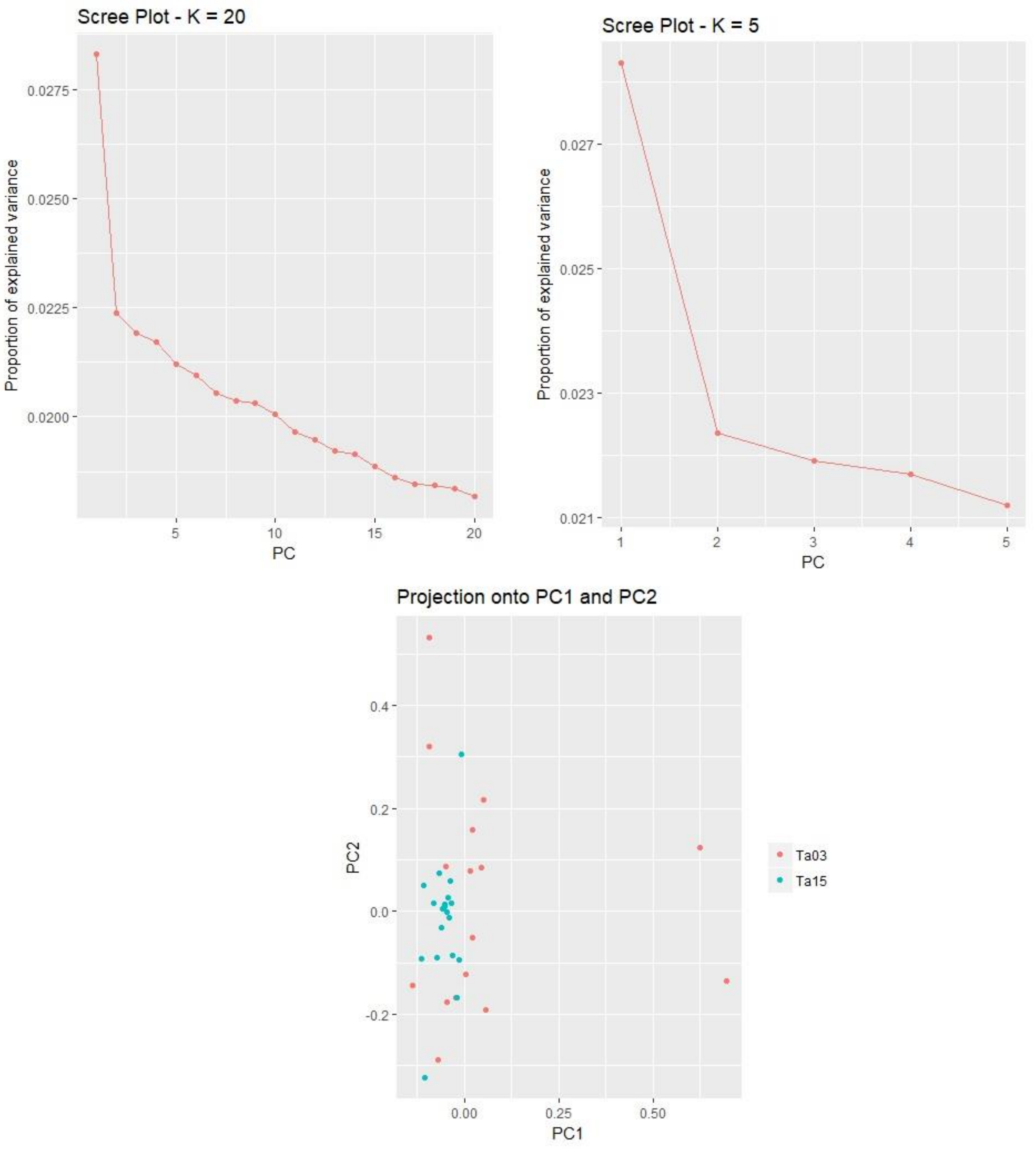

*Figure S15 Takwa River, f, k=2. Top panels are scree plots used to find the best k. Bottom panel is including outliers. Result: There is no structure between years within Takwa River. Outliers cannot be assessed.*

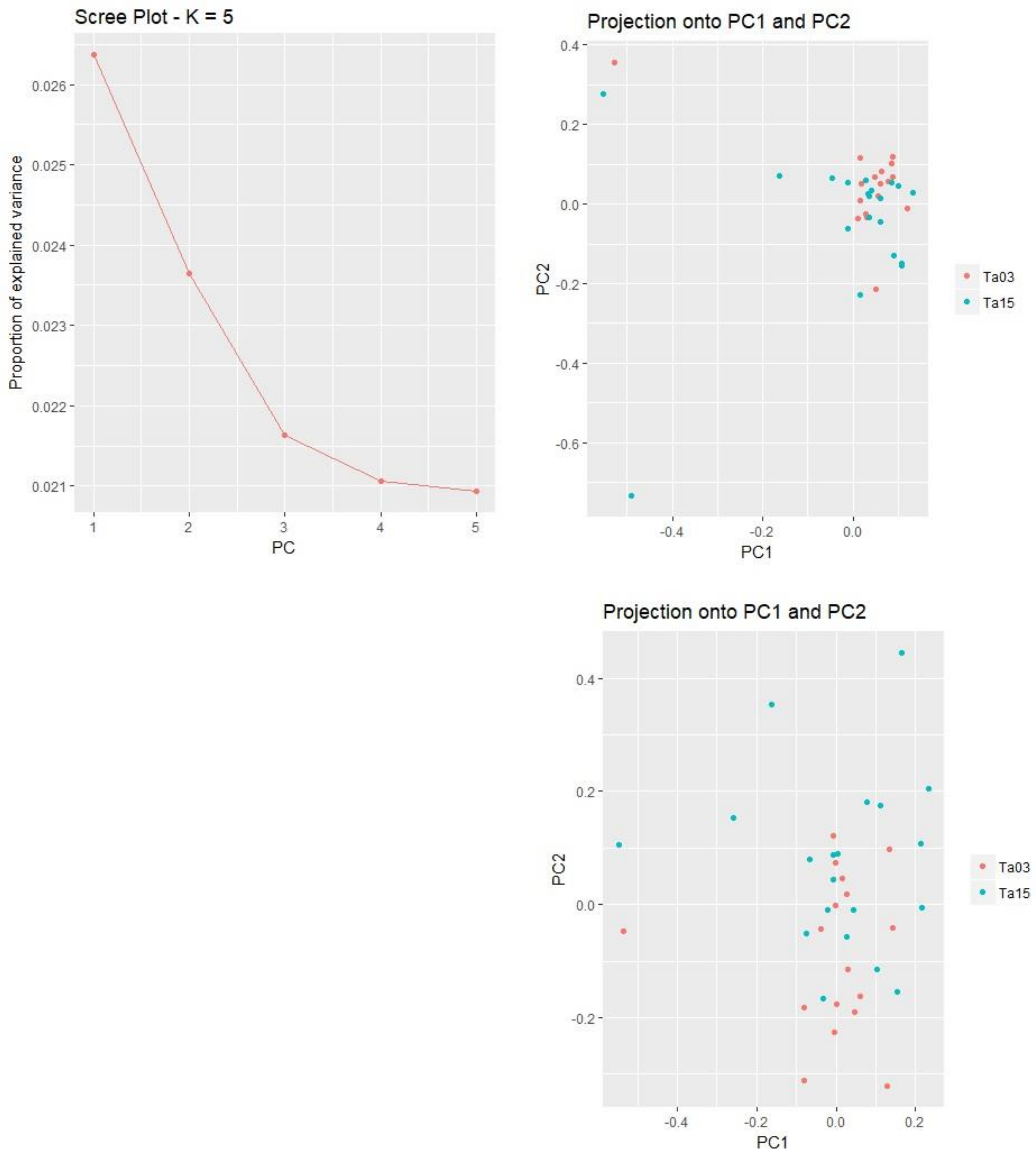

*Figure S16 Takwa m, k=3. Top left panel showing scree plot used to find best k. Top right panel is including outliers. Bottom right panel is excluding outliers. Result: there is no population structure between years, and thus no outliers can be found.*

Outlier loci in common

Comparing A (all_all) and B, (ALL_F): The sets have 8 SNPs in common:[1] "2764_47"   "16571_28" "34289_47"  "36864_48"  "45711_42"  "89842_9"   "93560_78"  "103420_43"

Comparing A (all_all) and C, (all, M): The sets have 59 SNPs in common: [1] "2764_47"   "5777_26" "7221_52"   "9664_69"   "10060_79"  "10881_53"  "11507_79"  "12647_55"
[9] "15362_57"  "15426_49"  "16571_28"  "23892_66"  "25728_47"  "26281_46"  "27021_71" "28192_35"
[17] "29788_76"  "31886_71"  "35235_60"  "36864_48"  "37605_43"  "38039_35"  "39685_62" "41527_40"
[25] "41635_58"  "45711_42"  "46331_65"  "48658_18"  "49504_49"  "50030_36"  "50404_26" "51264_20"
[33] "53498_56"  "62465_26"  "65438_60"  "65770_56"  "69567_54"  "71554_26"  "76517_75" "81466_14"
[41] "83593_61"  "84174_76"  "85630_70"  "88020_72"  "88250_71"  "89842_9"   "91022_14" "91315_70"
[49] "91507_23"  "93560_78"  "93677_42"  "97401_34"  "97840_17"  "99888_60"  "101083_6" "102928_19"
[57] "103420_43" "104231_8"  "104441_34"

Comparing A (all_all) and D (sou, all): The sets have 9 SNPs in common:[1] "16571_28"  "27021_71" "34289_47"  "37605_43"  "48658_18"  "53498_56"  "74122_76"  "103420_43"
[9] "104441_34"

Comparing A (all_all) and E, (SOU_F): The sets have 2 SNPs in common:[1] "16571_28" "34289_47"

Comparing A (all_all) and F: The sets have 13 SNPs in common: [1] "7221_52"   "9664_69" "16571_28"  "27021_71"  "34289_47"  "37605_43"  "38039_35"  "48658_18"
[9] "50030_36"  "53498_56"  "74122_76"  "103420_43" "104441_34"

Comparing A (all_all) and G, (IC-PE, ALL): The sets have 0 SNPs in common:character(0)

Comparing A (all_all) and H, (IC-PE, F): The sets have 0 SNPs in common:character(0)

Comparing A (all_all) and I, (CH_ALL): The sets have 1 SNPs in common:[1] "50404_26"

Comparing A (all_all) and J, (CH_F): The sets have 0 SNPs in common:character(0)

Comparing A (all_all) and K, (CH_M): The sets have 0 SNPs in common:character(0)

Comparing B, (ALL_F) and C , (all, M): The sets have 7 SNPs in common:[1] "2764_47"   "16571_28" "36864_48"  "45711_42"  "89842_9"   "93560_78"  "103420_43"

Comparing B, (ALL_F) and D (sou, all): The sets have 3 SNPs in common:[1] "16571_28"  "34289_47" "103420_43"

Comparing B, (ALL_F) and E (SOU_F), (SOU_F): The sets have 3 SNPs in common:[1] "16571_28" "27138_61" "34289_47"

Comparing B, (ALL_F) and F: The sets have 3 SNPs in common:[1] "16571_28"  "34289_47" "103420_43"

Comparing B, (ALL_F) and G, (IC-PE, ALL): The sets have 0 SNPs in common:character(0)

Comparing B, (ALL_F) and H, (IC-PE, F): The sets have 0 SNPs in common:character(0)

Comparing B, (ALL_F) and I, (CH_ALL): The sets have 0 SNPs in common:character(0)

Comparing B, (ALL_F) and J, (CH_F): The sets have 0 SNPs in common:character(0)

Comparing B, (ALL_F) and K, (CH_M): The sets have 0 SNPs in common:character(0)

Comparing C, (all, M) and D: The sets have 15 SNPs in common: [1] "9415_78" "9942_79" "16571_28" "27021_71" "31135_79" "34803_77" "34842_79" "37605_43"
[9] "48658_18" "53498_56" "62534_37" "72944_78" "87113_79" "103420_43" "104441_34"
Comparing C, (all, M) and E, (SOU_F): The sets have 3 SNPs in common:[1] "16571_28" "34803_77" "34842_79"

Comparing C, (all, M) and F (SOU_M): The sets have 20 SNPs in common: [1] "7221_52" "9415_78" "9664_69" "9942_79" "16571_28" "21844_79" "26667_79" "27021_71"
[9] "31135_79" "37605_43" "38039_35" "48658_18" "50030_36" "53498_56" "62534_37" "72944_78"
[17] "87113_79" "87783_79" "103420_43" "104441_34"

Comparing C, (all, M) and G, (IC-PE, ALL): The sets have 2 SNPs in common:[1] "31135_79" "87113_79"

Comparing C, (all, M) and H, (IC-PE, F): The sets have 4 SNPs in common:[1] "14431_76" "34803_77" "34842_79" "45557_77"

Comparing C, (all, M) and I, (CH_ALL): The sets have 10 SNPs in common: [1] "1721_78" "7393_79" "9415_78" "10696_78" "50404_26" "72447_18" "72944_78" "87783_79"
[9] "96940_76" "101049_10"

Comparing C, (all, M) and J, (CH_F): The sets have 0 SNPs in common:character(0)

Comparing C, (all, M) and K, (CH_M): The sets have 37 SNPs in common: [1] "1721_78" "7393_79" "9415_78" "9629_79" "9942_79" "10157_79" "10696_78" "14431_76" "19262_77"
[10] "21844_79" "23960_64" "31135_79" "32228_78" "34382_79" "34842_79" "41986_78" "43077_77" "43351_78"
[19] "44786_79" "45557_77" "59518_78" "63988_78" "71655_77" "72447_18" "72944_78" "79047_76" "80664_78"
[28] "82349_79" "82679_76" "86067_79" "87113_79" "87783_79" "87972_79" "90459_77" "91852_78" "94159_79"
[37] "96940_76"

Comparing D, (SOU_ALL) and E, (SOU_F): The sets have 23 SNPs in common: [1] "3193_79" "4975_78" "7583_78" "10957_77" "16571_28" "23360_77" "24957_79" "28075_78"
[9] "29693_78" "34289_47" "34803_77" "34842_79" "35262_76" "47695_78" "70726_79" "74568_79"
[17] "75276_78" "83973_79" "88895_78" "92110_78" "96076_79" "99202_76" "104310_77"

Comparing D, (SOU_ALL) and F (SOU_M): The sets have 27 SNPs in common: [1] "866_78" "5302_79" "7583_78" "9415_78" "9942_79" "10957_77" "11235_78" "16571_28"
[9] "24957_79" "27021_71" "31135_79" "34289_47" "37605_43" "47695_78" "48658_18" "53498_56"
[17] "56508_40" "62534_37" "70726_79" "72944_78" "74122_76" "75276_78" "87113_79" "89820_78"
[25] "103420_43" "104310_77" "104441_34"

Comparing D, (SOU_ALL) and G, (IC-PE, ALL): The sets have 4 SNPs in common:[1] "10957_77" "23360_77" "31135_79" "87113_79"

Comparing D, (SOU_ALL) and H, (IC-PE, F): The sets have 18 SNPs in common: [1] "3193_79" "4975_78" "7583_78" "10957_77" "23360_77" "24957_79" "28075_78" "29693_78"
[9] "34803_77" "34842_79" "35262_76" "47695_78" "70726_79" "75276_78" "83973_79" "88895_78"
[17] "96076_79" "104310_77"

Comparing D, (SOU_ALL) and I, (CH_ALL): The sets have 12 SNPs in common: [1] "866_78" "4975_78" "7583_78" "9415_78" "10957_77" "11235_78" "35262_76" "47695_78" "72944_78"
[10] "83973_79" "88895_78" "89820_78"

Comparing D, (SOU_ALL) and J, (CH_F): The sets have 0 SNPs in common:character(0)

Comparing D, (SOU_ALL) and K, (CH_M): The sets have 20 SNPs in common: [1] "866_78" "4975_78" "5302_79" "8206_78" "9415_78" "9942_79" "10957_77" "11235_78" "24957_79"
[10] "29208_61" "31135_79" "34842_79" "47695_78" "72944_78" "75276_78" "83973_79" "87113_79" "88895_78"
[19] "89820_78" "96076_79"

Comparing E, (SOU_F) and F (SOU_M): The sets have 9 SNPs in common:[1] "7583_78" "10957_77" "16571_28" "24957_79" "34289_47" "47695_78" "70726_79" "75276_78"
[9] "104310_77"

Comparing E, (SOU_F) and G, (IC-PE, ALL): The sets have 2 SNPs in common:[1] "10957_77" "23360_77"

Comparing E, (SOU_F) and H, (IC-PE, F): The sets have 31 SNPs in common: [1] "3193_79" "4975_78" "7583_78" "8206_79" "10957_77" "18297_78" "23360_77" "23519_75"
[9] "24957_79" "28075_78" "29693_78" "31509_74" "32700_77" "34803_77" "34842_79" "35262_76"
[17] "41258_78" "46573_78" "47695_78" "48757_78" "50171_79" "55665_78" "62150_77" "70726_79"
[25] "71278_78" "75276_78" "83973_79" "88895_78" "96076_79" "102675_77" "104310_77"

Comparing E, (SOU_F) and I, (CH_ALL): The sets have 14 SNPs in common: [1] "4975_78" "7583_78" "8206_79" "10957_77" "11815_79" "32700_77" "35262_76" "46439_76" "47695_78"
[10] "55665_78" "77273_78" "80680_79" "83973_79" "88895_78"

Comparing E, (SOU_F) and J, (CH_F): The sets have 2 SNPs in common:[1] "46439_76" "46592_79"

Comparing E, (SOU_F) and K, (CH_M): The sets have 13 SNPs in common: [1] "4975_78"  "10957_77" "11815_79" "24957_79" "31509_74" "32700_77" "34842_79" "47695_78" "75276_78"
[10] "80680_79" "83973_79" "88895_78" "96076_79"

Comparing F, (SOU_M) and G, (IC-PE, ALL): The sets have 4 SNPs in common:[1] "10957_77" "20378_78" "31135_79" "87113_79"

Comparing F, (SOU_M) and H, (IC-PE, F): The sets have 8 SNPs in common:[1] "7583_78" "10957_77"  "24957_79"  "47695_78"  "70726_79"  "73353_79"  "75276_78"  "104310_77"

Comparing F, (SOU_M) and I, (CH_ALL), (CH_ALL): The sets have 21 SNPs in common: [1] "866_78" "1137_78"   "1910_79"   "7583_78"   "9415_78"   "10957_77"  "11235_78"  "13218_79"
[9] "14431_79"  "18719_79"  "23745_79"  "35231_79"  "39974_49"  "47695_78"  "72944_78" "80019_79"
[17] "85905_79"  "87783_79"  "89820_78"  "101352_79" "102737_78"

Comparing F, (SOU_M) and J, (CH_F): The sets have 1 SNPs in common:[1] "23745_79"

Comparing F, (SOU_M) and K, (CH_M): The sets have 28 SNPs in common: [1] "866_78"    "1137_78" "1910_79"   "4431_19"   "4885_78"   "5302_79"   "9415_78"   "9942_79"
[9] "10957_77"  "11235_78"  "13218_79"  "21844_79"  "23745_79"  "24957_79"  "25144_79" "31135_79"
[17] "35231_79"  "47695_78"  "52351_78"  "72944_78"  "75276_78"  "80019_79"  "87113_79" "87783_79"
[25] "89820_78"  "95709_79"  "101352_79" "102737_78"

####the most interesting follow from here

Comparing G, (IC-PE, ALL) and H, (IC-PE, F): The sets have 4 SNPs in common:[1] "10957_77" "23360_77" "73469_62" "99581_77"

Comparing G, (IC-PE, ALL) and I, (CH_ALL): The sets have 1 SNPs in common:[1] "10957_77"

Comparing G, (IC-PE, ALL) and J, (CH_F): The sets have 0 SNPs in common:character(0)

Comparing G, (IC-PE, ALL) and K, (CH_M): The sets have 3 SNPs in common:[1] "10957_77" "31135_79" "87113_79"

Comparing H, (IC-PE, F) and I, (CH_ALL): The sets have 10 SNPs in common: [1] "4975_78" "7583_78"  "8206_79"  "10957_77" "32700_77" "35262_76" "47695_78" "55665_78" "83973_79"
[10] "88895_78"

Comparing H, (IC-PE, F) and J, (CH_F): The sets have 0 SNPs in common:character(0)
Comparing H, (IC-PE, F) and K, (CH_M): The sets have 18 SNPs in common: [1] "4975_78" "10957_77" "14431_76" "24957_79" "25199_77" "31509_74" "32700_77" "34842_79" "35267_78"
[10] "45557_77" "47695_78" "75276_78" "83973_79" "84062_75" "87476_79" "88895_78" "96076_79" "97365_77"

Comparing I, (CH_ALL) and J, (CH_F): The sets have 7 SNPs in common:[1] "3995_79"  "8525_79" "20708_79" "23745_79" "28075_79" "39743_77" "46439_76"

Comparing I, (CH_ALL) and K, (CH_M): The sets have 78 SNPs in common: [1] "866_78" "1137_78" "1721_78" "1910_79" "4975_78" "5410_79" "7083_79" "7205_77"
[9] "7393_79" "8039_8" "8525_79" "9415_78" "9985_79" "10154_65" "10696_78" "10903_77"
[17] "10957_77" "11235_78" "11521_76" "11815_79" "12116_77" "13218_79" "15115_77" "18117_78"
[25] "19386_76" "20708_79" "23670_78" "23745_79" "23907_79" "28547_78" "29693_79" "32583_79"
[33] "32700_77" "35231_79" "36788_78" "43313_77" "45325_76" "45642_79" "46825_12" "47058_78"
[41] "47354_76" "47695_78" "48329_79" "49162_60" "49921_75" "53073_79" "57110_77" "57652_79"
[49] "62869_78" "68634_75" "70882_78" "72447_18" "72620_77" "72944_78" "74712_73" "75225_79"
[57] "77093_45" "78044_40" "80019_79" "80680_79" "81351_78" "81730_77" "82672_77" "83973_79"
[65] "87141_78" "87342_79" "87783_79" "88453_78" "88895_78" "89820_78" "91449_77" "96940_76"
[73] "98454_43" "98768_69" "100129_79" "101352_79" "102737_78" "108226_76"

Comparing J, (CH_F) and K, (CH_M): The sets have 3 SNPs in common:[1] "8525_79" "20708_79" "23745_79"
